# Molecular mechanism of the chitinolytic monocopper peroxygenase reaction

**DOI:** 10.1101/541292

**Authors:** Bastien Bissaro, Bennett Streit, Ingvild Isaksen, Vincent G.H. Eijsink, Gregg T. Beckham, Jennifer DuBois, Åsmund K. Røhr

**Affiliations:** Faculty of Chemistry, Biotechnology and Food Science, Norwegian University of Life Sciences (NMBU), P.O. Box 5003, N-1432 Aas, Norway; Department of Chemistry and Biochemistry, Montana State University, Bozeman, MT 59717-3400, USA; National Bioenergy Center, National Renewable Energy Laboratory, Golden, CO 80401, USA

**Keywords:** LPMO, CHITIN, CELLULOSE, BIOMASS, QM/MM, HYDROGEN PEROXIDE, PEROXYGENASE, MONOOXYGENASE, STOPPED-FLOW KINETICS

## Abstract

Lytic polysaccharide monooxygenases (LPMOs) are a recently discovered class of monocopper enzymes, broadly distributed across the Tree of Life. We recently reported that LPMOs can use H_2_O_2_ as an oxidant, revealing a novel reaction pathway. Here, we aimed to elucidate the H_2_O_2_-mediated reaction mechanism with experimental and computational approaches. *In silico* studies suggest that a network of hydrogen bonds, involving both the enzyme and the substrate, brings H_2_O_2_ into a strained reactive conformation, and guides the derived hydroxyl radical towards formation of a copper-oxyl intermediate. The initial H_2_O_2_ homolytic cleavage and subsequent hydrogen atom abstraction from chitin by the copper-oxyl intermediate are suggested to be the main energy barriers. Under single turnover conditions, stopped-flow fluorimetry demonstrates that LPMO-Cu(II) reduction to Cu(I) is a fast process compared to the re-oxidation reactions. We found that re-oxidation of LPMO-Cu(I) is 2000-fold faster with H_2_O_2_ than with O_2_, the latter being several orders of magnitude slower than rates reported for other monooxygenases. In agreement with the notion of ternary complex formation, when chitin is added, re-oxidation by H_2_O_2_ is accelerated whereas that by O_2_ slows. Simulations indicated that Glu60, a highly-conserved residue, gates the access to the confined active site and constrains H_2_O_2_ during catalysis, and Glu60 mutations significantly decreased the enzyme performance. By providing molecular and kinetic insights into the peroxygenase activity of chitinolytic LPMOs, this study will aid the development of applications of enzymatic and synthetic copper catalysis and contribute to understanding pathogenesis, notably chitinolytic plant defenses against fungi and insects.

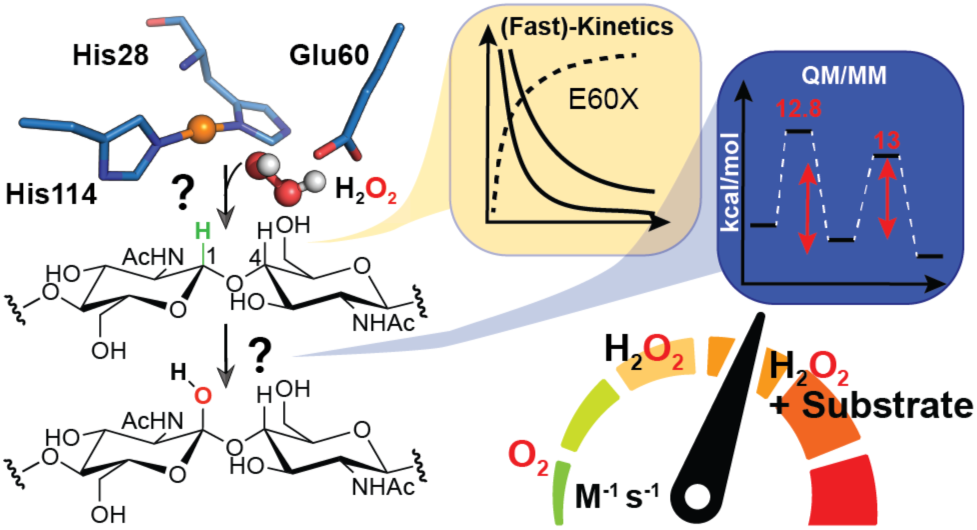

## INTRODUCTION

The landscape of bioinorganic chemistry has recently been enriched by the discovery that oxidative monocopper enzymes named lytic polysaccharide monooxygenases (LPMOs) can use H_2_O_2_ as a natural co-substrate,^1^ whereas O_2_ has long been invoked in all other established paradigms of copper-based enzymatic catalysis.^2^. The uniqueness of LPMOs lies mainly in the fact that they have the ability to catalyze the oxidative cleavage of glycosidic chains found in complex, recalcitrant environments such as the crystalline lattices formed by chitin,^3^ cellulose,^4–6^ or co-polymeric structures.^7,8^ LPMO action has been shown to disrupt the surface of recalcitrant polysaccharides,^9,10^ in turn enhancing the depolymerization catalyzed by canonical glycoside hydrolases.^3,11–13^ LPMOs are today classified as auxiliary activities (AA) in the Carbohydrate active enzyme database, in families AA9 to 11 and AA13 to 15. Given their biological abundance, notably in wood-decaying fungi,^14^ it is not surprising that LPMOs have rapidly become key elements of commercial enzymatic cocktails^15–17^ used in the biorefinery industry.^18^ Recent developments indicate that chitin-active LPMOs may play roles beyond biomass decomposition, such as acting as virulence factors in bacterial infections.^19–21^ On a similar note, a 2016 study showed that a “chitin-binding” protein (Tma12) is expressed by ferns as an insecticidal agent against whitefly.^22^ Importantly, although not recognized then as such, Tma12 resembles typical LPMOs: homology modeling shows a typical slightly distorted Fibronectin-like/Immunoglobulin-like β-sandwich core displaying the two surface-exposed, conserved histidines and phylogenetic analysis shows clustering with AA10 family members (**Figure S3**).

Despite substantial progress within the last decade, the mode of action of these monocopper enzymes is not yet fully mechanistically characterized.^16,23–25^ When acting on a carbohydrate substrate (R-H), and in the presence of excess reductant, a single oxygen atom derived from O_2_ is introduced into the final product,^3,26^ explaining why LPMOs have been widely recognized as monooxygenases.^26^ However, the monooxygenase reaction (R-H + O_2_ + 2e^-^ + 2H^+^ “ R-OH + H_2_O) requires a second electron (the first one being stored as Cu(I)) and protons during catalysis. These catalytic events remained highly puzzling considering (i) the monocopper, organic cofactor-free nature of LPMOs and (ii) that access of reductants to the active site is limited when bound to a polysaccharide substrate.^27^ Facing this question, Bissaro *et al.* demonstrated that LPMOs are capable of efficiently using H_2_O_2_ as a source of oxygen atoms in the hydroxylation reaction, thus making LPMOs a new kind of copper-dependent peroxygenase (R-H + H_2_O_2_ → R-OH + H_2_O; **Figure 1**).^1,28^ These findings suggest a straightforward answer to the “second electron conundrum”, since H_2_O_2_ brings the oxygen, electron, and proton equivalents necessary for a complete catalytic cycle. Such chemistry is new in the field of copper-enzymes and sparks interest in unraveling a new, H_2_O_2_-dependent mechanism. Although previous studies attempting to decipher the molecular details of the LPMO mechanism were carried out in a monooxygenase context, some conclusions drawn at that time are likely still valid but deserve to be re-investigated. Notably, a quantum mechanical (QM) study,^29^ followed by several other computational studies,^30,31^ suggested that the most likely species responsible for hydrogen atom abstraction (HAA) is a copper(II)-oxyl ([CuO]^+^) species. A similar downstream intermediate has been proposed for the H_2_O_2_ reaction mechanism,^1^ where [CuO]^+^ is obtained after reaction of H_2_O_2_ with the LPMO-Cu(I) state, concomitant with H_2_O release.

**Figure 1.**
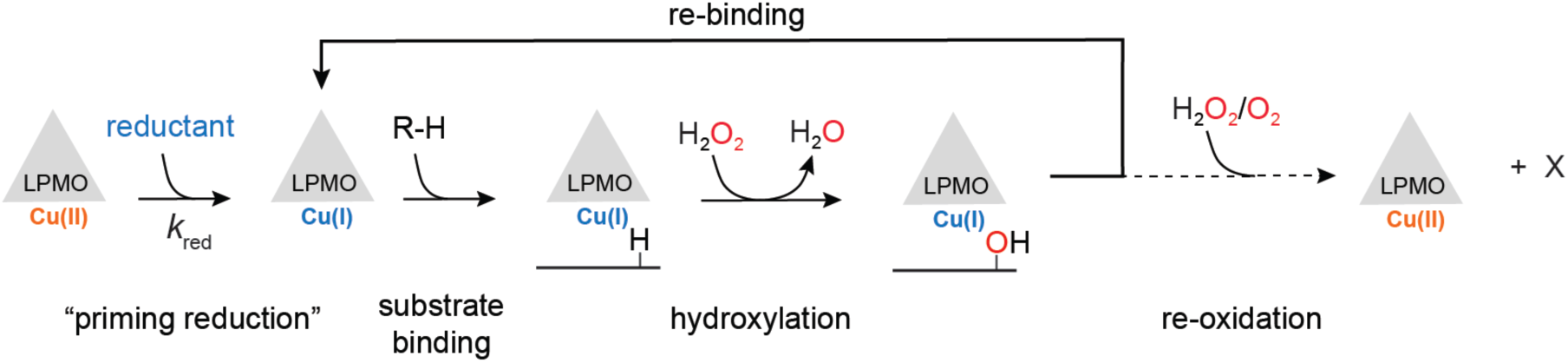
Simplified reaction scheme of *Sm*AA10A-catalyzed peroxygenation of chitin. Reduction of the ground state LPMO-Cu(II) to LPMO-Cu(I), which binds with greater affinity to substrate,^36,37^ is thought to precede substrate binding leading to a confined active site^27^ ready to react with H_2_O_2_, resulting in hydroxylation of the substrate and release of a water molecule.^1^ The hydroxylated product (either at the C1 or C4 carbon) is unstable and undergoes an elimination reaction inducing glycosidic bond cleavage.^26,32^ At the end of the catalytic cycle, the Cu(I) state is conserved, and the enzyme could potentially enter a new catalytic cycle or be oxidatively inactivates via non-productive reactions with oxidants. X denotes the reaction products (e.g. hydroxyl radicals, superoxide) originating from such de-activation reactions.

Wang et al.^32^ recently published a computational study on an C4-oxidizing AA9 in complex with a water soluble cellotriose molecule^33^ that is line with the proposed peroxygenase mechanism.^1^ However, no supporting biochemical data on H_2_O_2_ reactivity have been reported for that AA9. Subsequent to the discovery of the role of H_2_O_2_ in LPMO catalysis, a study by Kuusk *et al.* determined the steady-state kinetic parameters of H_2_O_2_-driven chitin oxidation by the C1-oxidizing LPMO from the soil bacterium *Serratia marscecens* (*Sm*AA10A, also known as CBP21).^34^ This study revealed a catalytic constant (*k*cat) of 6.7 s^-1^ and an apparent *K*_M_ for H_2_O_2_ in the low micromolar range (ca. 3 µM) resulting in a *k*_cat_/*K*_M_ ~ 10^6^ M^-1^s^-1^. These constants are consistent with values reported for well-characterized heme-dependent peroxygenases.^35^

Building on biochemical and steady-state kinetics evidence gathered for *Sm*AA10A,^1,34^ and also on experimentally-supported molecular models of *Sm*AA10A on crystalline chitin complexes,^27^ we aimed to decipher the molecular details of the newly proposed H_2_O_2_ reaction mechanism. Employing several computational methods, we calculated the energy barriers associated with hypothesized reaction coordinates and H_2_O_2_ diffusion pathways to the active site. As experimental support, we used pre-steady state kinetic methods to examine *Sm*AA10A reduction and single turnover reactions of *Sm*AA10A, isolated in the Cu(I) form, with both H_2_O_2_ and O_2_. QM/MM calculations suggested a key role in catalysis for Glu60, a highly conserved second-shell residue, which was subsequently interrogated experimentally by site directed mutagenesis. Collectively, these results provide a strong mechanistic and thermochemical foundation for describing chitinolytic LPMOs as unusual biological peroxygenases functioning through a [CuO]^+^ intermediate.

## EXPERIMENTAL AND COMPUTATIONAL DETAILS SECTION

A complete description of the experimental details is available in the **Supporting information** document, including details on materials, site-directed mutagenesis, production and purification of recombinant LPMOs, LPMO activity and binding assays, (stopped-flow) fluorimetry experiments and computational studies.

## RESULTS

### H_2_O_2_ is the co-substrate sustaining SmAA10A catalysis under steady-state turnover conditions

H_2_O_2_ has previously been shown to serve as an efficient co-substrate of *Sm*AA10A-catalyzed oxidation of chitin.^34^ However, the question remains whether *Sm*AA10A catalysis also relies on H_2_O_2_ in conditions where H_2_O_2_ is not added, i.e. just in presence of ambient O_2_ concentrations and excess reductant. The latter conditions are standard practice in the LPMOs field. We argued previously that they promote the *in situ* formation of H_2_O_2_, and therefore that most reported LPMO rates reflected a peroxygenase rather than a putative monooxygenase activity. Here, we first carried out a competition experiment in presence of a peroxidase (**Figure 2A**). This experiment shows that, in presence of ambient O_2_ (no added H_2_O_2_) and 1 mM reductant (ascorbic acid, AscA), 1 μM *Sm*AA10A can be almost totally inhibited by the H_2_O_2_-scavenging activity of HRP, suggesting that H_2_O_2_ is produced *in situ* and is the main (if not only) oxidant used by *Sm*AA10A. In agreement with the proposal of the formation of a ternary complex founded on the basis of steady-state kinetics studies,^34^ we show, after determining suitable reaction conditions (i.e. 50 nM LPMO and 50 μM EDTA to diminish non-enzymatic background reactions; see **Figure S6**), that fast and efficient H_2_O_2_ consumption only occurs in the presence of both *Sm*AA10A and β-chitin (**Figure 2B**).

**Figure 2.**
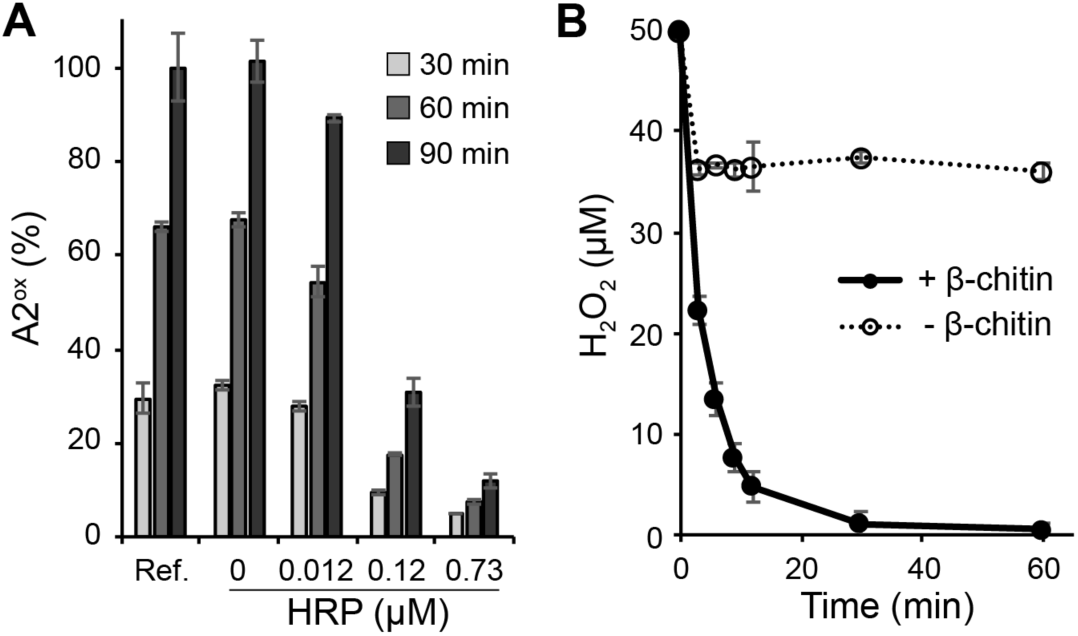
Assessing the underlying role of H_2_O_2_ in *Sm*AA10A-catalyzed oxidation of chitin. (A) The graph shows the time-course release of soluble oxidized products (chitobionic acid; A2^ox^) from β-chitin (10 mg·mL^-1^) by *Sm*AA10A (1 µM) in presence of AscA (1 mM) and O_2_ (ca. 200 µM at atmospheric pressure) with different amounts of horseradish peroxidase (HRP) and AmplexRed (200 µM, a co-substrate of HRP). Quantities are expressed as a percentage of the amount of A2^ox^ detected in the reference reaction (“Ref.”; i.e without HRP and AmplexRed) after 90 min reaction. **(B)** Time-course consumption of H_2_O_2_ (50 µM at t_0_) by *Sm*AA10A (50 nM) in a reaction containing AscA (20 µM) and EDTA (50 µM; see **Figure S6** for details) in presence and absence of β-chitin (10 mg·mL^-1^). Note that the fast drop observed for instance in the “no chitin” experiment is due to some experimental limitations (**Figure S7**). All reactions were carried out in sodium phosphate buffer (50 mM, pH 7.0) at 40 °C in a thermomixer (1,000 rpm). Error bars show ± s.d. (n = 3, independent experiments).

### QM/MM study of the chitinolytic peroxygenase reaction

Intriguingly, neither strict C1-oxidizing LPMOs, nor crystalline polysaccharide substrates, have been the subject of QM/MM-assisted investigations of the LPMO mechanism.^29^,^31,32,38–40^ Also, in the few available reports,^32,40^ the initial steps of H_2_O_2_ access, positioning, and one electron reduction by Cu(I) at the enzyme active site have not been described and have not been agreed upon (*vide infra*). Starting from a previously published, biochemically and spectroscopically supported model of *Sm*AA10A in complex with crystalline β-chitin (**Figures 3A and S4**),^27^ we performed QM/MD and QM/MM calculations to unravel the molecular details underlying the peroxygenase reaction (R-H + H_2_O_2_ → R-OH + H_2_O) catalyzed by *Sm*AA10A-Cu(I).

**Figure 3.**
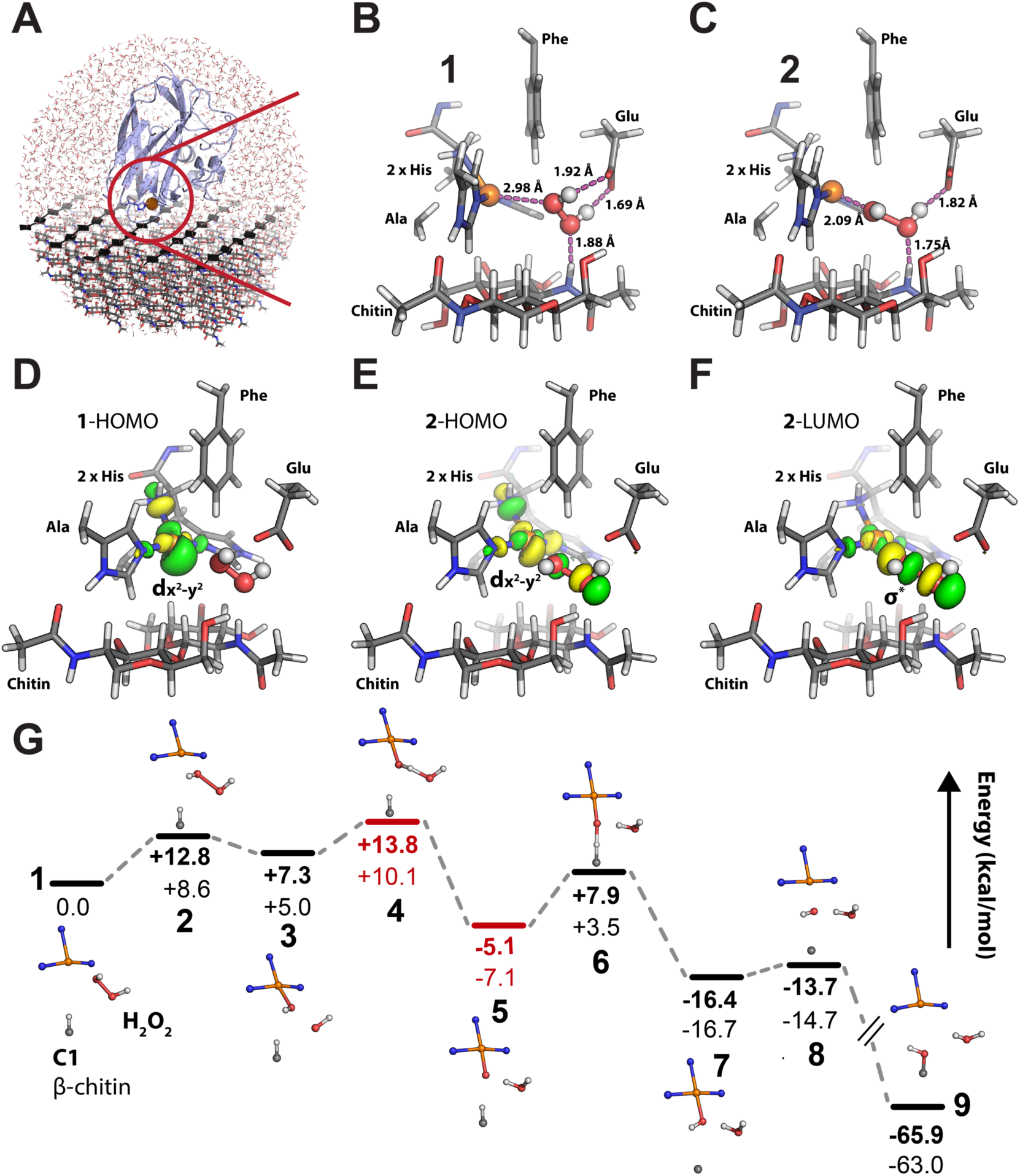
QM/MM study of the peroxygenase mechanism of *Sm*AA10A on crystalline β-chitin. **(A)** The QM/MM model containing ~24,000 atoms derived from experimentally-supported models of *Sm*AA10A in complex with chitin.^27^ See **Figure S5** for a definition of the QM region. **(B)** In the initial state of the reaction (**1**), the H_2_O_2_ molecule is aligned toward the Cu(I) ion in a strained conformation by both substrate and amino acid side chain hydrogen bonds. The first transition state (**2**) displays an elongated H_2_O_2_ O-O bond and the reactive hydroxyl radical to be formed is stabilized by H-bonds from Glu60 and a substrate amide H-atom. **(C)** The electron transfer from Cu(I) to H_2_O_2_ can be visualized by plotting the quasi-restricted orbitals of **1** and **2**. Before reaction with H_2_O_2_, the state **1** HOMO exhibits delocalized d_x2-y2_ character, indicating a Cu(I) state ready to react with H_2_O_2_. The state **2** (“**2**-HOMO” to “**2**-LUMO” transition) indicates that the Cu(I) electron moves from the delocalized d_x2-y2_ like orbital to a σ* anti-bonding orbital localized between the H_2_O_2_ O-atoms, promoting bond cleavage. **(D)** Complete reaction path where states **1**-**9** indicate energy minima and transition states (relative energies in kcal mol^-1^) (see **Figure S8** for snapshots of each states). Between states **3** and **6**, spin crossovers occur, and the triplet states are shown in red and the open shell singlet states are indicated in black. Numbers in bold are obtained by the B3LYP functional while the numbers immediately below were obtained by TPSSh. All protein residues and chitin (only a dimer is shown in panels B and C) are displayed as grey sticks. H_2_O_2_ is shown as white and red sticks-and-balls. For the sake of clarity, the MM-region is not shown. HOMO, highest occupied molecular orbital; LUMO, lowest unoccupied molecular orbital.

QM/MM/MD simulations (**Figure S5**, **Movie S1**), followed by NEB minimum energy path calculations resulted in starting models for calculating the energies of the different states (**1**-**9**) of the *Sm*AA10A-catalyzed reaction (**Figure 3D**). A global observation is that the overall reaction is thermodynamically favorable, with a relative energy difference of −65.9 kcal mol^-1^ between the initial and final states. State **1** represents the *Sm*AA10A-Cu(I) bound to β-chitin substrate with a H_2_O_2_ molecule confined in the reaction cavity and no water molecules coordinated to the copper ion. Hydrogen bonds from the Glu60 side chain and a substrate amide hydrogen align the H_2_O_2_ molecule towards the Cu(I) ion at a distance of 2.98 Å (**Figure 3B**). The H_2_O_2_ O-O bond length of **1** is 1.44 Å, which is equal to the value obtained by geometry optimization of H_2_O_2_ in vacuum (1.44 Å). Interestingly, the dihedral bond angle of H_2_O_2_ in **1** is 53°, far from the equilibrium angle (113.6° in gaseous state,^41^ 90.2° in solid state^42^), indicating that the protein and substrate introduce a strained conformation (+ 4.1 kcal mol^-1^) of the peroxide moiety. The *Sm*AA10A catalytic cycle is triggered by electron transfer from Cu(I) to H_2_O_2_, and the transition state of this step is represented by **2** (**Figure 3C**). The H_2_O_2_ bond is now elongated (1.76 Å), the dihedral bond angle has relaxed to 94°, and the proximal peroxide O-atom is only 2.1 Å away from the copper ion. The energy barrier of this step is estimated to be 12.8 kcal mol^-1^ and 8.6 kcal mol^-1^ by the functionals B3LYP and TPSSh, respectively. State **2** is best described as an open-shell singlet, but reaching that conclusion required detailed analysis of the electronic structure (see **Supplementary Results**). We plotted the quasi-restricted orbitals that best describe the transition of state **1** to **2** in **Figure 3C**. The highest occupied molecular orbital (HOMO) of state **1** has mainly Cu-d_x2-y2_ and ligand π character (**Figure 3C**, “**1**-HOMO”), and the largest lobe points directly towards the H_2_O_2_ molecule. Reaching the transition state geometry (state **2**), the HOMO orbital is also slightly delocalized to include the H_2_O_2_ molecule (**Figure 3C**, “**2**-HOMO”). According to frontier molecular orbital theory, the lowest unoccupied molecular orbital (LUMO) indicates where the Cu(I) electron will go. Here, the state **2** LUMO displays a decreased density at the copper d-orbital, while a σ^*^ anti-bonding orbital appears between the H_2_O_2_ O-atoms (**Figure 3C**, “**2**-LUMO”), rationalizing the homolytic cleavage of the O-O bond. The first local minimum in the reaction path is state **3** (**Figure 3D and S8**), where the calculations suggest a Cu-bound hydroxide with an adjacent hydroxyl radical. This state also collapses to an open shell singlet solution due to strong antiferromagnetic coupling between the Cu(II) and hydroxyl radical (See **Supplementary Results** for more details). Hydrogen bonds from both the substrate and the protein appear to confine and orient the reactive hydroxyl radical in the reaction cavity, yielding a “precision-guided HO•” poised for hydrogen atom abstraction from the Cu-bound hydroxide with a barrier of 6.5 kcal mol^-1^ (B3LYP; state **4**, **Figure S8**). At this point, the total spin of the system has changed from a singlet to a triplet state displaying spin crossover behavior. The resulting state **5** (**Figure 3D and S8**) is best described as a triplet copper-oxyl [CuO]^+^ species, where the oxyl-O atom, now only 2.08 Å away from the substrate H-atom that is to be abstracted, is hydrogen bonded to the water molecule formed in the previous reaction step. In this highly reactive state, a Löwdin spin population analysis reveals Cu and oxyl spins of 0.61 and 1.1, respectively (see **Table S2** for all values along the reaction path).

The energy barrier of the substrate H-abstraction step (state **6**, **Figures 3D and S8**) depends on the functional utilized in the calculations. The B3LYP and TPSSh functionals yield values of 13.0 and 10.6 kcal mol^-1^, respectively. At this point of the reaction, the system has changed back to the singlet energy surface indicating that the spin crossover transitions take place between state **3** and **6** in the reaction path. The hydroxide moiety formed by the H-abstraction step partly recombines with the copper ion (Cu-O distance of 1.91 Å) in a local energy minimum (state **7**, **Figures 3D and S8**). Finally, the Cu-interacting hydroxide passes through the low barrier (+2.6 kcal mol^-1^ for B3LYP) rebound-intermediate (state **8**, **Figure 3D and S8**), and recombines with the substrate radical to form the final hydroxylated product (state **9**, **Figure 3D and S8**). Note that the Cu(I) form of the enzyme has been regenerated at the end of the reaction cycle. Importantly, it appears that along the reaction path, hydrogen bonds from the enzyme Glu60 side chain and a strategically positioned substrate amide hydrogen control the positioning of several H_2_O_2_-derived reactive oxygen species (**Figure S8**), facilitating the LPMO reaction.

We also investigated an alternative mechanism, inspired by a previous study,^29^ where a Cu(I) activated O_2_ performs the substrate H-abstraction at the triplet energy surface (**Figure S9)**. The main conclusions are that substrate binding alters the orientation of the copper-bound superoxide, and that H-abstraction by a copper superoxide intermediate is thermodynamically plausible but kinetically very unlikely (energy barrier of 35 kcal mol^-1^) (see **Supplementary Results**).

### The reductive priming reaction is fast and monophasic

To gain insights into the priming reduction, i.e. the ability of an LPMO to catalyze the peroxygenase reaction once primed by sub-stoichiometric amounts of reductant,^1,37^ we investigated how fast such reductive reaction occurs and which amount of reductant is required to achieve complete LPMO reduction. We have previously shown^43^ that the fluorescence signal of an LPMO can be used as a reporter of the redox state of the LPMO (**Figure S10**), the Cu(II) and Cu(I) forms having respectively low and high fluorescence responses, both linear within the concentration range studied (**Figure S11**). Following rapid stopped-flow mixing of *Sm*AA10A-Cu(II) with 1 eq. AscA (5 μM each) under anaerobic conditions, a monophasic increase in fluorescence was observed. The experiment was subsequently repeated with AscA present in pseudo-first order excess of the *Sm*AA10A-Cu(II) (≥ 10 eq). The total fluorescence change remained the same, suggesting that reduction was complete with as little as 1 eq. of AscA. The reduction reaction remained monophasic but increased in rate with increasing concentrations of AscA (**Figure 4A**). Each fluorescence versus time curve could be fit with a single exponential function, indicative of a pseudo-first order rate reaction with respect to AscA. The pseudo-first order rate constant (*k*obs) varied linearly with [AscA], yielding a second order rate constant *k*AscA = 4.2 x 10^5^ M^-1^ s^-1^ (**Figure 4B**).

**Figure 4.**
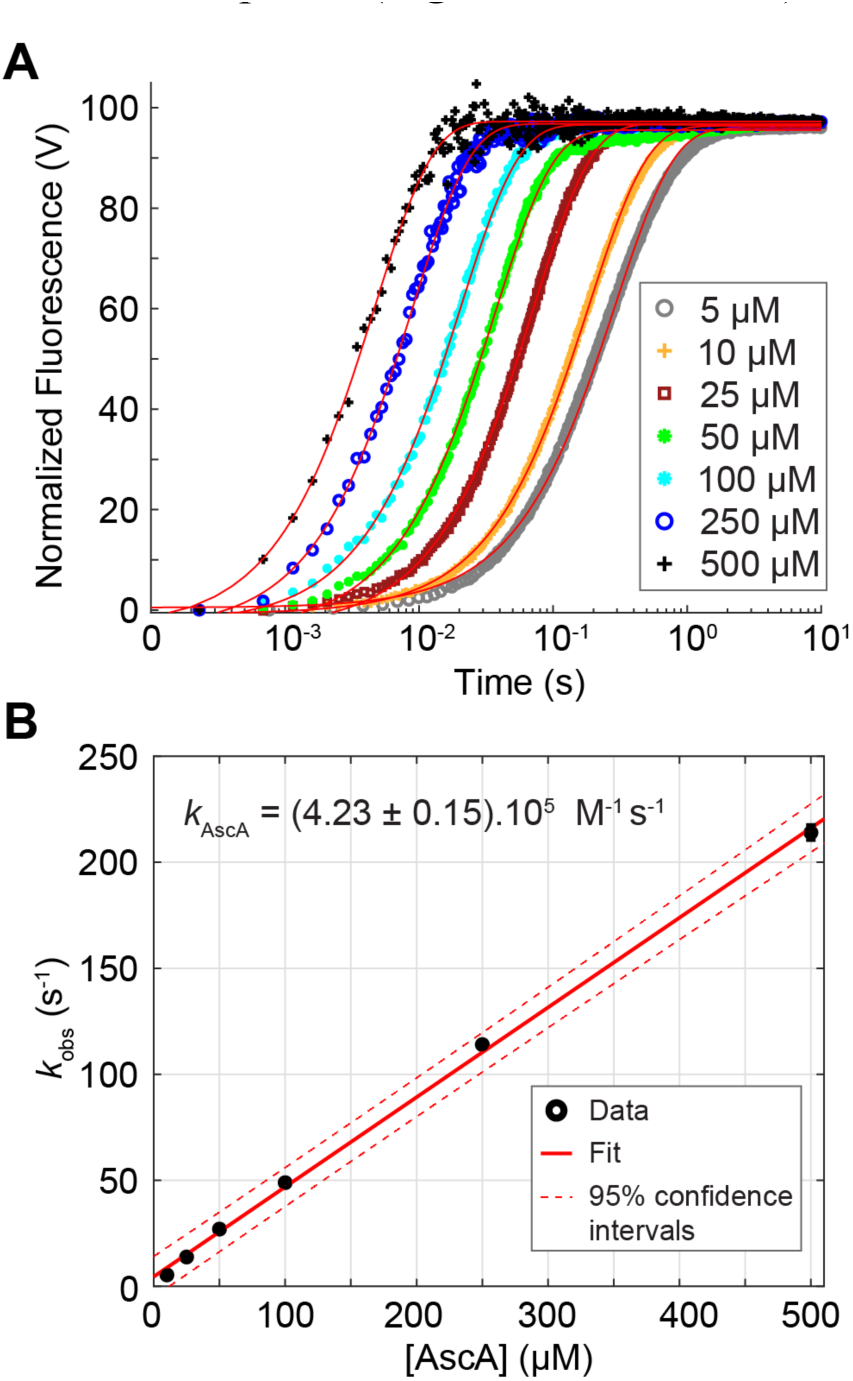
Kinetics and ascorbate concentration dependence of the reduction of *Sm*AA10A-Cu(II) to Cu(I). **(A)** *Sm*AA10A-Cu(II) (final concentration, 5 µM) was anaerobically mixed with varying concentrations of AscA and the changes in fluorescence associated with reduction of the Cu(II) monitored as a function of time. The reactions were carried out in potassium phosphate buffer (50 mM, pH 7.1) at 25 °C. The final concentrations of AscA are indicated in the figure. Data were fit with single exponential functions (red lines) to give observed rate constants (*k*_obs_) at each substrate concentration. Each experiment was performed in triplicate. For the sake of clarity, only the trace of one replicate is shown for each condition. **(B)** Plot of pseudo-first order *k*_obs_ as a function of AscA concentration. Error bars show ± s.d. (n = 3, independent experiments).

We showed (*vide infra*) that hexa-*N*-acetyl-chitohexaose (NAG_6_) is a suitable substrate for single turnover experiments. Here, we show that in the presence of 1 mM NAG_6_ a second order rate constant of 5.4 x 10^5^ M^-1^ s^-1^ is obtained for the reduction of *Sm*AA10A-Cu(II) by AscA (**Figure S12**). This value is 1.26-fold higher than that obtained in absence of NAG_6_, suggesting that the presence of a bound substrate does not strongly modulate the rate of reaction of the *Sm*AA10A-Cu(II) by AscA.

### The reaction between Sm*AA10A-Cu(I) and O_2_ is exceedingly slow*

To further understand the hypothetical competition between environmentally available H_2_O_2_ and ambient O_2_, we first monitored the single turnover reaction of 2 μM *Sm*AA10A-Cu(I) (produced anaerobically by reduction of the oxidized enzyme with ascorbate, followed by removal of excess reductant) and O_2_ over time (**Figure 5A**). The change in the fluorescence spectrum following mixing with 1 eq. O_2_ was not possible to discriminate from the background noise (**not shown**). Using pseudo-first order concentrations of O_2_ ([O_2_] at least a 10-fold excess relative to [*Sm*AA10A-Cu(I)]), the reaction led to re-oxidation of the Cu(I) to Cu(II) over a period of minutes, to an extent that was dependent on [O_2_]. When fitting the kinetic data, a number of models were explored (**Figures S13 and S14**).

Although we have based our analysis on a simple model, a single exponential function combined with a linear function **(Figure S13&14B)**, the residuals of the fits indicate that the re-oxidation of *Sm*AA10A-Cu(I) by O_2_ is a complex reaction. Indeed, a complex reaction model where O_2_ binds and is reduced by a binuclear copper cluster formed by a *Sm*AA10A-Cu(I) dimer results in a perfect fit **(Figure S15, scheme S1)**. In such an arrangement, the *Sm*AA10A-Cu(I) dimer binuclear active site would resemble well-studied type 3 copper enzymes such as tyrosinases and hemocyanins.^2^ This dimer-hypothesis is currently being explored in a separate study. Nonetheless, applying the single exponential model as best simple approximation shows that the *k*obs varied linearly with [O_2_], yielding a second order rate constant *k*O2 = 3.3 M^-1^ s^-1^ (**Figure 5B**).

**Figure 5.**
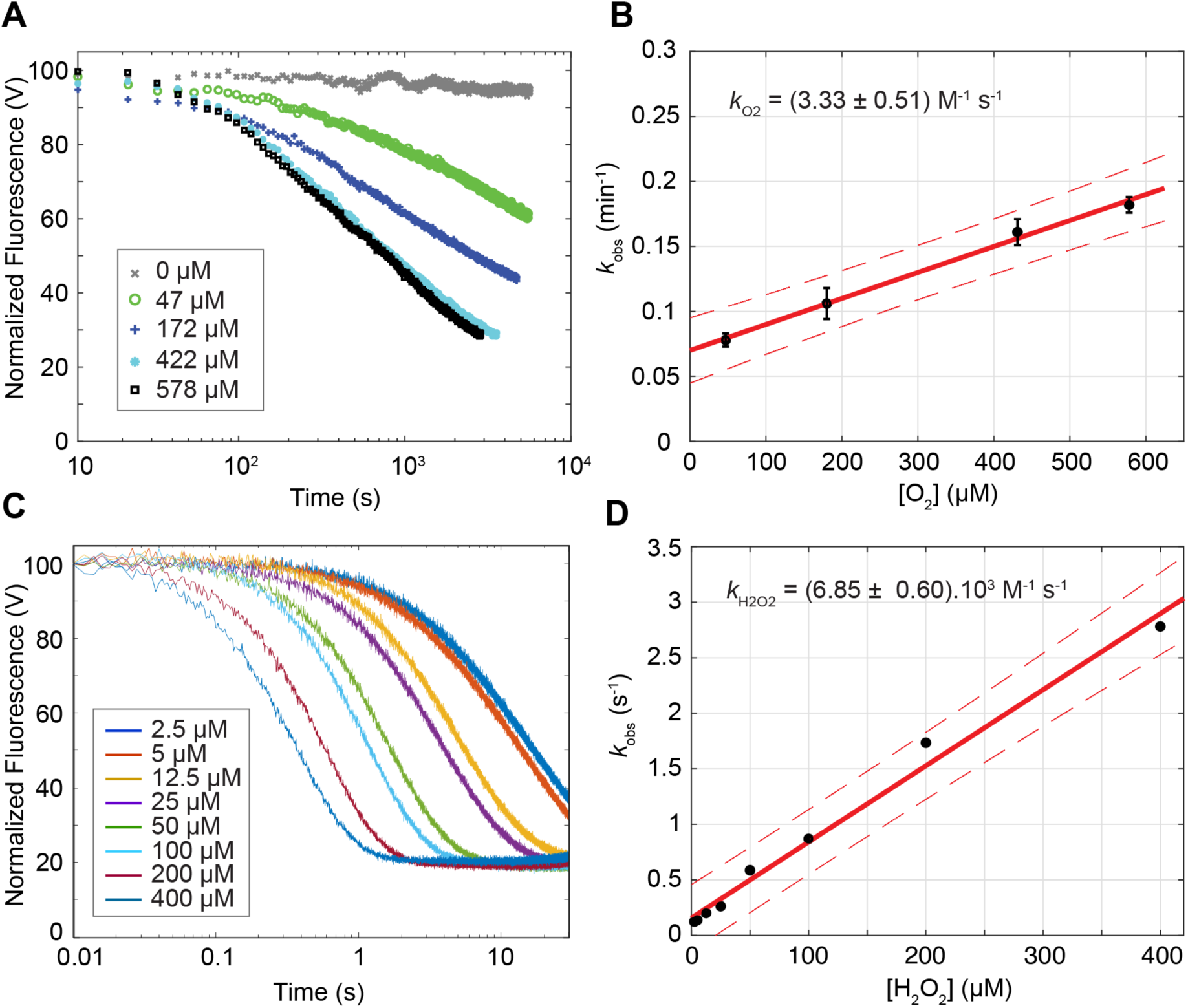
Single turnover re-oxidation of *Sm*AA10A-Cu(I) by (A-B) O_2_ or (C-D) H_2_O_2_ in solution. *Sm*AA10A-Cu(I) (final concentration 2 µM) was anaerobically mixed with varying concentrations of **(A)** O_2_ or **(C)** H_2_O_2_ (see color code in the figure) and the changes in fluorescence associated with re-oxidation of the Cu(I) monitored as a function of time. The reactions were carried out in sodium phosphate buffer (50 mM, pH 7.0) at 25 °C. Data were fit with single exponential functions (see **Figures S13&14-B and S16**) to give observed rate constants (*k*_obs_) at each co-substrate concentration and plotted versus the latter to obtain second order rate constants for **(B)** O_2_ or **(D)** H_2_O_2_. In panels A and C, for the sake of clarity, only one trace for each condition is shown (experiments were performed at least in triplicate). In panels B and D, error bars show ± s.d. (n = 3, independent experiments), red solid lines show the best linear fit and the red dotted line the 95% confidence interval.

### The reaction between Sm*AA10A-Cu(I) and H_2_O_2_ is monophasic and faster than the reaction with O_2_*

Monitoring the single turnover reaction of anaerobic/reduced *Sm*AA10A-Cu(I) (2 μM) with varying concentrations of H_2_O_2_ showed that the enzyme underwent a change in fluorescence consistent with complete re-oxidation of the copper within seconds to milliseconds (**Figure 5C**). The data can be described by single exponential functions (**Figure S16**), indicative of a pseudo-first order reaction with respect to H_2_O_2_. A stoichiometry of 1:1 between *Sm*AA10A-Cu(I) and H_2_O_2_ was verified (**Figure S17**). The fluorescence signature, together with spin trap experiments (**Figure S18**), indicate that the products of this reaction are *Sm*AA10A-Cu(II) and hydroxyl radicals (80 % OH• yield). Plots of the fitted pseudo-first order rate constants *k*obs as a function of [H_2_O_2_] could be fit to a straight line, yielding the second order rate constant *k*H2O2 = 6.9 x 10^3^ M^-1^ s^-1^ (**Figure 5D**). Hence, for comparable concentrations of oxidant, the time required to convert Cu(I) to Cu(II) is approximately three orders of magnitude faster with H_2_O_2_ than with O_2_.

Furthermore, we explored the catalytic potential of *Sm*AA10A-Cu(I) reacted with H_2_O_2_ in solution *prior* to addition of NAG_6_ substrate. If a stable reactive enzyme-oxygen intermediate formed in solution, we would expect to observe amounts of oxidized products proportional to the intermediate life-time when adding substrate after different incubation times with H_2_O_2_. We observed a time-dependent decay in the amount of oxidized products formed (**Figure S19**) that corresponds to the predicted decay in amounts of available reactants (i.e. *Sm*AA10A-Cu(I) and H_2_O_2_ remaining after a time-dependent fraction of *Sm*AA10A-Cu(I) has been consumed by the reaction with H_2_O_2_ in absence of substrate calculated using *k*H2O2 = 6.9 × 10^3^ M^-1^ s^-1^). This indicates that if a productive intermediate resulting from the reaction of *Sm*AA10A-Cu(I) with H_2_O_2_ would be formed in solution, then its lifetime is much shorter than the time required for a catalytic turnover in presence of substrate. Also, we show that reacting free *Sm*AA10A-Cu(I) with H_2_O_2_ yields only a very low loss in activity (**Figure S20**). This indicates that under single turnover conditions we monitor re-oxidation of *Sm*AA10A-Cu(I) to Cu(II) (see **Supplementary Discussion**) rather than LPMO inactivation, which thus appears to require multiple turnovers.

### The substrate accelerates the reaction of Sm*AA10A-Cu(I) with H_2_O_2_, but not with O_2_*

QM/MM calculations predict that several hydrogen bonds are established between H_2_O_2_, the LPMO, and the substrate during H_2_O_2_ activation (**Figure 3B**), and biochemical data indicate that H_2_O_2_ is productively turned-over only in presence of chitin (**Figure 2B**). Following the same procedure outlined above, we can experimentally observe the impact of substrate on the first re-oxidation event (corresponding to steps 1→3; **Figure 3C**). For this, we used (NAG)_6_, which is compatible with fluorimetry experiments. Although NAG_6_ is a low affinity substrate under steady-state conditions (see below), we have previously shown^27^ that NAG_6_ constitutes a good proxy of β-chitin in terms of molecular interaction with the enzyme. Most importantly, under single turnover conditions, *Sm*AA10A-Cu(I) fully engaged in NAG_6_ oxidation rather than non-productive deactivation pathways (**Figure S21**). However, we also note that the reduced LPMO can hardly catalyze more than one turnover with NAG_6_ as a substrate. In absence of excess of reductant, more than one turnover would require the Cu(I) form of the enzyme to be available at the end of the first catalytic cycle for the following reactions. We see that the Cu(I) species is apparently regenerated in presence of NAG_6_ only when low, sub-stoichiometric H_2_O_2_ concentrations are employed (**Figure S17**). This result indicates that at higher, supra-stoichiometric H_2_O_2_ concentrations the regenerated Cu(I) species will eventually be re-oxidized (as observed below; see **Supplementary discussion**). On the basis of energy profiles (**Figure 3C)**, the first re-oxidation event is likely to determine the overall reaction rate, including the step of final re-oxidation to Cu(II) state observed by fluorescence. **Figure 6A** shows that in presence of 0.5 mM NAG_6_ re-oxidation of *Sm*AA10A-Cu(I) by O_2_ is slightly slower. Contrarily, using 0.5 mM NAG_6_, re-oxidation of *Sm*AA10A-Cu(I) by H_2_O_2_ was faster and displayed monophasic time-course profiles similar to experiments without substrate (**Figure S22**). The experiment was repeated with higher concentrations of NAG_6_ showing a proportional increase in re-oxidation rates by H_2_O_2_, with second order rate constant up to *k*H2O2 = 21 x 10^3^ M^-1^ s^-1^ (**Figure 6B**). In a complementary spin-trapping experiment, we also showed that less hydroxyl radicals could be detected in presence of NAG_6_ (**Figure S18D**).

**Figure 6.**
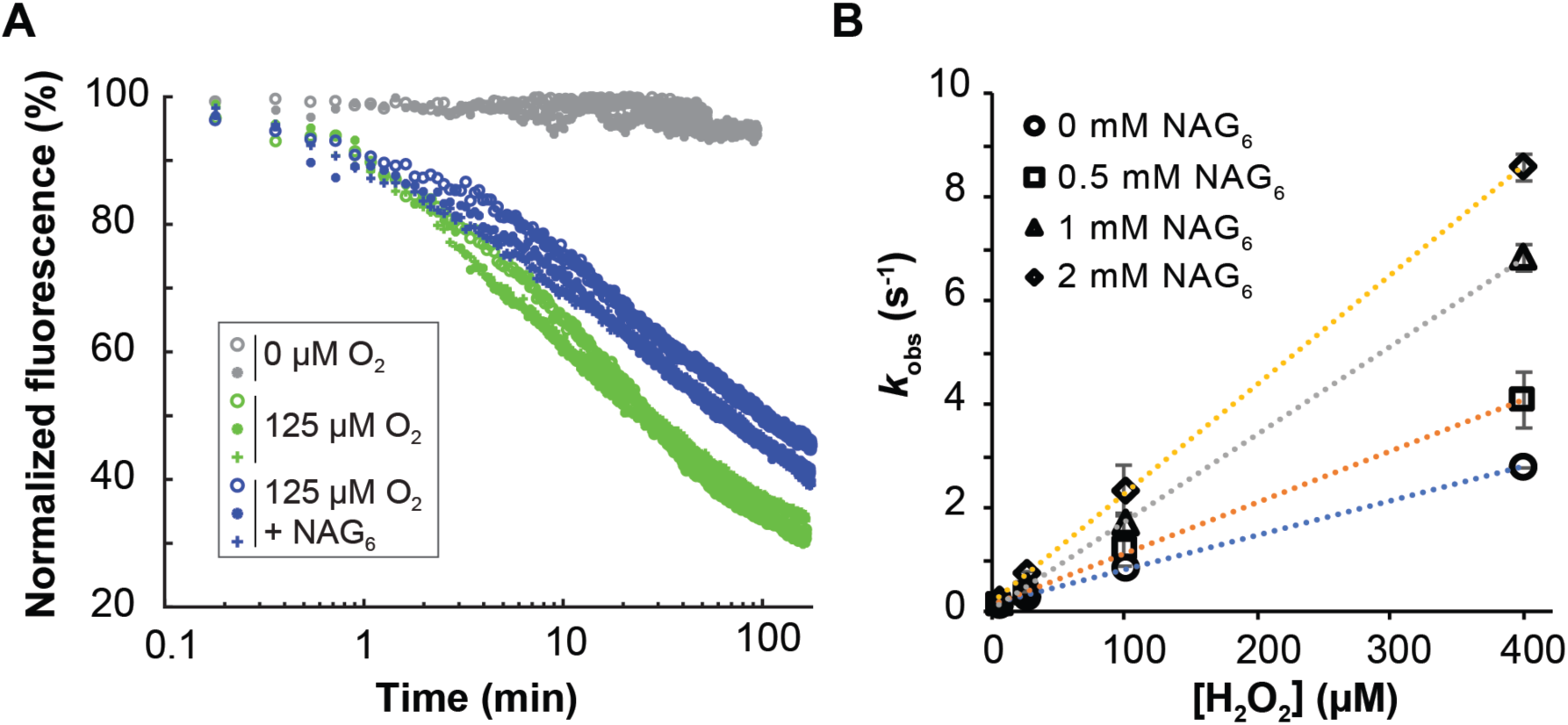
Re-oxidation of *Sm*AA10A-Cu(I) by (A) O_2_ and (B) H_2_O_2_ in presence of NAG_6_. (A) Time-curves of re-oxidation of *Sm*AA10A-Cu(I) (2 µM) by 125 µM O_2_ in presence (0.5 mM) and absence of NAG_6_. All given concentrations are concentrations after mixing. **(B)** Second order rate plot of *Sm*AA10A-Cu(I) re-oxidation by H_2_O_2_ in presence (0.5 to 2 mM) and absence of NAG_6_ (see **Figure S22** for example of time curves obtained with 0.5 mM NAG_6_). Panel A shows the traces obtained for each replicate (n = 3) and error bars in panel B show ± s.d. (n = 3, independent experiments).

### H_2_O_2_ access to the active site

We have previously revealed the existence of a water tunnel, formed in the enzyme-polysaccharide complex interspace, connecting bulk solvent to the monocopper active site.^27^ Here, we investigated the energetics associated with H_2_O_2_ diffusion into the active site by MD simulations combined with umbrella sampling, estimating free energy profiles of H_2_O_2_ moving in this interspace (See **Supplementary Results** and **Figures S1, S23** and **movie S2)**. The free energy barrier associated with diffusion of H_2_O_2_ into the reaction cavity from the bulk solvent was estimated to be less than 2 kcal/mol. Also, during the course of simulations, Glu60 and Asn185 appear to play a gating role, restricting H_2_O_2_ access to the active site cavity. In spite of the small energetic cost associated with H_2_O_2_ entering the confined reaction cavity, such confinement may offer greater control on reagents involved in catalysis. In agreement, we show that under steady-state conditions *Sm*AA10A is inefficient at consuming H_2_O_2_ when β-chitin is replaced by the soluble substrate NAG_6_ (**Figure S2**).

### Glu60 is important for efficient peroxygenase activity

QM/MM and MD calculations presented above suggest that Glu60 is involved in gating access to the active site and also constrains H_2_O_2_ positioning for reacting with Cu(I) (**Figure 3B**). This residue is structurally conserved throughout the entire LPMO superfamily, either as a glutamate or as a glutamine.^44^ Therefore, to probe and complement our simulations, we investigated the role of Glu60 in *Sm*AA10A catalysis via site-directed mutagenesis. We first investigated the impact of Glu60 mutation on the apparent oxidative activity on β-chitin (**Figure 7A**). **Figure 7A** shows that mutations of Glu60 lead to a drastic decrease in product yield after 24 h of reaction compared to the WT enzyme (83 to 97% lower). However, in terms of initial rates (see **Figure 7A inset**), the impact of Glu60 mutations is less dramatic but still significant since mutants E60D, N, Q and S show 64, 28, 62, and 48% of apparent activity relative to the wild-type enzyme, respectively. Therefore, in conditions where H_2_O_2_ is produced gradually *in situ* (e.g. **Figure 2A**), Glu60 appears important but not crucial for activity in the initial phase. Gradual inactivation of the LPMO, as shown by very low final yields (**Figure 7A**), suggests that the mutants are less efficient in consuming H_2_O_2_ produced *in situ*, which results in H_2_O_2_ accumulation that is known to be detrimental for LPMO stability in reducing conditions. ^1^,^34,45,46^ This can potentially be related to poorer binding to β-chitin ^45^ and/or, conversely, to intrinsically less efficient turnover of H_2_O_2_. We show that Glu60 mutants retained between 43 and 75% of the WT binding capacity in the early phase of the process (i.e. after 15 min of incubation) and between 60 and 82% at a later stage (i.e. after 4 h incubation) (**Figure 7B and S24)**. In comparison, the ability of Glu60 mutants to consume H_2_O_2_ was much more affected (**Figure 7C**). Also, in absence of substrate, and under single turnover conditions, the reactivity of the free reduced Glu60 mutants with H_2_O_2_ was equivalent or slightly lower than that of the WT (**Figure S25 and Table S5**). Therefore, considering that reactivity of H_2_O_2_ leading to productive catalysis is three orders of magnitude faster (10^6^ M^-1^ s^-1^) than the one leading to inactivation (ca. 10^3^ M^-1^ s^-1^),^34^ the most likely scenario is that Glu60 mutations impair in first place the ability of substrate-bound LPMOs to turnover efficiently H_2_O_2_, which in turn leads to an accumulation of H_2_O_2_ inducing inactivation.

**Figure 7.**
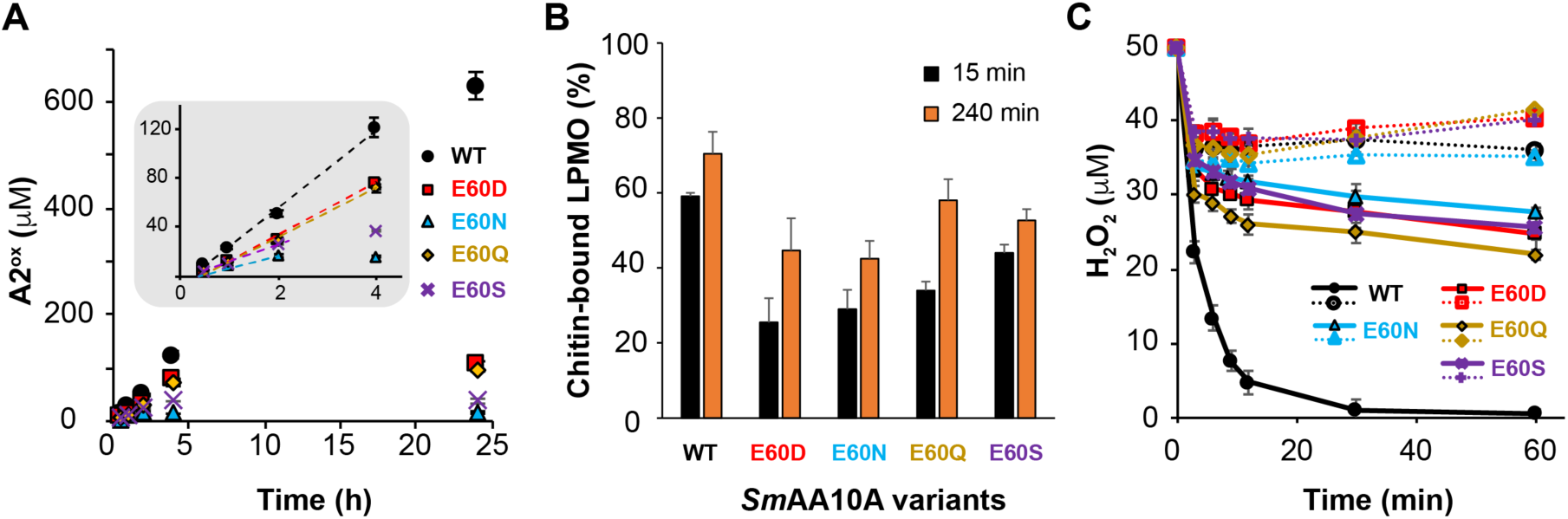
Biochemical characterization of *Sm*AA10A and Glu60 mutants. Time-course monitoring for **(A)** the release of soluble chitobionic acid (A2^ox^) from β-chitin in presence of AscA (1 mM) and **(B)** binding to β-chitin by *Sm*AA10A wildtype and mutants (1 µM) (see **Figure S24** for entire time-courses). The inset (grey zone) in panel A is a zoom-in view of the first 4 hours to determine the initial oxidative rate (dotted lines) of each enzyme, equal to 0.53, 0.34, 0.15, 0.33 and 0.26 min^-1^ for *Sm*AA10A-WT, E60D, N, Q and S, respectively. **(C)** H_2_O_2_ consumption (50 µM at t_0_) by *Sm*AA10A wildtype and mutants thereof (50 nM) in presence (solid lines) and absence (dotted lines) of β-chitin. Reactions were initiated by addition of AscA (20 µM). Note that for comparative purposes the data shown here for *Sm*AA10A-WT are a reproduction of data displayed in **Figure 2B**. All reactions were carried out in sodium phosphate buffer (50 mM, pH 7.0) with β-chitin (10 g.L^-1^) and incubated at 40 °C in a thermomixer (1000 rpm). The color/symbol code is provided in each panel. Error bars show ± s.d. (n = 3, independent experiments).

## DISCUSSION

When LPMOs were identified as monocopper enzymes, it soon became apparent that they may work differently than hitherto characterized copper enzymes due to their peculiar structure and lack of additional redox-active cofactors. In this study, we present complementary experimental and computational data shedding light on the still debated^1,46^ LPMO reaction. Importantly, while cellulose active LPMOs have received most of the attention so far, our study proposes the first atomistic-level description of the mode of action of solid-state chitin active LPMOs, whose functionalities in Nature, notably in pathogenesis mechanisms,^19,22,47^ only are now being revealed.

We demonstrated by performing two complementary experiments, that not only cellulose, as shown in previous work,^1,46^ but also chitin oxidation by LPMOs, depends mainly (if not only) on H_2_O_2_ availability in standard aerobic conditions (**Figure 1**). Two recent computational (QM/MM) studies have addressed the reaction between H_2_O_2_ and the LPMO-bound Cu(I),^32,40^ both starting with a crystal structure of the AA9 C4-oxidizer from *Lentinus similis* (*Ls*AA9A) obtained in complex with cellotriose (PDBid 5ACF).^33^ While we suggest that the torsional strain imposed on the H_2_O_2_ molecule (dihedral angle = 53°) in *Sm*AA10A in state **1** is important for the progress of the reaction, QM/MM models obtained by Wang et al. suggesta torsional *unstrained* H_2_O_2_ molecule (dihedral angle = 116°) that is 2.8 Å away from the active site Cu(I), hydrogen bonded to a glutamine side chain (analogous to the *Sm*AA10A Glu60) and a water molecule.^32^Differences in the second sphere environment between AA10 and AA9s may explain such difference. However, we also observe that the QM region in both QM/MM studies on *Ls*AA9A is composed of different residues, which we think impacts significantly the outcome of calculations (see **Supplementary Discussion and Figure S26**). Wang et al. identified a transition state akin to state **2** presented in this work with an energy barrier of 5.8 kcal mol^-1^ (B3LYP). This value is lower than the one calculated for our *Sm*AA10A-chitin model (12.8 kcal mol^-1^ (B3LYP) and 8.6 kcal mol^-1^ (TPSSh)). The two previous computational studies suggest substrate H-abstraction energy barriers (states **5** to **6**; **Figure 3**) of 5.5 kcal mol^-1^ (B3LYP)^32^ and 16.5 kcal mol^-1^ (TPSS)^40^ whereas we obtained values of 13.0 kcal mol^-1^ (B3LYP) and 10.6 kcal mol^-1^ (TPSSh) for the *Sm*AA10A-chitin model. Not only the use of different DFT-functionals may explain the discrepancies observed between the two different *Ls*AA9A-cellotriose models^32^,^40^, but it is also apparent that the cellotriose ligand has been re-positioned during model preparation, resulting in complexes different from the crystal structure (**Figure S26A**).

While Wang et al.^32^ report that all the steps of the LPMO catalyzed reaction take place on a singlet energy surface, HedegÅrd & Ryde^32,40^ identify two intermediates that have triplet ground states, those resembling states **5** and **7** in this work. Although the QM/MM/MD simulation (**Movie S1**) of the *Sm*AA10A-chitin model, that provided the underpinning molecular geometries for our mechanistic approach, also progressed on a singlet surface, subsequent refinement provided more details. We found that state **4** and **5** have triplet ground states, and this was confirmed by unrestricted corresponding orbital (UCO) analysis (zero UCO overlap). A previous study, employing a higher level of theory (CASSFC/CASPT2) than DFT, reported that the copper-oxyl complexes of various (not LPMO related) model compounds had triplet ground states,^48^ supporting our findings for state **5**. Similarly to some studies on other metalloenzymes,^2^ our results indicate that LPMOs-catalyzed chemistry would rely on spin-crossover transitions (here in a singlet-triplet-singlet sequence) to reduce energetic barriers of reactions involving unpaired electrons. Despite differences in TS energy barriers and spin states, our study together with work on *Ls*AA9A-cellotriose models^32,40^ point collectively towards a LPMO peroxygenase reaction that is thermodynamically feasible and kinetically more favorable than a monooxygenase reaction, this for different families of LPMO and types of substrate.

The nature of the rate limiting step of LPMO catalysis, and the order of catalytic events are not settled. So far, only one study has estimated the overall steady-state kinetic parameters of LPMO catalysis in presence of H_2_O_2_, precisely for *Sm*AA10A (*k*cat = 6.7 s^-1^ *K*_M_, _H2O2_ ~ 3 μM).^34^ For comparative purposes, the few available kinetic studies carried out under O_2_ conditions (i.e. no added H_2_O_2_) reported catalytic efficiencies on soluble substrates more than two orders of magnitude lower than that of the peroxygenase reaction.^33,49^

Here, we took advantage of a) the fluorophore-properties of the LPMO-Cu(I)/Cu(II) redox couple previously described by Bissaro et al.,^43^ and b) an experimental approach where we worked with isolated species to dissect reduction and re-oxidation catalytic events. Our data indicate that the reduction rate is unlikely to be a rate-limiting event, provided that sufficient amount of reducing power is present in the reaction. Indeed, using AscA > 20 μM (the lowest AscA concentration employed here) would yield rates > 8 s^-1^, which is slightly faster than the *k*_cat_ determined for *Sm*AA10A.^34^ Under standard aerobic steady-state conditions, the reduction step is definitely not rate-limiting since mM concentrations of reductant are usually employed.^16^ The reduced *Sm*AA10A-Cu(I) is the precursor for all suggested LPMO catalytic pathways.^1,23,24^ The facts that the LPMO-Cu(I) form binds more strongly to substrates than LPMO-Cu(II) ^37,49,50^ and that access to the active site is severely hindered when the LPMO is bound to substrate^27^ suggest that LPMO reduction occurs in solution. In agreement, our stopped-flow experiments indicate that *Sm*AA10A reduction is not significantly influenced by the presence of carbohydrate substrate(**Figure S12**). Furthermore, our data on the lifetime of LPMO-Cu(I) exposed to H_2_O_2_ (**Figure S19**) show that productive catalysis with H_2_O_2_ (i.e. peroxygenation) most certainly occur when the reduced enzyme is pre-bound to the carbohydrate substrate. Nevertheless, depending on the rate of association to substrate, the reduced LPMO active site will be exposed to oxidants such as O_2_ and H_2_O_2_ before it settles on the carbohydrate surface. Thus, as an initial experiment, O_2_ and H_2_O_2_ reactivity with *Sm*AA10A-Cu(I) was monitored in absence of substrate. The second order rate constant for the reaction with O_2_, *k*_O2_ = 3.33 M^-1^ s^-1^, is surprisingly low (see discussion below), a typical reaction proceeding over several minutes with an enzyme:O_2_ molar ratio of 1:100. This is in striking contrast with what has been reported for an AA9 C1/C4 oxidizer from *Thermoascus aurantiacus* (*Ta*AA9A), that was shown to be re-oxidized in seconds (~0.15 s^-1^) at an enzyme:O_2_ molar ratio of ca. 1:1.^38^ It is a possibility that this difference in rates underpins a more general, fundamental difference between AA9s and AA10s with respect to O_2_ activation pathways. In the frame of the Marcus theory, we can predict the rate of outer-sphere electron transfer between LPMO-Cu(I) and O_2_, as outlined by Kjaergaard et al.^38^ (See **Supporting discussion for details**). Thereby, we estimate an electron transfer (ET) rate *k*_ET_, O2 = 3.39 M^-1^ s^-1^, a value remarkably similar to the experimentally determined *k*_O2_ = 3.33 M^-1^ s^-1^. This result implies that reduction of O_2_ would be subject to an outer-sphere reduction, in contrast to an inner sphere reduction suggested by data on *Ta*AA9A.^38^ An outer-sphere reduction entails that O_2_ would be first reduced in a “pre-bound” configuration and only then the resulting O_2_^•-^ and Cu(II) would form a covalent bond. Such pre-bound O_2_ has been suggested by neutron protein crystallography and DFT calculations for an AA9,^51^ but not for any AA10 yet.

Comparing the *k*_O2_ value of *Sm*AA10A with other “classical” monooxygenases provides some interesting insights. Indeed, bimolecular rate constants for re-oxidation by O_2_ are several orders of magnitude higher (10^5^-10^7^ M^-1^ s^-1^) for bi-nuclear copper enzymes such as tyrosinases^2^ and dopamine monooxygenase,^52^, monocopper amine oxidase^53^ and galactose oxidases,^54^ as well as some multi-copper laccases.^55,56^ In light of this comparison, together with steady-state kinetics studies^34^ and QM/MM results presented here, we conclude that *Sm*AA10A has not been evolved to act as a O_2_-using oxygenase.

Regarding the reaction of *Sm*AA10A-Cu(I) with H_2_O_2_, we determined a second order rate constant of 6.85×10^3^ M^-1^ s^-1^, which is three orders of magnitude faster than the corresponding reaction with O_2_. Also, in the second order rate constant plots (**Figure 5B&D**), the y intercept is close to zero when using H_2_O_2_ as oxidant but not when using O_2_. This is indicative of a reaction of *Sm*AA10A-Cu(I) with H_2_O_2_ that is mainly irreversible while it denotes a certain extent of reversibility with O_2_. The reaction with H_2_O_2_ resembles a Fenton reaction, generating highly reactive hydroxyl radicals (**Figure S18**) while consuming equimolar amounts of *Sm*AA10A-Cu(I) and H_2_O_2_ (**Figure S17**). We have no indications that a Cu-oxyl intermediate is formed in the reaction between *Sm*AA10A-Cu(I) and H_2_O_2_ in solution, and we could not detect any long-lived Cu-oxyl intermediate that could bind and cleave substrate after it was formed (**Figure S19**). Remarkably, it appears that the reaction between *Sm*AA10A-Cu(I) and H_2_O_2_ is slower in solution than in presence of substrate. If we compare the V_max_ determined by Kuusk et al.^34^ (at saturating [H_2_O_2_] of 20 μM) with the reaction rate calculated from the second order rate constant *k*^H^2^O^2^ determined here, it appears that the reaction with H_2_O_2_ is accelerated by about 50 times in presence of substrate (*k*_cat_ × [E] / *k*^H^2^O^2^ × [E] × [H_2_O_2_] = 6.7 s^-1^ / 6.85×10^3^ M^-1^ s^-1^ × 20 μM = 48.9). Repeating single turnover re-oxidation experiments in presence of NAG_6_ showed that re-oxidation by O_2_ was slowed down whereas, re-oxidation by H_2_O_2_ was boosted in a NAG_6_ concentration-dependent manner. Considering these observations in light of the computational data that predict steric clashes between the substrate and superoxide (**Figure S9**), we may speculate that O_2_ and NAG_6_ compete for the active site. On the other hand, the accelerated re-oxidation observed with H_2_O_2_ and NAG_6_ indicate that oxidative cleavage of insoluble chitin and NAG_6_ follow the same reaction path. In our QM/MM models, the roles of interactions with the substrate appear to be threefold along the reaction coordinates: (i) together with Glu60, the substrate H-bond assists H_2_O_2_ molecule positioning in the reaction cavity (state **1**; **Figure 3B**) and (ii) helps stabilizing the reactive “precision-guided HO^•^” (states **2** – **4**); (iii) these interactions seem to position the H_2_O_2_ derived water molecule that contributes with a H-bond to the reactive species in the final stages of the reaction mechanism (states **5** – **9**) (**Figure S9**). Thereby, we suggest a substrate-assisted mechanism in which substrate H-bond donation facilitates the initial one-electron reduction of H_2_O_2_ in the *Sm*AA10A-chitin complex and thus results in an increased reaction rate in presence of substrate, in agreement with ternary complex formation predicted by steady-states kinetic.^34^ The cellulose “analog” of the chitin amide group that could act as hydrogen bond donor is the 2’-OH moiety, which is hydrogen bonded to the adjacent chain glucose residue and not readily available for H-bonding to H_2_O_2_. It is not known if the presence of substrate increases the reaction rate of AA9 enzymes with H_2_O_2_. When the *Sm*AA10A Glu60 residue was mutated to Gln, Asp, Asn and Ser, the ability of the Cu(I) site to react with H_2_O_2_ in absence of substrate was retained (**Table S3**). Nevertheless, the catalytic efficiencies of these mutants with insoluble chitin substrate decreased considerably (**Figure 7**), indicating that formation of a productive ternary complex was obstructed by altering the H_2_O_2_ interacting environment in the reaction cavity.

After it was suggested that H_2_O_2_ is the relevant co-substrate for LPMOs,^1^ only one hypothesis relating to the rate limiting step of the reaction has appeared in the literature. In their hypothesis, Wang et al. proposed that (non-enzymatic) hydrolysis of the hydroxylated carbohydrate, i.e. state **9** in this work, is the rate limiting step. The rationale behind this suggestion is that they found, by computational methods, a high transition state energy barrier for the glycosidic bond cleavage reaction (18.2 kcal/mol) that coincided with *Ls*AA9A catalytic constant (*k*cat = 0.11 s^-1^) determined under ambient O_2_ (i.e. uncontrolled, limiting H_2_O_2_) conditions. In spite of limitations when attempting to relate energy barriers to catalytic constants (see **Supplementary discussion**), it is a possibility, awaiting experimental validation, that such hypothesis holds for LPMOs acting on soluble substrates and using O_2_ as co-substrate. For polysaccharide-active LPMOs, both from AA9 and AA10 families, there are multiple experimental evidences that the apparent LPMO rate, under stationary conditions (i.e. no excess of H_2_O_2_), is limited by the rate of *in situ* production or supply of H_2_O_2_.^1,34,46,57^ Therefore, in biologically-relevant settings, where low steady-state concentrations of H_2_O_2_ are expected, LPMO catalysis will be mainly driven by the probability of encounter between H_2_O_2_ and substrate-bound pre-reduced LPMOs. Under H_2_O_2_ saturating conditions, it is not likely that the LPMO-Cu(II) to LPMO-Cu(I) reduction (**Figure 8**, **step 1**) is the rate limiting step when sufficient amounts of reductant are available (see above). As summarized in **Figure 8,** we propose the following order of events for the LPMO peroxygenase reaction. **Step 1**: the substrate-dissociated enzyme is reduced via close contact between the enzyme copper site and the reducing species. **Step 2**: the LPMO binds in a specific manner, positioned to facilitate formation of a ternary complex. **Step 3**: H_2_O_2_ diffuses freely into the reaction cavity via a tunnel access and is guided into a strained position, optimal for H_2_O_2_ reduction, by specific hydrogen bonding interactions. **Steps 4-7**: the reaction progress as indicated by states **1** to **9** described in this work. **Steps 8** and **9**: glycosidic bond cleavage (via elimination reaction) and enzyme dissociation from the substrate occur in an order that remains to be clarified. **Steps 10-10’**: in a substrate-free case, the LPMO can engage into inactivation processes provided that H_2_O_2_ accumulated in the system. It is interesting to note that *k*H2O2 is nearly one order of magnitude higher than the apparent inactivation rate previously determined in presence of excess of reductant (10^3^ M^-1^ s^-1^),^34^ indicating that this reaction (**steps 10**) is not the rate limiting factor in the inactivation process. Our data also indicate that multiple turnovers are required to inactivate the enzyme (**Figure S20**). **Steps 11-11’**: Substrate-free LPMOs can also engage into H_2_O_2_ production pathway. In the latter, the reactivity of *Sm*AA10A-Cu(I) with O_2_ seems to be the rate-limiting step since the second order rate of 3.3 M^-1^ s.^-1^ yields a turnover of 8.10^-4^ s^-1^ at ambient [O_2_], a value strikingly similar to the apparent H_2_O_2_ production rate.^16^

**Figure 8.**
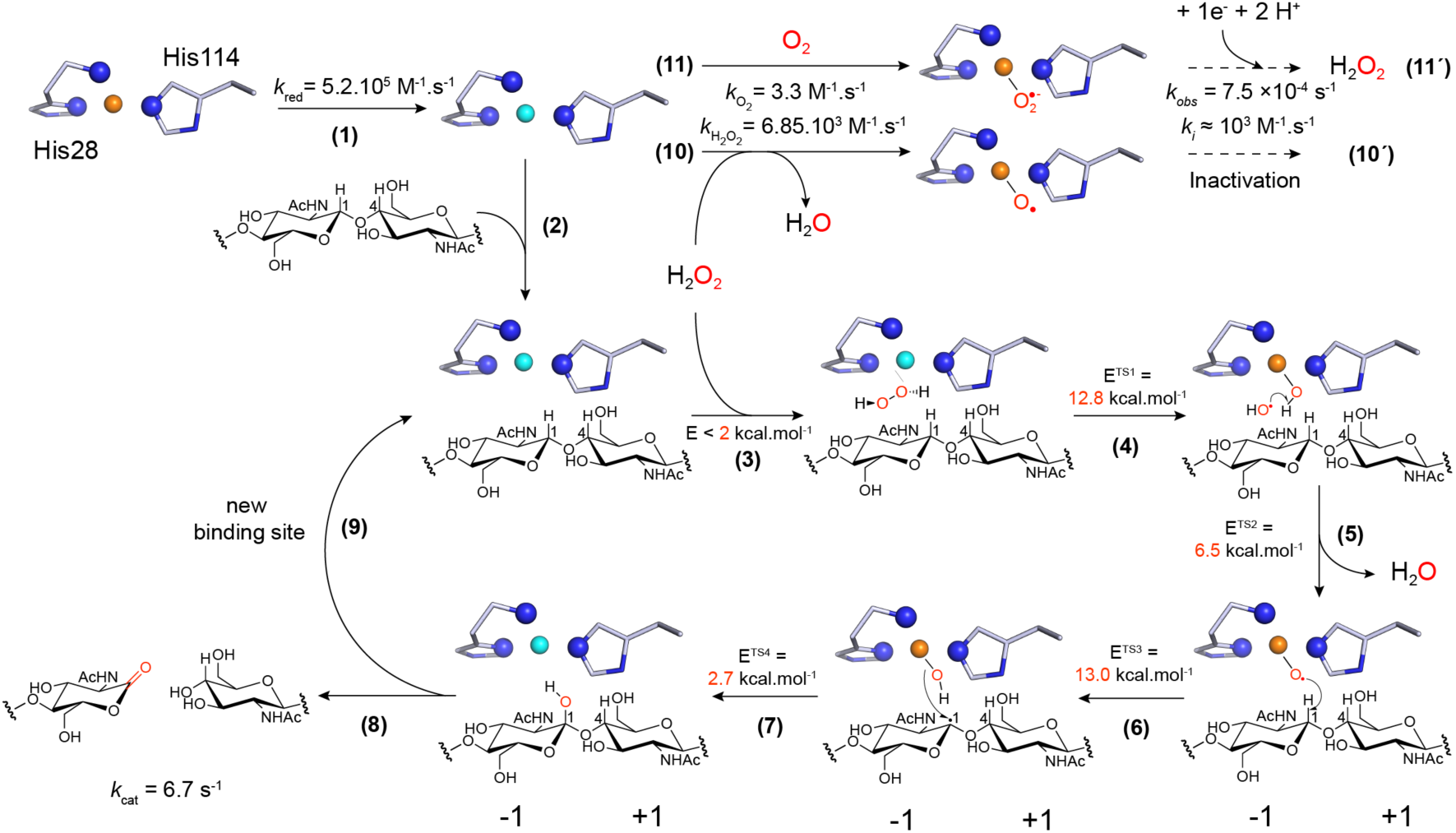
Proposed molecular mechanism of chitin oxidation by *Sm*AA10A and associated rate constants or energy barriers. Step 1: the resting LPMO-Cu(II) is reduced to the LPMO-Cu(I) active state (4.23 x 10^5^ M^-1^ s^-1^;i.e. 423 s^-1^ or 8.5 s^-1^ when using 1 mM or 20 µM AscA, respectively). **Step 2**: polysaccharide binding. **Step 3**: H_2_O_2_ diffusion to the active site cavity through a tunnel access. **Step 4**: reaction of H_2_O_2_ with Cu(I) inducing homolytic bond cleavage yielding a Cu(II)-OH species and a hydroxyl radical. **Step 5**: the hydroxyl radical reacts with Cu(II)-OH yielding a water molecule and a [CuO]^+^intermediate. **Step 6**: [CuO]^+^catalyzes HAA from the substrate. **Step 7**: hydroxylation of the C1 carbon via oxygen-rebound mechanism.^29^ **Step 8**: The hydroxylated product undergoes an elimination reaction,^26^ suggested to be enzyme independent,^32^ inducing glycosidic bond cleavage. **Step 9**: The LPMO-Cu(I) can enter a new catalytic cycle. **Step 10:** substrate-free LPMO-Cu(I) can react with H_2_O_2_ (6.85 x 10^3^ M^-1^ s^-1^) leading eventually, after multiple turnovers, to enzyme inactivation (**step 10’**, ca. 10^3^ M^-1^ s^-1^)^34^. **Step 11:** The LPMO-Cu(I) can also react with O_2_ (3.3 M^-1^ s^-1^; i.e. 8.10^-4^ s^-1^ at 250 µM (atmospheric) O_2_) leading *in fine* to H_2_O_2_ production (**step 11’**, 7.5.10^-4^ s^-1^).^16^ Note that steps 10’ and 11’ proceed via mechanisms that remain to be fully elucidated.

The experimental conditions of the LPMO reaction will affect the outcome of the chain of events. With low affinity substrates and high concentrations of H_2_O_2_ a single turnover may be observed. At optimal conditions, where reducing equivalents and H_2_O_2_ are delivered in a timely fashion in presence of a suitable substrate, multiple turnovers are expected before the enzyme is inactivated.^57^ The proposed substrate-assisted mechanism provides a competitive advantage to polysaccharide-bound LPMOs for turning over H_2_O_2_ compared to the reaction with H_2_O_2_ in solution that leads to enzyme inactivation via formation of hazardous hydroxyl radicals

## CONCLUSIONS

We have presented how LPMOs can harness the intrinsic oxidative power of H_2_O_2_, in a controlled and substrate-assisted manner to oxidize crystalline chitin. Our study reveals how unique LPMO chemistry is, since relying on the formation of a confined active site reaction cavity upon association of the enzyme with the polysaccharide. Resembling what we first described as an enzyme-guided Fenton reaction,^28^ the formation of a ternary complex provides the adapted environment to allow efficient H_2_O_2_ reduction and generation of a “precision-guided HO^•^”, leading to a reactive [CuO]^+^ core. In the context of frontier molecular orbital theory, we presented in great details the electronic features of the reactivity of H_2_O_2_ with LPMO-Cu(I), which will assist future spectroscopic studies. We have also shown the occurrence of spin crossover events along the reaction coordinates, suggesting that the multiplicity can be tuned to cope with energetic barriers of reactions involving unpaired electrons. We reported for the first time single turnover kinetics of reactions of LPMO-Cu(I) with O_2_ and H_2_O_2_, indicating, together with steady-state kinetics and computational simulations, that the use of O_2_ as co-substrate by *Sm*AA10A to oxidize chitin is very unlikely and supporting the peroxygenase paradigm. Questioning the universality of this mechanism across LPMOs families will no doubt be a topic of intense investigations for the coming years.

We foresee that the molecular basis of this unusual, copper-based chemistry catalyzed by LPMOs will inspire and assist the design of new model complexes for oxy-functionalization of inert C-H bonds. Furthermore, witnessing strong indications of the chitinolytic role of LPMOs in pathogenesis and defense mechanisms, we believe our study will set foundations for a better understanding of these biologically important questions.

## ASSOCIATED CONTENT

### Supporting information

The supporting information contains supplemental **Figures S1-26, Tables S1-4, Scheme S1, Movie S1 and Movie S2.**

## AUTHORS INFORMATION

### Corresponding authors

*\**

## Conflicts of interest

The authors declare no conflicts of interest with the contents of this article.

### Abbreviations

The abbreviations used are:

AscA: Ascorbic acid
AR: AmplexRed®
A2^ox^: Chitobionic acid or GlcNAcGlcNAc1A
GH: Glycoside hydrolases
HAA: Hydrogen atom abstraction
HRP: Horseradish peroxidase
LPMO: Lytic polysaccharide monooxygenases
*Ls*AA9A: LPMO9 from *Lentinus similis*
MD: Molecular dynamics
MM: Molecular mechanics
NAG_6_: Hexa-*N*-acetyl-chitohexaose
NEB: Nudged elastic band
QM: Quantum mechanics
*Sm*AA10A (or CBP21): LPMO10 from *Serratia marcescens*
*Sm*GH20: Chitobiase from *Serratia marcescens*
EPR: Electron paramagnetic resonance
SOMO: single occupied molecular orbital
HOMO: highest occupied molecular orbital
LUMO: Lowest unoccupied molecular orbital

## ACKNOWLEDGMENT

This work was supported by the Research Council of Norway grants 240967 (ÅKR) and 269408 (VE), and National Science Foundation grant MCB1715176 (JB). Computational work was performed on the Abel Cluster, owned by the University of Oslo, and the Norwegian metacenter for High Performance Computing (NOTUR) and the Extreme Science and Engineering Discovery Environment (XSEDE), which is supported by National Science Foundation grant number ACI-1548562, through allocation TG-MCB090159 to GTB. This work authored in part by Alliance for Sustainable Energy, LLC, the manager and operator of the National Renewable Energy Laboratory for the U.S. Department of Energy (DOE) under Contract No. DE-AC36-08GO28308. Funding was provided to GTB by the U.S. Department of Energy Office of Energy Efficiency and Renewable Energy the Bioenergy Technologies Office.

