## Supplemental information for "Molecular mechanism of the chitinolytic monocopper peroxygenase reaction"

### Molecular mechanism of the chitinolytic mono-copper peroxygenase reaction

**This PDF file includes:**

1. Complete experimental and computational details
2. Supplementary results
3. Supplementary discussion
4. List of supplementary figures
5. Supplementary Figure 1 to 26
6. Abbreviations list
7. Supplementary references
8. QM/MM optimized xyz coordinates of states **1-9**

### 1. Complete experimental and computation details

***Materials.*** Most of the chemicals were purchased from Sigma-Aldrich. β-chitin extracted from squid pen was purchased from France Chitin (Orange, France). Ascorbic acid (AscA; 100 mM) stock solutions was prepared in metal-free water (Trace SELECT®, Sigma-Aldrich), aliquoted and stored at -20 °C and thawed in the dark for 10 min just before use. $H_2O_2$ (35%) was purchased from Merck, aliquoted and stored at -20 °C, and its concentration was systematically controlled for each experiment by measuring the absorbance at 240 nm (extinction coefficient of 43.6 $M^{-1}$ $cm^{-1}$).

***Site-directed mutagenesis.*** The plasmid pRSETB containing the gene encoding for *Sm*AA10A-WT was used as template for site-directed mutagenesis using the QuikChange II XL site-directed mutagenesis kit (Agilent Technologies) and PCR primers listed in **Table S4**. Mutated plasmids were verified by sequencing and thereafter used for protein expression and purification in the same way as for the wild-type enzyme (see below).

***Production and purification of recombinant LPMOs.*** The recombinant LPMO10A from *Serratia marcescens* (*Sm*AA10A or CBP21) and mutants thereof were produced and purified according to previously described protocols.[1] All LPMOs used in this study were prepared in sodium phosphate buffer (50 mM, pH 7.0), copper-saturated with Cu(II)$SO_4$ and desalted (PD MidiTrap G-25, GE Healthcare) before use.[2] The concentration of *Sm*AA10A-WT and Glu60 mutants thereof was determined by measuring the absorbance at 280 nm and using an extinction coefficient of 35,200 $M^{-1}$ $cm^{-1}$. The concentration of horseradish peroxidase (HRP) was determined by measuring the absorbance at 403 nm and using an extinction coefficient of 102,000 $M^{-1}$ $cm^{-1}$.

**Sm*AA10A activity test.*** Reactions were carried out in 2 mL Eppendorf tubes and the reaction volume was 200 µL (for final time point analysis) or 500 µL (for time-course monitoring). Typical reactions contained the LPMO (1 µM) and β-chitin (10 g.$L^{-1}$), mixed in sodium phosphate buffer (pH 7.0, 50 mM) and were pre-incubated during 20 min at 40 °C in a Thermomixer (1000 rpm). Then, the reaction was initiated by adding AscA (to a final concentration of 1 mM). Experiments carried out to evaluate the effect of HRP were prepared as described above except that HRP (0-730 nM final concentration) and AmplexRed® (200 µM final concentration) were added to the mixture before initiation of the reaction by addition of AscA (1 mM). In control reactions, *Sm*AA10A was replaced by Cu(II)$SO_4$ (1 µM). For time course monitoring, 55 µL samples were taken from the reaction mixtures at regular intervals and soluble fractions were immediately separated from the insoluble substrate by filtration using a 96-well filter plate (Millipore) operated with a vacuum manifold. Samples were then frozen (-20 °C) prior to further analysis.

Quantitative product analysis was performed by incubating the soluble products with 2 µM of a chitobiase from *Serratia marcescens* (also known as *Sm*GH20A) at 37 °C overnight in order to convert the LPMO products to *N*-acetylglucosamine (GlcNAc) and chitobionic acid ($A2^{ox}$ or GlcNAcGlcNAc1A). The hydrolyzed samples were analyzed by high performance anion exchange chromatography (HPAEC) coupled to pulsed amperometric detection (PAD) using a Dionex Bio-LC equipped with a CarboPac PA1 column as previously described.[3] To quantify $A2^{ox}$, a standard was produced in-house by treating chitobiose (Megazymes) with a chitooligosaccharide oxidase (ChitO) from *Fusarium graminearum*, which yields 100% conversion of chitobiose to chitobionic acid.[2,4] All chromatograms were recorded using Chromeleon 7.0 software.

***Chitin binding assay.*** The capacity of *Sm*AA10A-WT and mutants thereof to bind β-chitin was tested by suspending 10 mg/mL of substrate in sodium phosphate buffer (50 mM, pH 7.0) in a total volume of 600 μL in 2 mL Eppendorf tubes. Reactions were started by the addition of *Sm*AA10A (1 μM final concentration) and were incubated and stirred in an Eppendorf Comfort Thermomixer (at 40 °C, 1000 rpm). Samples were taken (100 μL) after 15, 30, 60, 120 and 240 min and immediately filtrated using a 96-well filter plate (Millipore) operated with a vacuum manifold to obtain the unbound protein fraction. In order to assess the percentage of bound proteins to the substrate, control samples with only enzyme and buffer were included, representing the maximum quantity of protein present in the samples (i.e. 100% unbound). The protein concentration in each sample was determined using the Bradford assay (Bio-Rad, Munich, Germany).

***$H_2O_2$ consumption experiments.*** $H_2O_2$ consumption by *Sm*AA10A-WT and mutants thereof was measured according to a previously described protocol[5] using conditions that were slightly different from the standard reaction conditions described above: in order to be able to monitor the $H_2O_2$ consumption within a reasonable timescale the enzyme concentration had to be reduced and EDTA was added to reduce the background reaction of free metals-catalyzed $H_2O_2$ reduction (see **Figure S6**). After optimization, a standard reaction mixture contained the LPMO (50 nM) and $H_2O_2$ (100 μM) and EDTA (50 μM), without or with β-chitin (10 $g.L^{-1}$), in sodium phosphate buffer (50 mM, pH 7.0), and the mixtures were incubated at 40 °C in a thermomixer (1000 rpm). The reactions were initiated by addition of AscA (20 μM final concentration). At regular intervals (t = 3, 6, 9, 12, 30 and 60 min), 70 μL of the reaction mixture was sampled, filtered as described above and 25 μL of the filtrate was mixed with 75 μL of a pre-mix of HRP (5 $U.mL^{-1}$ final concentration) and Amplex® Red (ThermoFisher) (100 μM final concentration) in sodium phosphate buffer (50 mM pH 7.0). $H_2O_2$ concentrations waere then determined spectrophotometrically by measuring the absorbance at 540 nm in a microtiter plate reader. An $H_2O_2$ standard curve was prepared in the same conditions.

***Bioinformatics analysis.*** The sequence of the chitin-binding protein from the fern *Tectaria macrodonta* (Tma12; GenBank ID AFR32946.1) together with the sequences of the catalytic domains of 19 selected LPMOs were aligned using the T-Coffee Expresso online tool.[6] The structures of *Sm*AA10A (PDB 2BEM),[1] of the AA9A from *Thermoascus aurantiacus* (*Ta*AA9A, PDB 2YET),[7] and of the AA13A from *Aspergillus orizae* (*Ao*AA13A, PDB 4OPB)[8] were supplied as additional structural data input to the T-Coffee algorithm. The resulting multiple sequence alignment (MSA) was employed as input to build a phylogenetic tree, using PhyML (bootstrapping procedure with default substitution model and

100 bootstraps) available via the online platform Phylogeny.fr.[9] The phylogenetic tree was visualized using the iTOL platform.[10] A structural model of Tma12 was generated using the online Protein Homology/analogY Recognition Engine V 2.0 (Phyre2).[11]

***Isolation of LPMO-Cu(I).*** In a typical experiment, solutions of AscA (100 µL at 100 mM) and *Sm*AA10A (300 µL at 75 µM) were submitted to 3 cycles (10 min/2 min) of vacuum/$N_2$ using a Schlenk line before being transferred into an anaerobic chamber (Whitley A35 anaerobic workstation). All buffer and water solutions were extensively flushed with $N_2$. Following this first $O_2$ removal, all solutions were placed in the anaerobic chamber for at least 16 hours to ensure complete $O_2$-free conditions (the lids of the vessels were slightly loose). In the anaerobic chamber, the LPMO was reduced by adding 20 eq. of AscA (i.e. 4.5 µL of the 100 mM solution) to the LPMO solution, followed by incubation for 10 min. The excess of AscA was then removed by desalting the reduced enzyme on a PD MidiTrap G-25 column (GE Healthcare) that had been equilibrated beforehand with anaerobic sodium phosphate buffer (50 mM, pH 7.0). This procedure yielded 1 mL of a ca. 40 µM LPMO-Cu(I) solution. An aliquot (20 µL) of the latter solution was then sampled to determine the exact enzyme concentration by measuring absorbance at 280 nm.

***Monitoring reduction of LPMO-Cu(II) by AscA.*** Prior experiments had established that changes in intrinsic fluorescence of *Sm*AA10A can be used to monitor the oxidation state of the Cu center, with the Cu(I) form of the enzyme having greater fluorescence intensity than the Cu(II) form.[12] Fluorescence was measured using a KinetAssyst stopped-flow spectrometer (TgK Scientific, Bradford-on-Avon, UK) in single mixing mode with fluorescence detection using an excitation wavelength of 280 nm and a bandpass filter that allows for fluorescent light above 320 nm to be detected at the photomultiplier tube with an applied voltage of 600-750 V. The protein sample was sealed in an airtight tonometer interfaced with the stopped flow sample handling unit. Deoxygenated buffer and ascorbate were prepared in the anaerobic chamber, sealed in gastight syringes, and then introduced to the sample handling unit of the stopped-flow spectrophotometer for reaction. Anaerobic *Sm*AA10A (5 µM, 50 mM phosphate buffer, pH 7.0,) was rapidly mixed (< 5 ms) with an equivalent volume of variable concentrations of degassed anaerobic ascorbate (5 –500 µM after mixing), at 25 °C, and fluorescence changes were then monitored over time.

***Re-oxidation of LPMO-Cu(I) by $H_2O_2$.*** Most kinetic experiments for measuring changes in fluorescence emission upon re-oxidation of LPMO-Cu(I) were carried out using a SFM4000 stopped-flow spectrophotometer (BioLogic Science Instruments, Grenoble, France). Data were measured using a KinetAssyst stopped flow spectrometer (Hi-Tech Scientific) in single mixing mode with fluorescence detection using an excitation wavelength of 295 nm and a bandpass filter that allows for fluorescent light above 320 nm to be detected at the photomultiplier tube (PMT) with an applied voltage of 650 V. Prior to measurements, the spectrophotometer was made anaerobic by flushing the entire system with a dithionite solution (large excess) and, subsequently, anaerobic sodium phosphate buffer (50 mM, pH

7.0). In the anaerobic workstation, the isolated LPMO-Cu(I) (see above) was diluted into anaerobic sodium phosphate buffer (50 mM, pH 7.0) to prepare 5 mL of a 4 µM LPMO-Cu(I) solution, which was transferred into a sealed gastight syringe. $H_2O_2$ solutions were made via serial dilution of the 35% (ca. 10.8 M) $H_2O_2$ stock solution in anaerobic buffer to generate working stock solutions with concentrations of 10 – 800 µM. $H_2O_2$ solutions were prepared in gastight syringes in the anaerobic chamber, and then introduced to the sample handling unit of the stopped-flow spectrophotometer for reaction. Following rapid stopped-flow mixing of the reduced protein with oxidant, the reactions with $H_2O_2$ were monitored via time resolved fluorescence. The concentration of prepared $H_2O_2$ solutions was systematically controlled spectrophotometrically as described above. Note that the reactivity with $O_2$ was so slow (up to 120 min) that an alternative, simpler method was developed (see below).

***Re-oxidation of LPMO-Cu(I) by $O_2$; alternative method.*** Solutions of $O_2$ with different concentrations were prepared by sparging solutions of sodium phosphate buffer (50 mM, pH 7.0) for 10 min with variable $N_2/O_2$ gas mixtures (at 200 mL/min total gas flowrate) in glass bottles (20 mL), which were sealed at the end of the sparging period. The actual $O_2$ concentration was then measured using an oxygen sensor (a micro fiber optic oxygen transmitter, OXY-4 micro, PreSens, Germany) as previously described.[5] The $O_2$ concentration in each bottle was controlled before each set of experiments and bottles were re-sparged when necessary. Initially, re-oxidation experiments were performed using the stopped-flow spectrophotometer similarly to experiments described above for $H_2O_2$. However, given the slowness of the reaction we developed a simpler approach. In the anaerobic workstation, the isolated LPMO-Cu(I) (see above) was diluted into anaerobic sodium phosphate buffer (50 mM, pH 7.0) to prepare 500 µL of a 4 µM LPMO-Cu(I) solution, which was transferred into a sealable fluorescence Quartz cuvette equipped with screw cap and septum (Hellma). The sealed cuvettes were then inserted into a Cary Eclipse Fluorescence spectrophotometer (Agilent Technologies). Excitation and emission wavelength were set to 280 and 340 nm and the PMT detector voltage was set to 600 V. The reaction was initiated by adding into the cuvette, via a Hamilton syringe, 500 µL of an appropriate $O_2$ solution (final $O_2$ concentrations after mixing were 0-600 µM)). The change in fluorescence was monitored over a period of 180 min, with data points recorded every 10 sec. With this method, we could run simultaneously 3 replicates and 1 control in which anaerobic buffer was added instead of the $O_2$ solution.

***Kinetics data analysis.*** Fluorescence data were expressed as normalized fluorescence, i.e. as $1-\frac{\Delta F}{\Delta Fmax}$ $= 1-\frac{F_{max}-F(t)}{F_{max}-F_0}$, where $F_{max}$ and $F_0$ are the fluorescence signals of fully reduced and ground state LPMO, respectively. Note that $1-\frac{\Delta F}{\Delta Fmax}=\frac{LPMO-Cu(I)}{LPMO_{tot}}$ when considering LPMO-Cu(I) and LPMO-Cu(II) as the two main fluorophores and provided that the fluorescence signal of both species is linear with concentration (as is indeed the cases; **Figure S11**) (see Bissaro et al.[12] for further details concerning these equations). Single exponential decay functions with a baseline correction factor ($y = a.e^{-k_{obs}.t} + c.t + d$) were fit to the data (with an in-house Matlab script) to determine first order rate constants ($k_{obs}$) for re-oxidation of LPMO-Cu(I) by either $O_2$ or $H_2O_2$. Data describing reduction of LPMO-Cu(II) to LPMO-Cu(I) were fitted with a single exponential function ($y = a + b.e^{-k_{obs}.t}$) using the Kinetic Studio (Hi-Tech Scientific) software. For each experimental condition, all data were measured at least in triplicate and averaged. Plots of $k_{obs}$ versus AscA, $O_2$ and $H_2O_2$ concentrations were fit with linear least squares regression analysis to determine second order rate constants (using an in-house Matlab script). $O_2$ re-oxidation experiments performed for *Sm*AA10A-WT using the stopped-flow approach or the conventional fluorimeter gave equivalent results, both yielding second order rate constants of 3.3 $M^{-1}$ $s^{-1}$.

***Electron paramagnetic resonance spectroscopy***

EPR spectra were recorded using a BRUKER EleXsys 560 SuperX instrument equipped with an ER 4122 SHQE SuperX high sensitivity cavity. Spectra were recorded using 20 mW microwave power and 2 G modulation amplitude at room temperature. All samples were prepared with 0.2 mM spin trap probe CAT1H (1-Hydroxy-2,2,6,6-tetramethylpiperidin-4-yl-trimethylammonium chloride). The CAT1H stock solution was kept on ice, under an inert $N_2$ atmosphere, and the signal of CAT1H in buffer was measured at the beginning and the end of the experiment. Typically, all sample contents were mixed in less than 10 seconds, drawn into a 50 µL capillary (Brand gmbh) that subsequently was sealed by haematocrit (Brand gmbh), and measured exactly after 4 minutes. Spectra were imported into Matlab using Easyspin (Stoll&Schweiger_2006) and integrated using Matlab functions. Sample composition is shown with the results in **Figure S18**.

***Molecular dynamics simulations (classical only).*** The starting model for investigating the *Sm*AA10A reaction mechanism was taken from an experimentally informed enzyme-β-chitin model (~150,000 atoms, **Figure S4**) that had previously been equilibrated for 272 ns.[13] Force field parameters for $H_2O_2$ were derived using Paramfit[14] and Gaussian 09[15](**Table S1**) while the Cu(I)-histidine-brace force field parameters were taken from a previous study.[13] A water molecule ~3 Å away from the Cu-atom was replaced by a $H_2O_2$ molecule and the new model was equilibrated applying the following procedure. The model was first subjected to 2,500 steps of energy minimization with 5 kcal.mol$^{-1}$·Å$^{-2}$ positional restraints on all atoms but the introduced $H_2O_2$ molecule. Then, a 50 ns equilibration step was carried out in the NVT ensemble at 300 K using the weak coupling algorithm and a time constant of 10 ps to regulate the temperature. In this step, 2 kcal·mol$^{-1}$·Å$^{-2}$ positional restraints were applied to the C1 atoms of the lowest layer of NAG chains of the chitin model (the "highest" layer being the one interacting with the LPMO). Throughout the MD simulation, a one-sided harmonic potential of 5 kcal mol$^{-1}$ Å$^{-2}$ was applied to one of the $H_2O_2$ O-atoms to keep it between 2.0 and 3.5 Å from the Cu-atom. A time step of 2 fs, periodic boundary conditions with a 12 Å cutoff for non-bonded interactions, and particle mesh ewald treatment of long-range electrostatics were applied, while hydrogen atoms were constrained by the SHAKE algorithm.[16,17] All MD-simulations were conducted using the CUDA version of PEMEMD included in AMBER16.[18] Analysis of trajectories was performed using the *cpptraj* module included in AmberTools.[19]

***Molecular dynamics (MD) simulations (with QM-region).*** A snapshot of the *Sm*AA10A-Cu(I)-$H_2O_2$-β-chitin MD-trajectory with an active site geometry similar to what is observed in the crystal structure (PDB ID 2BEM:C)[1] and a Cu-H1 distance of 3.8 Å (the average distance observed in the ensemble) was selected from the 50 ns trajectory. The QM-region included the $H_2O_2$ molecule, Cu(I), the enzyme residues H28, H114, and E60, and two NAG units (see **Figure S5** for the definition of link atoms and QM- and MM-regions). The QM/MM interface of AMBER16[20] was utilized to execute the QM/MD simulations while ORCA[21] provided energies and gradients externally (UBP86/Def2-SVPP). The MD parameters listed in the previous section were also applied to the QM/MM/MD simulation, except that the time step was reduced to 0.5 fs. After a few hundred steps, the $H_2O_2$ O-O bond began to elongate (>0.2Å), and the restraint keeping $H_2O_2$ in the reaction cavity was immediately turned off after which the reaction continued to form an oxyl-intermediate. Another 5 kcal $mol^{-1}$ $Å^{-2}$ restraint was then applied to pull the oxyl O-atom closer to the substrate H1-atom that was approximately 2 Å away. When the Cu-oxyl bond started to elongate (>0.2Å), the restraint was turned off, after which H-abstraction and hydroxyl rebound happened spontaneously. The complete reaction is summarized in **Movie S1**.

***QM/MM calculations.*** The simulation in the previous section did not provide any information on the energetics of the reaction path, thus a QM/MM approach was chosen to estimate transition state energy barriers. A snapshot from the QM/MM/MD simulation taken before $H_2O_2$ reacted with Cu(I) was minimized using the same AMBER parameters as listed above and by applying the three step QM/MM minimization scheme described previously.[13] The minimized system, containing an intact $H_2O_2$ molecule ~3 Å from the copper ion, was truncated from ~150,000 to ~24 000 atoms, only keeping water, NAG units, and amino acid residues closer than 40 Å from the copper ion (**Figure 3A**). This initial truncated model was subjected to geometry optimization using ChemShell[22] in combination with ORCA, selecting an extended QM/MM-region (**Figure S5B**) and a large active-region (the part of the model that is allowed to move) that allowed the active site environment to relax (**Figure S5C**). This geometry optimization yielded the QM/MM starting model, named state **1**. In all further calculations a smaller active-region was employed (**Figure S5D**) to avoid potential non-relevant changes of hydrogen bonding patterns in the MM-region far from the enzyme active site. To generate models of the oxyl intermediate and final hydroxylated product, the initial positions of the corresponding $H_2O_2$ derived atoms were estimated from the QM/MM/MD simulation, replacing the $H_2O_2$ coordinates in state **1**, and then subjected to geometry optimization. Typically, all calculations were carried out utilizing the unrestricted functionals BP86[23,24] B3LYP[25] and TPSSh,[26] applying the Def2-SVP basis set for all atoms except the copper ion, which was described by Def2-TZVP. All final single point energies were calculated by B3LYP or TPSSh and Def2-TZVPP (see Results section for details). For all states of the system potentially containing unpaired electrons (i.e. all states except states **1** and **9**), calculations were carried out assuming open shell singlets, triplet states, or broken symmetry states. Dispersion was included through the Grimme's DFT-D3[27] approach with Becke-Johnson dampening,[28] and the RI[29] or RIJCOSX approximation[30] was used to speed up the calculations. The DL-find optimizer in ChemShell was applied to calculate nudged elastic band (NEB) minimum energy paths, carry out geometry optimizations to obtain energy minima, ad to apply the dimer method to locate transition states.[31,32] All transition states were subjected to frequency analysis to confirm a single imaginary frequency. Note that the frequency analysis of TPSSh transition state **6** resulted in two imaginary frequencies. Molecular coordinates and AMBER force field parameters were imported directly into ChemShell, securing consistency between the programs. The QM/MM electrostatic interactions were handled by ORCA, including charges from the MM-region in the QM-calculation. For calculations with $O_2$, the model named state **1** was taken as the initial model. The H-atoms of $H_2O_2$ were deleted and dummy AMBER parameters were generated for $O_2$. When imported into ChemShell, these dummy parameters were ignored.

***Free energy calculations of $H_2O_2$ diffusion into the reaction cavity.*** To investigate how accessible the confined reaction cavity is to small molecules such as $H_2O_2$, and to assess the energetics associated with the transport process, biased MD simulations (umbrella sampling) were carried out. Two start models were generated from the previously equilibrated *Sm*AA10A-Cu(I)-$H_2O_2$-β-chitin complex containing $H_2O_2$, one with $H_2O_2$ located at the entrance of the tunnel suggested by Bissaro et al.,[13] and one with $H_2O_2$ in the reaction cavity. A biasing potential of 5.0 kcal mol$^{-1}$ Å$^{-2}$ was applied in all simulations, and when necessary (see **Figure S23**) resampling was carried out using a biasing potential of 10.0 kcal mol$^{-1}$ Å$^{-2}$. The multistate Bennett acceptance ratio method[33] as implemented in PyMBAR was used to estimate the free energy associated with $H_2O_2$ diffusion in and out of the reaction cavity.

#### 2. Supplementary results

**Detailed analysis of the electronic structures of states 1-9.** The corresponding orbital transformation (COT) method was used to generate unrestricted corresponding orbitals (UCOs),[34] and the orbital overlaps calculated for UCOs indicate how the outer shell electrons interact. An UCO overlap value close to 1 indicates doubly occupied orbitals, values in the range ~0.85-0.05 indicate spin-coupled pairs within an orbital, and an UCO overlap equal to 0 indicates two orbitals, each occupied by one electron having parallel spins.[35] An example is state **2** that yields an UCO overlap of 0.82 for the HOMO, indicating a borderline case of doubly occupied orbitals and spin coupled pairs. However, the open shell singlet and broken symmetry approaches yield the same energy, indicating that the broken spin (BS) solution has collapsed to the closed shell state due to strong coupling between the two spins (J = 7940 $cm^{-1}$), a phenomenon previously described in the literature.[34] The UCO overlaps and the corresponding coupling constants are shown in **Table S2.**

**Modeling of H-abstraction from chitin by *Sm*AA10A copper-superoxide.** Even though previous computational studies have all found hydrogen abstraction by Cu-superoxide less feasible than with Cu-oxyl,[36–38] all these studies have utilized cello-oligosaccharides as model substrates. Thus, it was of interest to investigate the potential of superoxide to perform hydrogen abstraction on crystalline chitin substrate. The state **1** model from the $H_2O_2$ mechanism was used as a starting point, in which $H_2O_2$ was replaced by $O_2$. We probed several initial models where $O_2$ was placed at different places in the reaction cavity and they all resulted in the same final Cu-superoxide geometry (**Figure S9**). When comparing this complex with a *Sm*AA10A-Cu-$O_2$ complex that was geometry optimized in vacuum without substrate or solvent, it became apparent that the presence of substrate induces a different conformation of $O_2$ as a result of steric clashes with the chitin substrate (**Figure S9A)**. This difference in conformation does not appear to change the nature of the molecular bonding orbital of the *Sm*AA10A-Cu-$O_2$ complex (**Figure S9A)**. Compared to H-abstraction by a Cu-oxyl species (barrier of 10.6 kcal/mol, TPSSh functional), the corresponding reaction with Cu-superoxide is unfavorable (35 kcal/mol transition state energy barrier, TPSSh functional) (**Figure S9D)**. Thus, provided that $O_2$ can access the active site cavity and form a Cu-superoxide complex, we conclude that H-abstraction by superoxide is thermodynamically plausible by 16 kcal/mol, but kinetically very unlikely.

***$H_2O_2$ access to the active site.*** We have previously revealed the existence of a water tunnel, formed in the enzyme-polysaccharide complex interspace, connecting bulk solvent to the monocopper active site.[13] Here, we investigated the energetics associated with $H_2O_2$ diffusion into the active site by MD simulations combined with umbrella sampling, estimating free energy profiles of $H_2O_2$ moving in this interspace. Different from our previous work,[13] where the tunnel and its accessibility were assessed primarily using steric criteria , the methodology applied here is founded on assesing non-covalent interactions between $H_2O_2$ and the enzyme-substrate complex in a dynamic environment. During the course of the simulations, Glu60 and Asn185 appeared to play a gating role, restricting $H_2O_2$ access to the active site cavity (see **Figures S1, S21** and **movie S2**). Asn185 points away from the copper ion and do not swing into the active site cavity like Glu60. While it is apparent from our simulations that $H_2O_2$ can move around freely within the reaction cavity, one main entrance route stood out as the most feasible. This preferred route is indicated by a blue tunnel in **Figure S1A** and described by the blue potential of mean force (PMF) curve in **Figure S1B**. The free energy profile was obtained by translocating $H_2O_2$ into the reaction cavity from the bulk solvent. The second tunnel shown in **Figure S1** (red tunnel, red PMF) indicates a dead end, where the $H_2O_2$ molecule is restricted from escaping into bulk solvent by residues Asp182 and Ser115 and the protein main chain. The free energy barrier associated with diffusion of $H_2O_2$ into the reaction cavity from the bulk solvent was estimated to be less than 2 kcal/mol, indicating that this process is far from rate limiting.

### 3. Supplementary discussion

##### On the positioning of $H_2O_2$ in AA9 LPMOs

The second-sphere residues Gln162 and His147 in *Ls*AA9A (**Figure S26**), highly conserved within the AA9 family,[39] could potentially play a role similar to that of Glu60 in *Sm*AA10A, which is conserved in all AA10 C1-oxidizers. The equivalent histidine in another AA9 has been shown to be important for enzyme catalysis.[40] Surprisingly, His147 was not included in the QM-region by Wang et al.,[41] and we speculate that, depending on the protonation state of its side chain, His147 may either hydrogen bond (as proposed by Hedegård and Ryde)[42] with the distal O in $H_2O_2$ or possibly bind $H_2O_2$ in a strained conformation resembling state **1** in this work (**Figure S26**).

##### On the interpretation of the LPMO fluorescence signal

We previously reported the change in fluorescence upon addition of reductant and oxidant to *Sm*AA10A.[12] Here, we detailed the kinetics of both reduction and re-oxidation steps under single-turnover conditions. While the reduction step is rather straightforward to analyze and translates into an increase in fluorescence upon formation of the LPMO-Cu(I) species, the re-oxidation steps entails more complex reactions and we would like to detail our interpretation of data and openly address some unanswered questions that warrant further investigations.

When LPMO-Cu(I) reacts with $H_2O_2$ in the absence of substrate, one could expect the formation of Fenton reaction-like products, namely hydroxyl radicals and hydroxide ions. In agreement, we have previously shown that the copper-coordinating histidines are heavily oxidized when an LPMO is exposed to excess of reductant and $H_2O_2$ in the absence of substrate, suggesting the formation of oxidizing species that do damage within a diffusion-limited range. Under such conditions, total enzyme inactivation is observed.[5,43,44] Here, we show by spin-trap experiments that hydroxyl radicals are indeed formed (**Figure S17**). Nonetheless, it is important to note that the formation of such hydroxyl radical under single turnover conditions leads to very little loss of activity of *Sm*AA10A (**Figure S19**), indicating that multiple turnover in absence of substrate is required to modify the enzyme active site or active site surroundings so that inactivation is achieved.

In presence of substrate, a faster decay of the fluorescence was observed, which is consistent with a substrate-assisted initial step of Cu(I) re-oxidation by $H_2O_2$. Also, the suggested peroxygenase mechanism predicts that at the end of the catalytic cycle the Cu(I) state should be regenerated. Therefore, in an ideal experiment, reacting for instance 1 umole of SmAA10A-Cu(I) with 0.5 umole of H2O2 in presence of substrate should yield 1 umole of SmAA10A-Cu(I) at the end of the reaction. In facts, 0.8 umoles were detected. If SmAA10A-Cu(I) was not regenerated, only 0.5 umoles would have been measured. This result is therefore in support of the regeneration of the Cu(I) state, and of the proposed mechanism more generally. However, it is also clear that not 100% of SmAA10A-Cu(I) was regenerated, and this became even more true when increasing the amount of H2O2 (Figure S16). It is therefore plausible that Cu(I) is re-generated but that it is not observed because it is rapidly re-oxidized at the end of the catalytic cycle when an excess of $H_2O_2$ is employed (**Figure S19**). Slow dissociation of the oxidized $NAG_6$ and the regenerated LPMO-Cu(I) state may also favour re-oxidation by a new $H_2O_2$ molecule before the enzyme can enter a new catalytic cycle. Regardless of these “downstream” complications, the fact that substrate accelerates the reaction between Cu(I) and $H_2O_2$ is clearly established here (**Figure 6**).

**On relating experimental constant rates, energy barriers and electron transfer rates**

Attempts to relate energy barriers derived from QM/MM studies to experimental apparent catalytic constants should be considered with caution. Notably, setting the pre-exponential factor (A) in the Arrhenius equation ($k_{\text{cat}} = A.e^{\frac{-E_{TS}}{R.T}}$) is complicated since this factor is meant to reflect productive collisions between reactants, a parameter likely to be greatly affected by the confined nature of the active site when the enzyme is in complex with the polysaccharide. This point is well illustrated by the constrained diffusion of $H_2O_2$ towards the active site through a tunnel access. A common value of $A = 10^{12}$ has been selected in recent LPMO work.[41,42] Using a supposedly similar value (deduced by us, not stated by the authors), Wang et al. have concluded that the rate constant of 0.11 $s^{-1}$ reported by Frandsen et al.[45], and determined under ambient $O_2$ (i.e. without control of potential $H_2O_2$ levels) would correspond to a barrier of approximately 18 kcal/mol, which is much higher than the highest energy barrier coming out of their calculations on $H_2O_2$-driven catalysis (6.9 kcal/mol).[41] This difference led the authors of that study to suggest that the rate-determining event of the apparent LPMO rate is a step not intrinsic to the LPMO catalytic cycle, namely the hydrolysis of the hydroxylated product. Following the same logic (with $A = 10^{12}$), we calculate that an energy barrier of 6.9 kcal/mol would translate into a rate constant > $10^7$ $s^{-1}$ for the H-abstraction reaction, which seems unlikely. Importantly, it has been claimed that in standard reaction conditions, where $H_2O_2$ is generated *in situ* at low, non-harmful concentrations, the rate of $H_2O_2$ generation determines the overall LPMO apparent rate.[5,44,46] Therefore, apparent LPMO rates, as in Frandsen et al., 2011, may reflect the probability of encounters between $H_2O_2$ and a reduced substrate-bound LPMO rather than any intrinsic reaction of the LPMO cycle, which complicates the comparison of experimental and theoretical rates. $H_2O_2$-driven catalysis by *Sm*AA10A yielded a $k_{\text{cat}}$ of 6.7 $s^{-1}$ (at 25 ºC) on chitin nanowhiskers [44], which can be translated into an energy barrier of ~13.0 kcal/mol (i.e. the highest energy barrier calculated here between states **5** to **6** (**Figure 3D**) only when considering a collision factor 40-fold lower (i.e. $A = 0.23 \times 10^{11}$) than the A value used by others.

In the frame of the Marcus theory, we can estimate the outer sphere electron transfer rate, knowing the reduction potentials of the half-oxidation *Sm*AA10A-Cu(I) → Cu(II) and half-reduction $O_2$ → $O_2^{\bullet-}$ (see calculations below). Regarding re-organization energies, we followed the approach outlined by Kjaergaard et al. [47]:

Marcus´ theory equation: $k_{ET}=Z.e^{\frac{-\Delta G^{\neq}}{R.T}}$

*Where* Z (frequency factor) = $10^{11}$ $M^{-1}$ $s^{-1}$ (according to Kjaergaard et al.)[47]

$\Delta G^{\neq} = (\lambda + \Delta G°)^2/ 4\lambda$

R = 8.3415 $J.K^{-1}.mol^{-1}$

T = 298.15 (K) (experiments performed at 25 °C)

$\lambda$ (total reorganization energy) = $(\lambda_{Cu\text{-}SmAA10A,\ donor} + \lambda_{O2,\ acceptor})/2$ = 1.74eV,
where $\lambda_{O2,\ acceptor}$ =1.89 eV (experimentally determined, taken from Kjaergaard et al.)
and $\lambda_{Cu\text{-}SmAA10A,\ donor}$ = 1.58 eV (taken from Kjaergaard et al.; note that experimentally determined value for LPMO-Cu is lacking so we use the reorganization energy for a Cu(I/II)-complex with a (tris(2-pyridyl-methyl)amine ligand as best approximation).

$\Delta G° = -nF\Delta E°$ = 65,467 J/mol (eq. 15.7 kcal/mol)

*Where* $\Delta E°_{(with\ O2)}$ = E° ($O_2/O_2^{\bullet-}$) - E° (Cu(I/II)-*Sm*AA10A) = **- 0.440 V**

n = 1

F = 96485 C/moles

*With*

E° (Cu(I/II)-*Sm*AA10A) = 275 ± 6 mV *vs* NHE, determined at pH 6.0[48]

E° ($O_2/O_2^{\bullet-}$) = -165 mV *vs* NHE, at pH 7.0.[49]

With these parameters, Marcus theory predicts a $\boldsymbol{k_{ET,\,O2}}$ of **3.39 $M^{-1}$ $s^{-1}$**, a value that is remarkably similar to the second order rate constant resulting from the experiments described here, $\boldsymbol{k_{O2}}$ **= 3.33 $M^{-1}$ $s^{-1}$**.

### 4. List of supplementary figures

**Figure S1.** Diffusion of $H_2O_2$ from bulk solvent to the copper active site of *Sm*AA10A in complex with β-chitin.
**Figure S2.** Comparison of $H_2O_2$ consumption by *Sm*AA10A wildtype in presence of polysaccharide β-chitin or soluble oligosaccharide.
**Figure S3.** Structural model of Tma12 and phylogenetic location within the LPMO superfamily.
**Figure S4.** The equilibrated *Sm*AA10A-Cu(I)-β-chitin complex.
**Figure S5.** QM-region and active-region definitions.
**Figure S6.** Screening of reaction conditions to monitor $H_2O_2$ consumption by *Sm*AA10A.
**Figure S7.** Underestimation of $H_2O_2$ levels due to competing AscA-peroxidase activity of HRP.
**Figure S8.** Snapshots along the reaction coordinate of the 9 main reaction states and corresponding relative energies.
**Figure S9.** QM/MM models of $O_2$ bonding by *Sm*AA10A and H-abstraction by Cu-superoxide.
**Figure S10.** Illustration of LPMO reductive and oxidative half-reactions and expected related change in intrinsic fluorescence.
**Figure S11.** Relationship between fluorescence signal and concentration of *Sm*AA10A-Cu(I) and Cu(II).
**Figure S12.** Kinetics of the reduction of *Sm*AA10A-Cu(II) to Cu(I) in presence of soluble substrate $NAG_6$.
**Figure S13.** Fitting of $O_2$-mediated *Sm*AA10A-Cu(I) re-oxidation data, acquired at low $O_2$ concentration (47 μM), using several simple models.
**Figure S14.** Fitting of $O_2$-mediated *Sm*AA10A-Cu(I) re-oxidation data, acquired at high $O_2$ concentration (578 μM), using several simple models.
**Figure S15.** Fitting of $O_2$-mediated *Sm*AA10A-Cu(I) re-oxidation data by a model describing the dimer hypothesis.
**Figure S16.** Fitting $H_2O_2$-mediated *Sm*AA10A-Cu(I) re-oxidation data by single exponential model.
**Figure S17.** Titration of *Sm*AA10A-Cu(I) with $H_2O_2$.
**Figure S18.** Spin-trapping of reaction products generated when incubating *Sm*AA10A-Cu(I) with $H_2O_2$.
**Figure S19.** Monitoring *Sm*AA10A-Cu(I) decay upon aging with $H_2O_2$ by measuring the residual oxidative activity.
**Figure S20.** Residual activity of *Sm*AA10A after treatment of *Sm*AA10A-Cu(I) wild-type with different $H_2O_2$ concentrations.
**Figure S21.** $NAG_6$ is a suitable substrate for single turnover conditions.
**Figure S22.** Re-oxidation of *Sm*AA10A-Cu(I) by $H_2O_2$ in presence (0.5 mM) and absence of $NAG_6$.
**Figure S23.** Quality evaluation of umbrella-sampling data.
**Figure S24.** Time-courses for binding of *Sm*AA10A wild-type and mutants to β-chitin.
**Figure S25.** Reactivity between $H_2O_2$ and *Sm*AA10A-Cu(I) wild-type and mutants.
**Figure S26.** Hydrogen bonding patterns involving $H_2O_2$ in the active site cavity of *Ls*AA9A.

**Scheme S1.** Proposed reaction scheme of *Sm*AA10A-Cu(I) with $O_2$ involving LPMO dimerization.

**Table S1.** AMBER force field parameters for $H_2O_2$.
**Table S2**. Population and spin parameters along the calculated reaction pathway,
**Table S3**. Second order rate constants of the re-oxidation by $H_2O_2$ of *Sm*AA10A-Cu(I) wild-type and mutants thereof.
**Table S4.** PCR primers used for site-directed mutagenesis of *Sm*AA10A.

**Movie S1.** QM/MM simulation of the proposed chitinolytic peroxygenase mechanism.
**Movie S2.** Diffusion of $H_2O_2$ through the tunnel access from bulk solvent to the copper active site of *Sm*AA10A in complex with β-chitin.

#### 5. Supplementary figures

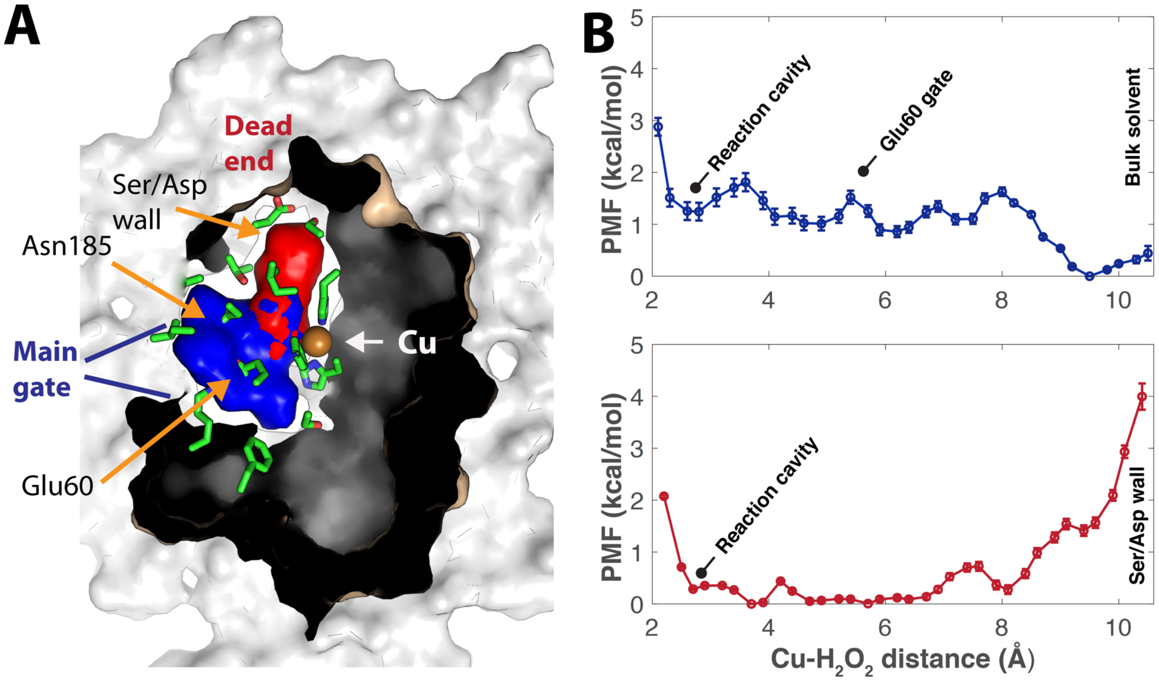


**Figure S1. Diffusion of $H_2O_2$ from bulk solvent to the copper active site of *Sm*AA10A in complex with β-chitin.** Panel **A** shows a top-down view of the complex. The blue (pulling $H_2O_2$ into the reaction cavity from bulk solvent) and red (pushing $H_2O_2$ out of the reaction cavity) tunnels overlap in the reaction cavity and extend towards bulk solvent and a dead end, respectively. Amino acid side chains lining the two tunnels are displayed in green. The crystalline chitin surface is light grey. The corresponding free energy (potential of mean force, PMF) profiles of pulling (blue line) or pushing (red line) $H_2O_2$ are shown in panel **B**. The position of the reaction cavity, bulk solvent, gating residues (Glu60/Asn185) and residues blocking the red tunnel are indicated along the profiles. Quality evaluation of the umbrella sampling data is provided in **Figure S23**.

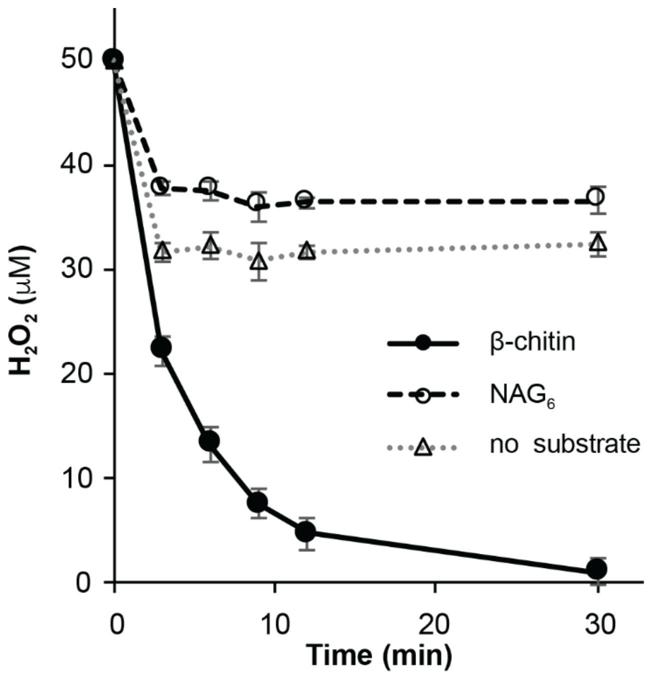


**Figure S2. Comparison of $H_2O_2$ consumption (50 µM at $t_0$) by *Sm*AA10A wildtype (50 nM) in presence of polysaccharide (β-chitin; 10 g.$L^{-1}$; solid line) or a soluble oligosaccharide ($NAG_6$; 2 mM; dotted line).** Reactions were initiated by addition of AscA (20 µM). Note that in the "no substrate" reaction the observed consumption of $H_2O_2$ is due to the AscA-peroxidase activity of HRP (see **Figure S6**). In the $NAG_6$ reaction, a few turnovers are predicted to occur (see **Figure S20**) whereas consumption of $H_2O_2$ due to the AscA-peroxidase activity of HRP is reduced due to AscA consumption during the LPMO reaction. Reactions were carried out in sodium phosphate buffer (50 mM, pH 7.0) at 40 °C in a thermomixer (1000 rpm). Error bars show ± s.d. (n = 3, independent experiments).

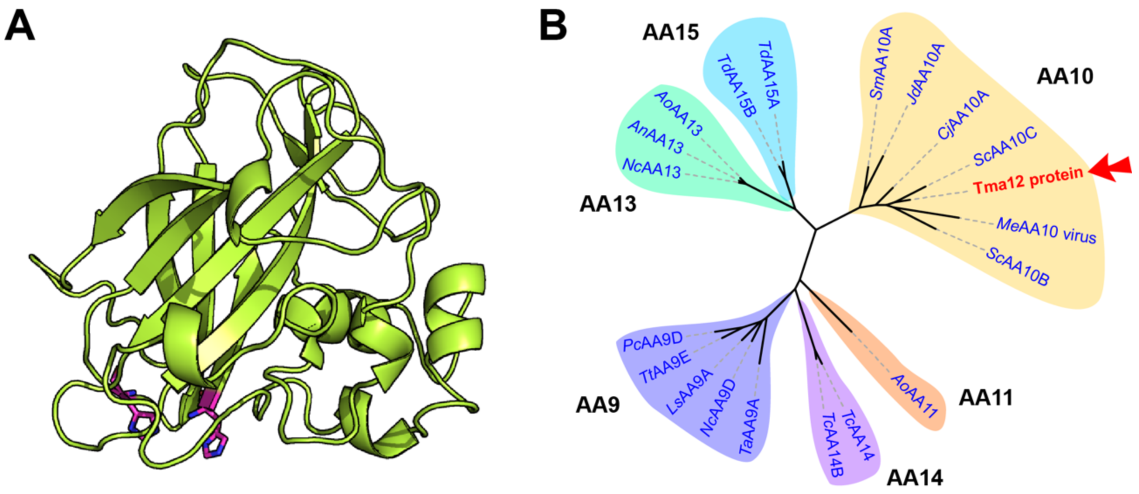


**Figure S3. (A) Structural model of Tma12 and (B) phylogenetic location of Tma12 within the LPMO superfamily**. **(A)** Phyre2 was used to predict a structural model (shown as green cartoon) of the mature chitin-binding protein from *Tectaria macrodonta* (Tma12; GenBank ID AFR32946.1). 96% of residues were modelled with > 90% confidence, according to Phyre2 criteria. The two conserved and surface-exposed histidines coordinating the copper atom in functional LPMOs are shown as magenta sticks. **(B)** Phylogenetic tree of Tma12 and 19 characterized LPMOs spanning all families hitherto reported (i.e. AA9 to 11 and 13 to 15). On the basis of a previous analysis of the structural diversity of LPMOs [39] we selected a set of LPMOs sequences that represent various structural clusters. The selected sequences include the following: for AA9s, AA9D from *Phanerochaete chrysosporium* (*Pc*AA9D, PDB 4B5Q),[50] AA9D from *Neurospora crassa* (*Nc*AA9D, PDB 4EIR),[51] AA9A from *Lentinus similis* (*Ls*AA9A, PDB 5ACF),[45] AA9A from *Thermoascus aurantiacus* (*Ta*AA9A, PDB 2YET),[7] AA9E from *Thielavia terretris* (*Tt*AA9E, PDB 3EII);[52] for AA10s, AA10A from *Serratia marcescens* (*Sm*AA10A, PDB 2BEM),[1] AA10B and AA10C from *Streptomyces coelicolor* (*Sc*AA10B, PDB 4OY6 and *Sc*AA10C, PDB 4OY7),[53] AA10A from *Cellvibrio japonicus* (*Cj*AA10A, PDB 5FJQ),[54] AA10A from *Jonesia denitrificans* (*Jd*AA10A, PDB 5AA7),[55] insect poxvirus fusolin (*Me*AA10 virus, PDB 4OW5);[56] for AA11s, the only hitherto characterized AA11, from *Aspergillus orizae* (*Ao*AA11, PDB 4MAI);[57] for AA13s, AA13 from *Aspergillus orizae* (*Ao*AA13, PDB 4OPB), AA13 from *Aspergillus nidulans* (*An*AA13, Uniprot ID Q5B1W7), AA13 from *Neurospora crassa* (*Nc*AA13, Uniprot ID Q7SCE9);[8] for AA14s, AA14A and AA14B from *Trametes coccinea* (*Tc*AA14A, GenBank ID AUM86166.1, *Tc*AA14B, PDB 5NO7);[58] for AA15s, AA15A and AA15B from *Thermobia domestica* (*Td*AA15A, PDB 5MSZ, *Td*AA15B, GenBank ID SIW61373.2).[59]

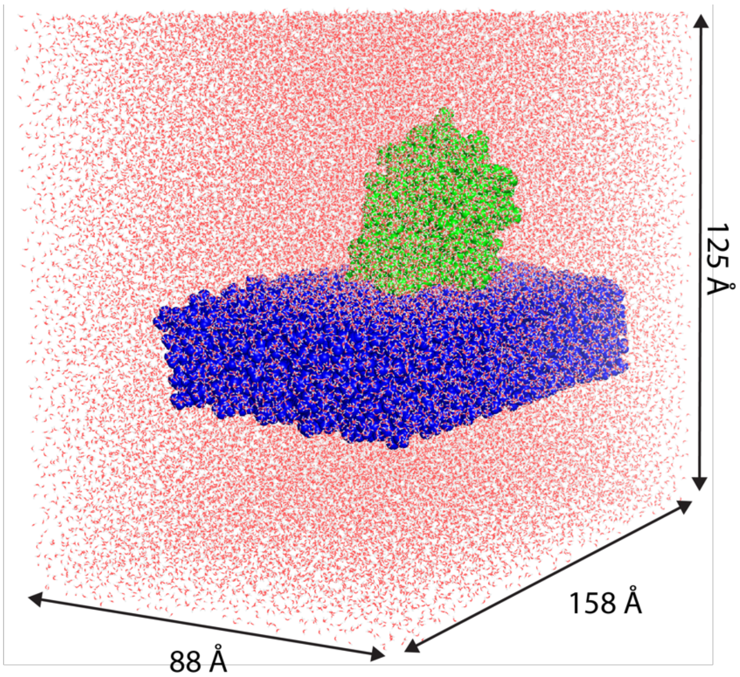


**Figure S4. The equilibrated *Sm*AA10A-Cu(I)-β-chitin complex**. The complex, consisting of ~150,000 atoms, was initially assembled in an experiment-guided approach.[13] *Sm*AA10A and **β**-chitin are shown with green and blue surface, respectively. Water molecules are shown as white and red sticks.

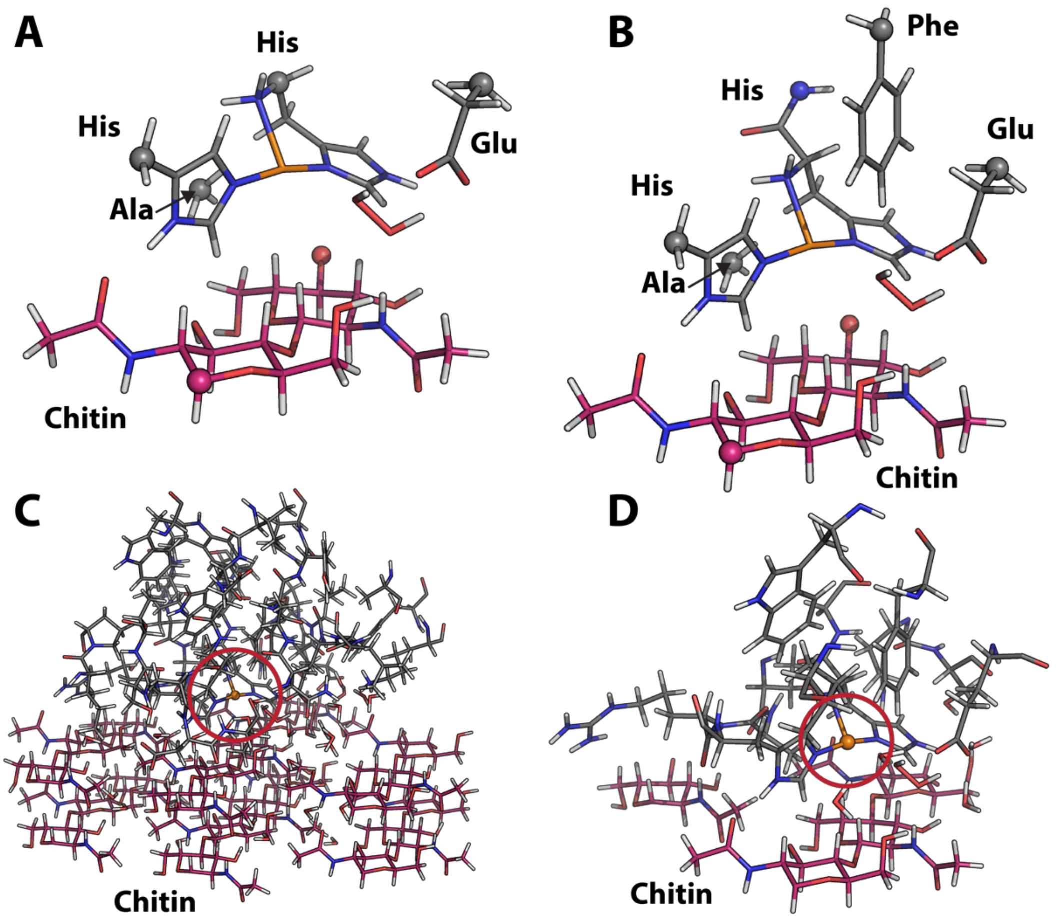


**Figure S5. QM-region and active-region definitions.** **(A)** QM-region of the QM/MD simulation (AMBER/ORCA) and **(B)** QM-region of the QM/MM calculations (ChemShell/ORCA). The link-atoms are connected to the C, N or O-atoms displayed as spheres in panel **A** and **B**. **(C)** The (large) part of the LPMO-chitin complex that was allowed to relax in the initial ChemShell QM/MM geometry optimization. **(D)** The smaller active-region that was allowed to move in subsequent QM/MM calculations. Residues marked as Ala, His, Phe, and Glu are Ala112, His28 (N-terminus) and His114, Phe187, and Glu60.

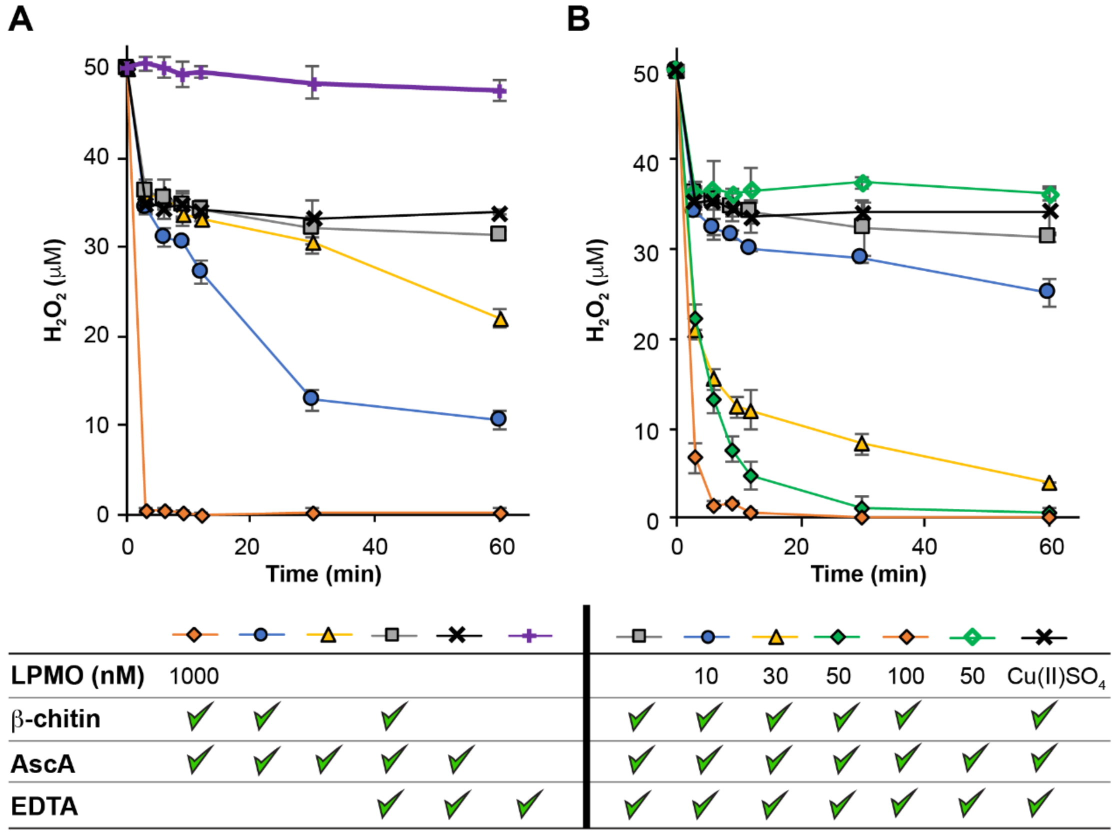


**Figure S6. Screening of reaction conditions to monitor $H_2O_2$ consumption by *Sm*AA10A. (A)** Assessment of the impact of reaction components on $H_2O_2$ consumption and **(B)** screening of the enzyme concentration. All reactions were carried out in sodium phosphate buffer (50 mM, pH 7.0) at 40 °C in a thermomixer (1000 rpm) and contained $H_2O_2$ (50 µM initial concentration). As indicated below the graphs (green ticks = presence in mixture), reactions varied in terms of the presence of *Sm*AA10A (in nM; no number means no enzyme), β-chitin (10 $g.L^{-1}$), EDTA (50 µM) and AscA (20 µM). The reactions were initiated by addition of ascorbic acid. Error bars show ± s.d. (n = 3, independent experiments).

Reactions shown in **panel A** show that using 1000 nM of enzyme (orange squares) leads to extremely fast $H_2O_2$ consumption and is thus not a suitable condition for monitoring purposes. We also see that in absence of enzyme a significant background reaction between AscA, $H_2O_2$ and other compounds contained in β-chitin (likely free metals) occurs (blue circles). Removing β-chitin from the mixture reduces non-enzymatic $H_2O_2$ consumption (yellow triangles). The addition of EDTA leads to a more moderate non-enzymatic background reaction in presence of β-chitin (grey squares), confirming the presence of free metals in the β-chitin suspension. This background reaction cannot be further minimized since a similar profile is obtained in reactions with only AscA and EDTA (black crosses). This "minimal" background $H_2O_2$ consumption is thus mainly due to the presence of AscA since $H_2O_2$ in presence of only EDTA is relatively stable (purple crosses). We show in **Figure S7** that this "background" is related to the competing AscA-peroxidase activity of HRP leading to underestimation of $H_2O_2$ levels. We decided to use a concentration of AscA of 20 µM, as a compromise between efficient enzyme reduction and not too high background reactions.

**Panel B** shows enzyme concentration-dependent consumption of $H_2O_2$ (from 0 to 100 nM of *Sm*AA10A), which is consistent with $H_2O_2$ consumption being an enzyme-catalyzed reaction. 50 nM of enzyme allows to reach complete consumption of $H_2O_2$ while initial time points are still measurable (filled green diamonds). Therefore, 50 nM was chosen as a working enzyme concentration for further experiments. A control reaction in absence of β-chitin and presence of 50 nM of *Sm*AA10A leads to a background-like $H_2O_2$ consumption profile (empty green diamonds). A similar profile was observed when the enzyme was replaced by 50 nM of Cu(II)$SO_4$ (black crosses). Error bars show ± s.d. (n = 3, independent experiments).

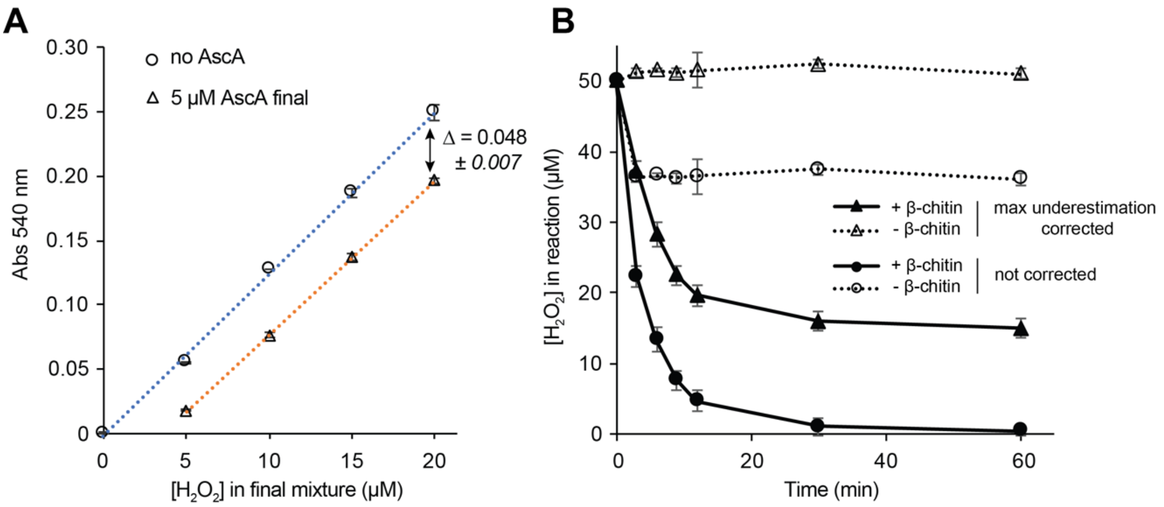


**Figure S7. Underestimation of $H_2O_2$ levels due to competing AscA-peroxidase activity of HRP. (A)** Effect of residual AscA on $H_2O_2$ standard curve as measured by absorbance of resorufin at 540 nm. The AmplexRed/HRP assay is based on the oxidation of AmplexRed (colorless) to yield resorufin (pink color, absorbing at 540 nm) by HRP using $H_2O_2$ as oxidant with a 1:1 stoichiometry. In all experiments related to $H_2O_2$ consumption 20 µM of AscA was employed in the reaction mixture. As a standard procedure, 25 µL of reaction mixture was mixed with 75 µL of a pre-mix of HRP (100 nM final), AmplexRed (100 µM final) and sodium phosphate buffer (50 mM, pH 7.0), resulting in a final maximum concentration of AscA of 5 µM (assuming no consumption of AscA in the preceding LPMO reaction, which. Obviously, is not true and wil vary). In panel A, 25 µL of a solution of $H_2O_2$ (0-80 µM) was mixed with 75 µL of a premix of HRP/AmplexRed/buffer with or without AscA (5 µM final concentration). The graphs show the absorbance measured at 540 nm. One can observe that when residual AscA is present a lower absorbance is measured, due to the AscA-peroxidase activity of HRP scavenging a fraction of the $H_2O_2$.[60] Therefore, in reactions containing AscA a fraction of $H_2O_2$ will not translate into the absorbance measured at 540 nm, leading to an underestimation of the actual levels of $H_2O_2$. **(B)** The graph shows the time-course of $H_2O_2$ conversion in reactions containing *Sm*AA10A (50 nM) in the absence (empty circles) or presence (solid circles) of β-chitin (10 $g.L^{-1}$), in reactions containing 50 µM $H_2O_2$ and 20 µM AscA at $t_0$ (reproduced from **Figure 2B** in the main article). The absorbance is at maximum underestimated by 0.048 AU (see panel A) if 20 µM of AscA from the initial reaction (i.e. 5 µM in final assay) are carried over into the AmplexRed/HRP assay. Correcting the raw absorbance data by the maximum underestimation values yields the data points shown as solid triangles (with β-chitin) and empty triangles (without β-chitin). The rapid decrease in the $H_2O_2$ concentarion observed in the reaction without chitin (empty circles), i.e. a condition in which AscA is not consumed significantly, is mainly due to AscA carry over. Thus, we provide here an explanation for the abrupt drop in the $H_2O_2$ concentration and also point out limitations associated with the assay. Error bars show ± s.d. (n = 3, independent experiments).

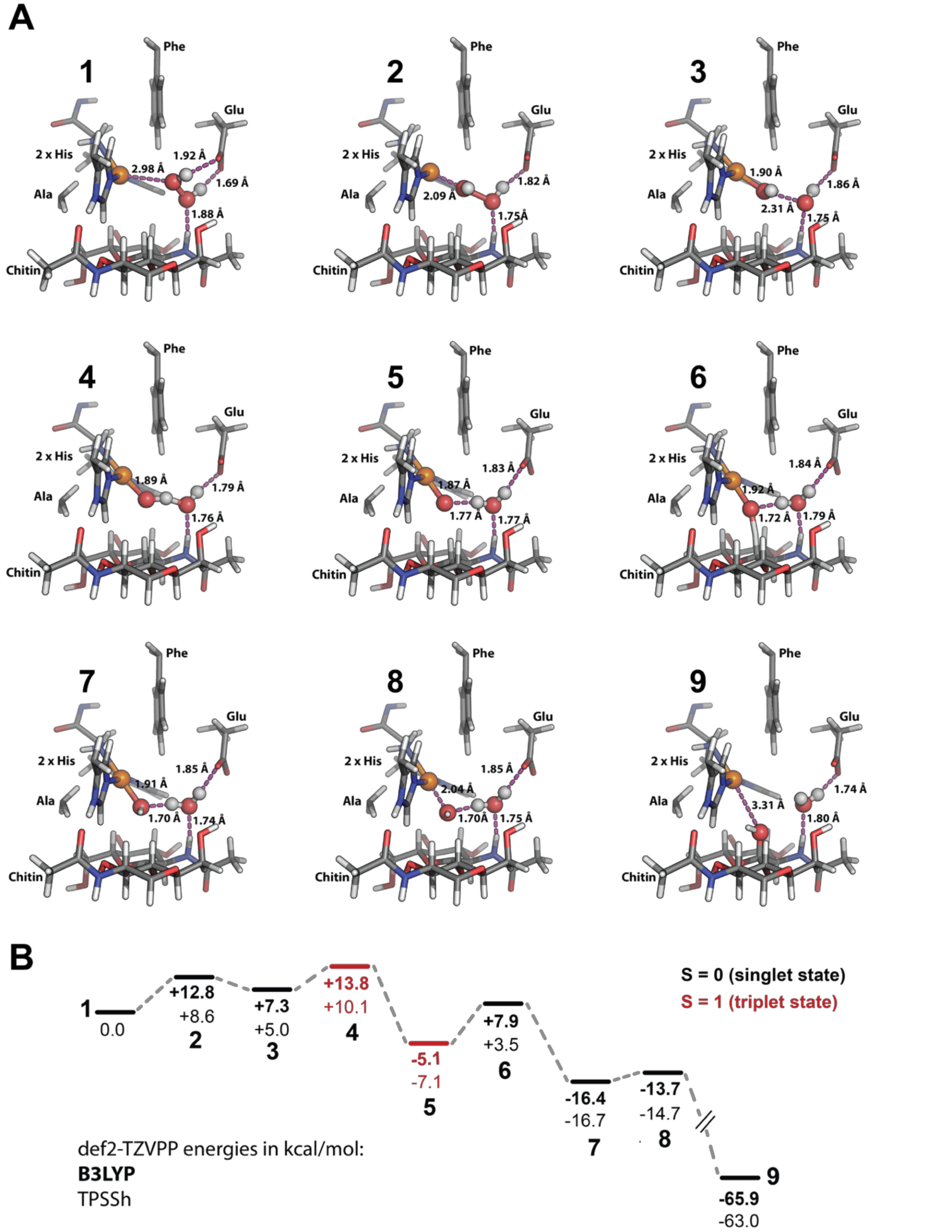


**Figure S8. (A) Snapshots along the reaction coordinate represented by 9 main reaction states and (B) corresponding relative energies. (A)** For each reaction state (**1** to **9**), the hydrogen bonding network involving $H_2O_2$ and derived species is show by pink dotted lines. Residues marked as Ala, 2x His, Phe, and Glu are Ala112, His28 (N-terminus), His114, Phe187 and Glu60. Panel B is a reproduction of panel D in **Figure 3** of the main manuscript. For the sake of clarity the MM-region is not shown.

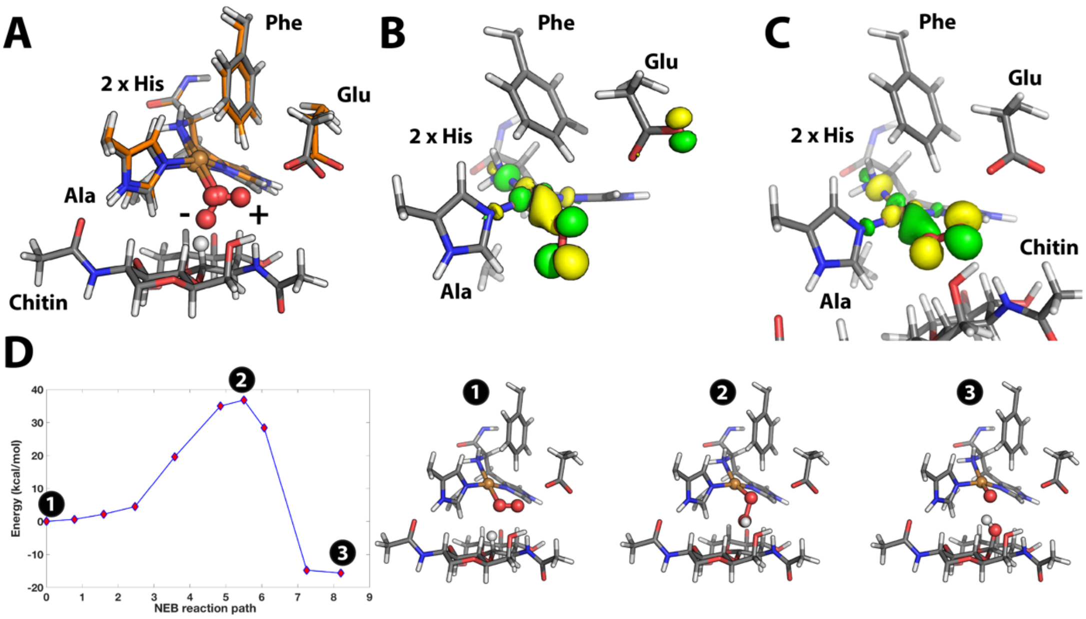


**Figure S9. QM/MM models of $O_2$ bonding by *Sm*AA10A and H-abstraction by Cu-superoxide. (A)** Superimposed models of $O_2$ binding to the active site Cu in the presence (grey carbons) or absence (orange carbons) of chitin. In absence of substrate, the $O_2$ moiety adopts a downward orientation (indicated by "-" sign in panel A) whereas it lies parallel to the substrate plane in the presence of chitin (indicated by "+"). This indicates a strained conformation for $O_2$ in the presence of substrate. The quasi restricted orbital describing binding of $O_2$ is shown for the models **(B)** without chitin substrate and **(C)** with chitin substrate. **(D)** Energy landscape of H-abstraction by Cu-superoxide, indicating a transition barrier of ~35 kcal/mol. The starting (1), converged climbing image (transition state) (2) and product (3) geometries are provided in panel **D**. For the sake of clarity the MM-region is not shown. Residues marked as Ala, His, Phe, and Glu are Ala112, His28 (N-terminus), and His114, Phe187 and Glu60.

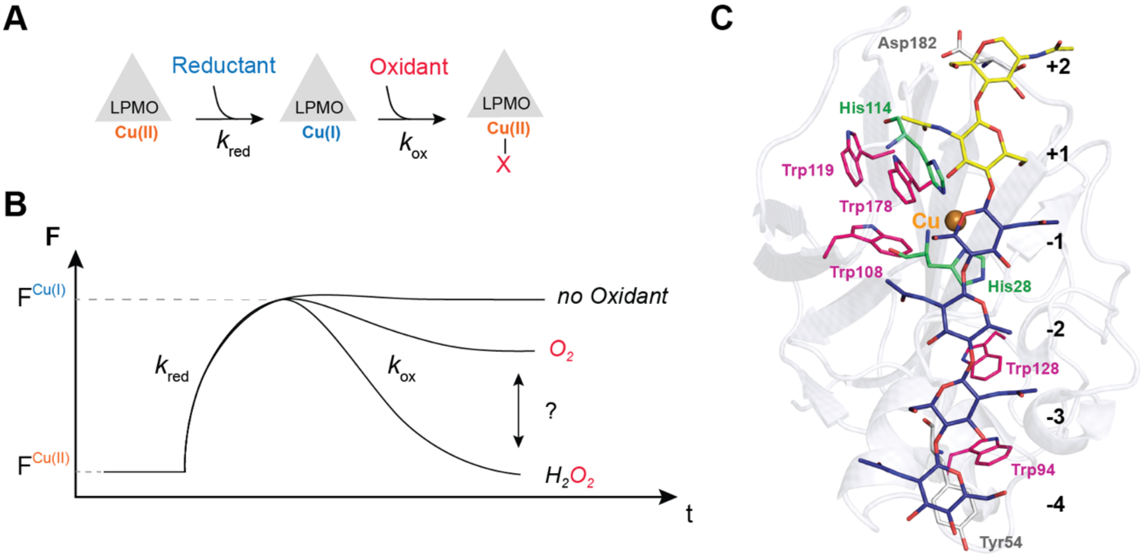


**Figure S10. Illustration of LPMO reductive and oxidative half-reactions and expected related changes in intrinsic fluorescence. (A-B)** Upon reduction, the intrinsic fluorescence (F) of the LPMO increases rapidly and then decreases slowly back to its ground state due to a putative re-oxidation by an oxidant (e.g. $O_2$ or $H_2O_2$).[12] The rates of the increase ($k_{red}$) and decrease ($k_{ox}$) in fluorescence are expected to reflect the efficiency of the reduction and re-oxidation steps, respectively. Such rates were determined in the present study under single turnover conditions. **(C)** Top-view of a predicted model of *Sm*AA10A in complex with $NAG_6$[13] showing the location of the histidines (green sticks) coordinating the single copper atom (orange sphere) and the five Trp residues (magenta sticks) responsible for the fluorescence. The figure shows the experimentally determined principal binding mode, where $NAG_6$ occupies subsites -4 to +2, leading to a product profile where $A4^{ox}$ (i.e. C1-oxidized chitotetraose) is the main oxidized product.[13]

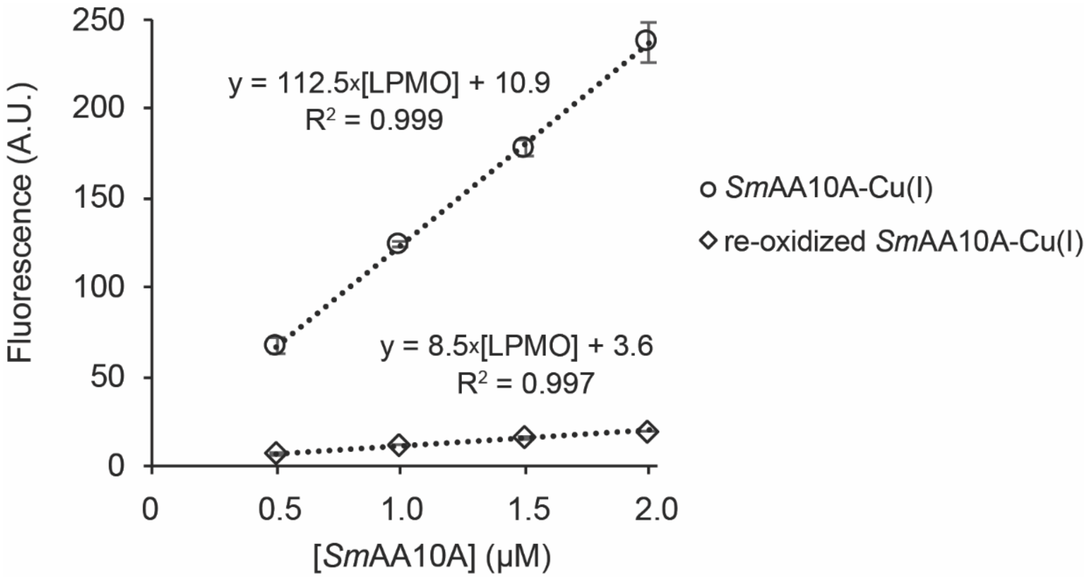


**Figure S11. Relationship between the fluorescence signal and the concentration of *Sm*AA10A-Cu(I) and *Sm*AA10A-Cu(II).** *Sm*AA10A-Cu(I), at dfferent concentrationsm was prepared anaerobically in a screw-cap quartz cuvette and the fluorescence signal was measured (excitation 280 nm, emission 340 nm). Then, $H_2O_2$ (4 µM final concentration) was added to each preparation to obtain the fluorescence signal of *Sm*AA10A-Cu(II) (i.e. fully re-oxidized *Sm*AA10A-Cu(I), see **Figure S17**). All reactions were performed at 25 °C in sodium phosphate buffer (50 mM, pH 7.0). Error bars show ± s.d. (n = 3, independent experiments).

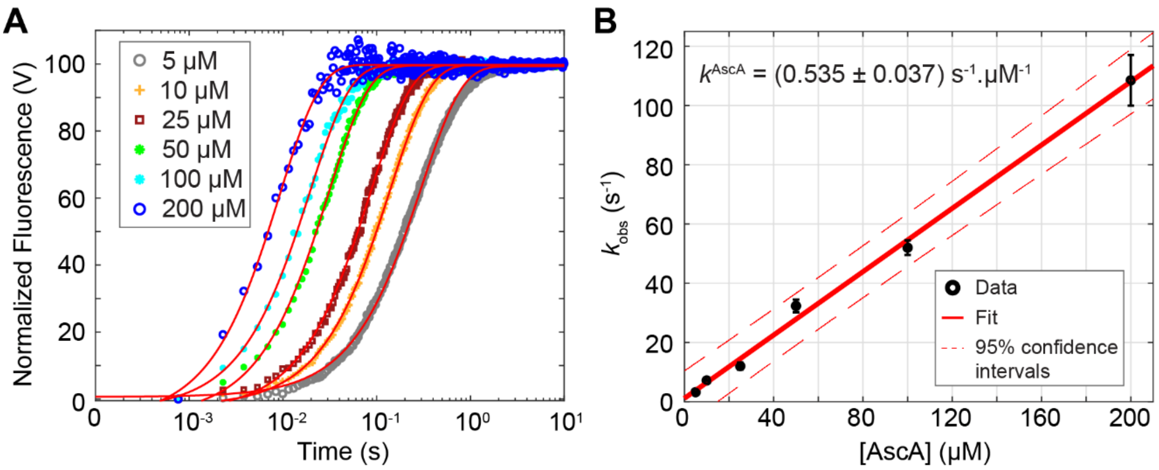


**Figure S12. Kinetics of the reduction of *Sm*AA10A-Cu(II) to *Sm*AA10A-Cu(I) in the presence of soluble substrate, NAG$_6$. (A)** Anaerobic solutions of *Sm*AA10A-Cu(II) (final concentration, 5 µM) and NAG$_6$ (1 mM final) were anaerobically mixed with varying concentrations of AscA and the changes in fluorescence associated with reduction of the Cu(II) were monitored as a function of time. The reactions were carried out in potassium phosphate buffer (50 mM, pH 7.1) at 25 °C. The final concentrations of AscA are indicated in the figure. Data were fit with single exponential functions (red lines; $y = a+b.e^{-k_{obs}.t}$) to give observed rate constants ($k_{obs}$) at each AscA concentration. Each experiment was performed in triplicate. For the sake of clarity, only the trace of one replicate is shown for each condition. **(B)** Plot of pseudo-first order $k_{obs}$ as a function of AscA concentration. Error bars show ± s.d. (n = 3, independent experiments).

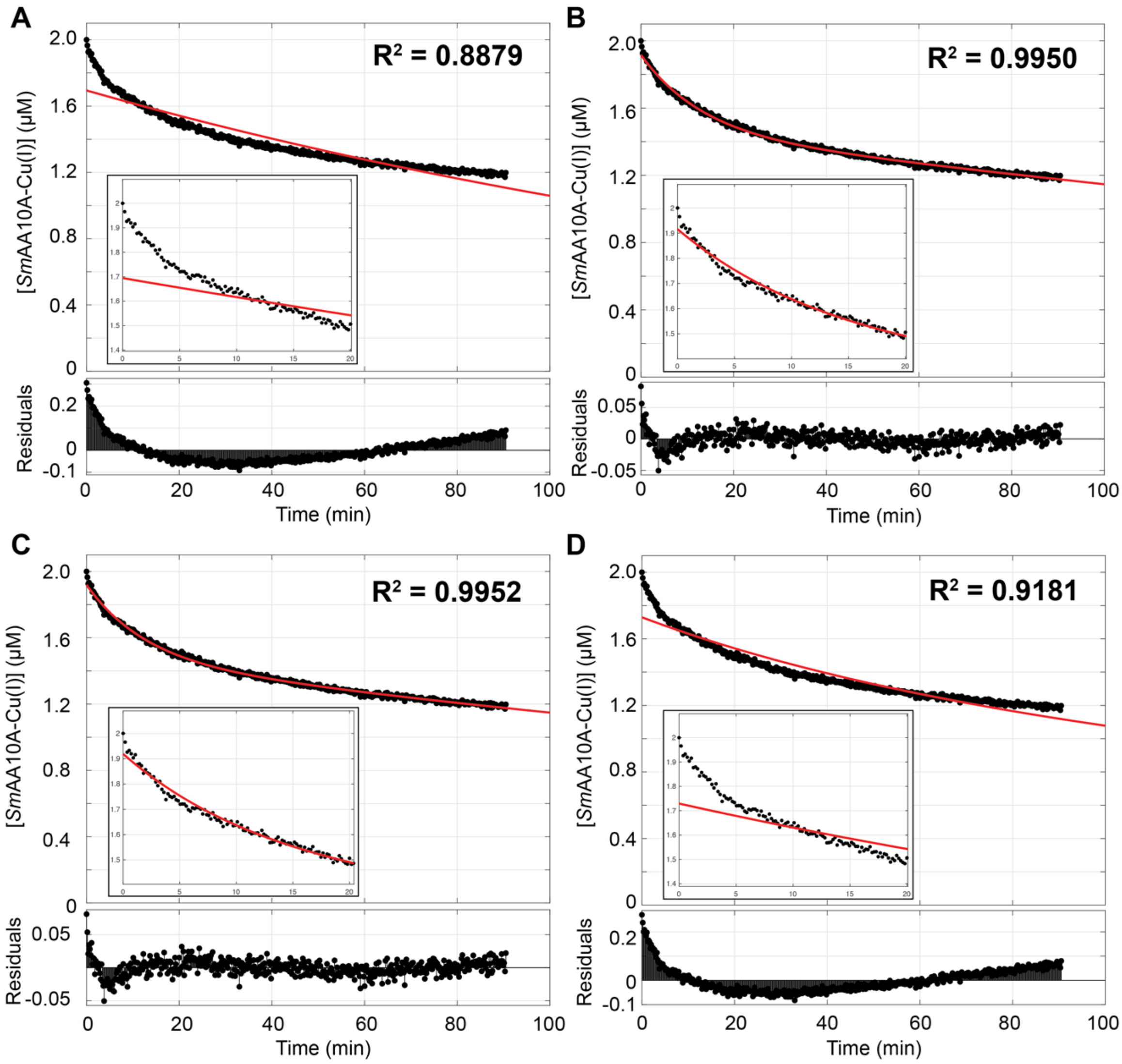


**Figure S13. Fitting of data for $O_2$-mediated re-oxidation of *Sm*AA10A-Cu(I), acquired at low $O_2$ concentration (47 µM), using several simple models.** The graphs show experimental data (black dots) and best fit (red line) for the re-oxidation of *Sm*AA10A-Cu(I) (2 µM) by $O_2$ (47 µM) over time. The inset shows a zoom-in view of the 20 first minutes. Reactions were carried out in sodium phosphate buffer (50 mM, pH 7.0) at 25 °C. See **Figure 5** in the main article for full set of experiments (i.e. for all O2 concentrations). Different models were tested to find how to better describe the data: **(A)** a single exponential function ($y = a.e^{-bt}$), **(B)** a single exponential function with a baseline correction factor ($y = a.e^{-bt} + c.t + d$), **(C)** a double exponential function ($y = a.e^{-bt} + c.e^{-dt}$) and **(D)** a function corresponding to a second order reaction ($1/y = 1/y_0 + a.t$). See **Figure S15** for alternative modelling of data. For the sake of clarity, only one replicate is displayed. The residuals are plotted below each corresponding graph. Fitting was performed with the Matlab "Curve Fitting" application.

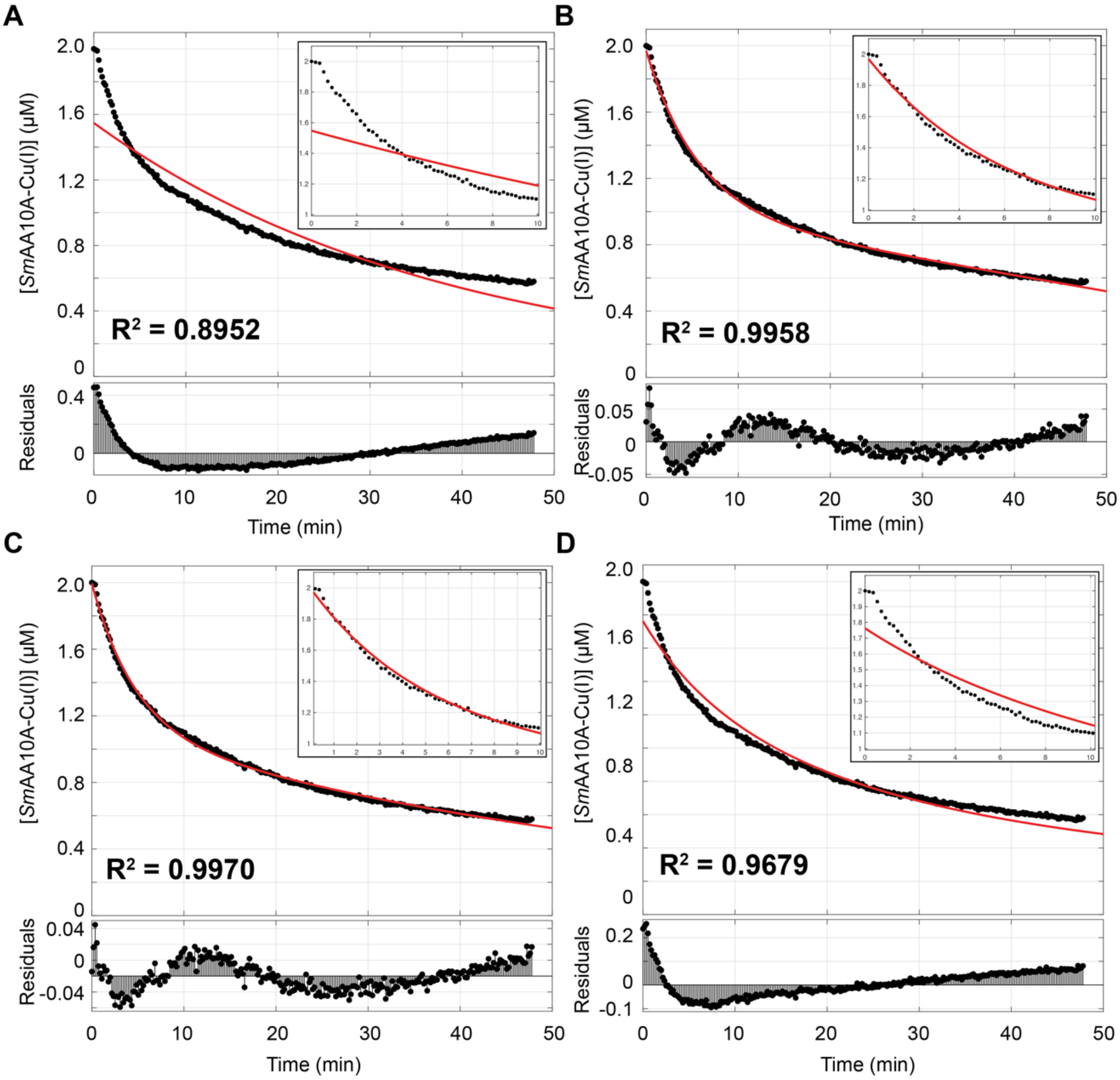


**Figure S14. Fitting of data for $O_2$-mediated re-oxidation of *Sm*AA10A-Cu(I) at high $O_2$ concentration (578 µM), using several simple models.** The graphs show experimental data (black circles) and best fit (red line) for re-oxidation of *Sm*AA10A-Cu(I) (2 µM) by $O_2$ (578 µM) over time. See **Figure S13** for details.

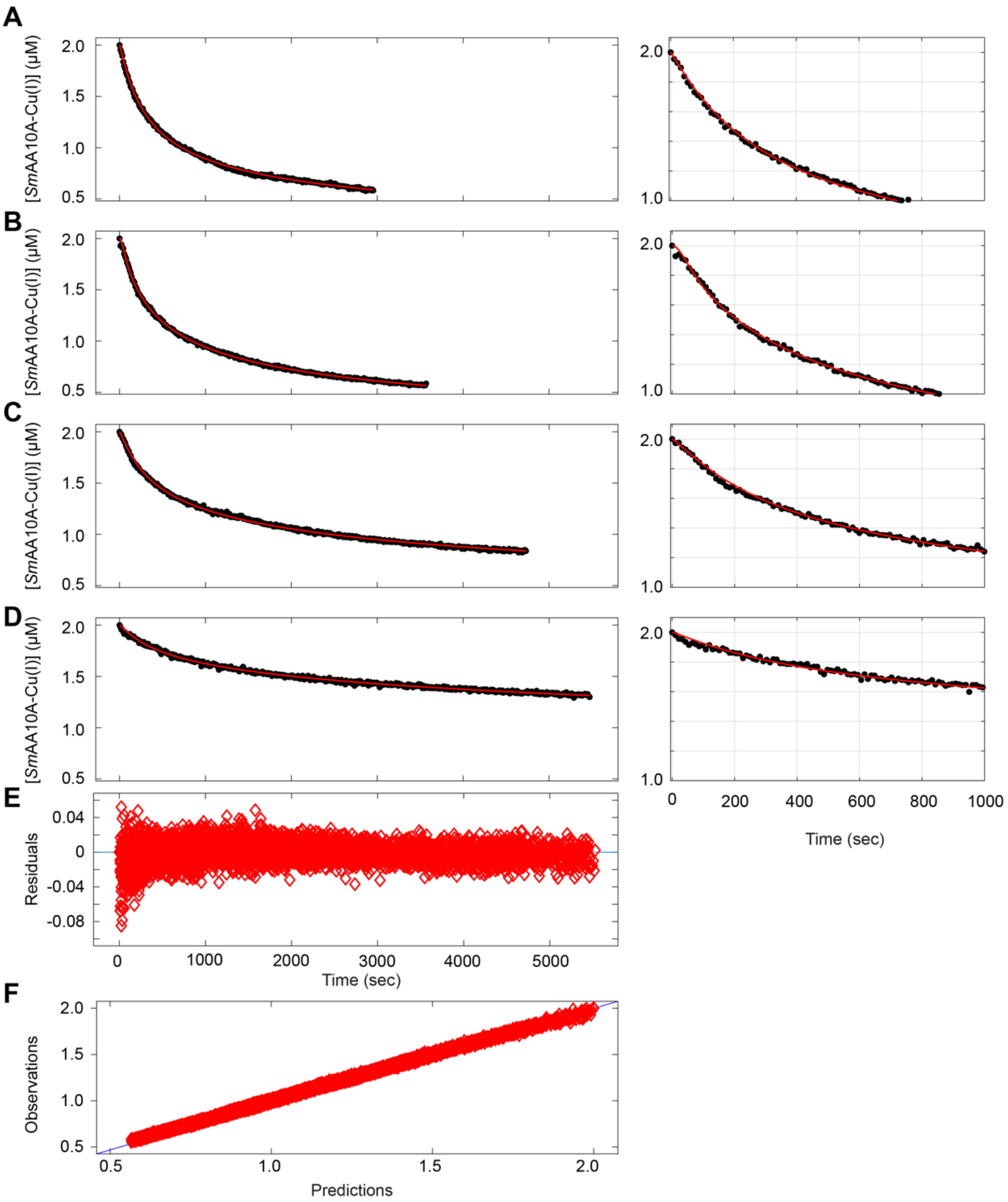


**Figure S15. Fitting data for $O_2$-mediated re-oxidation of *Sm*AA10A-Cu(I) to a model describing the dimer hypothesis. (A-D)** The graphs show experimental data (black circles) and best fit (red line) for re-oxidation of *Sm*AA10A-Cu(I) (2 µM) by $O_2$ over time using **(A)** 578 µM, **(B)** 422 µM, **(C)** 172 µM and **(D)** 47 µM of $O_2$. The fit corresponds to a system of differential equations describing the reaction scheme depicted in **Scheme S1**. Simulations were performed with Matlab-Simbiology application. All reactions were carried out in sodium phosphate buffer (50 mM, pH 7.0) at 25 ◦C. For the sake of clarity, only one replicate is displayed. A zoom-in view of the first 1000 sec is provided on the right-hand side. **(E)** Residuals and **(F)** plot of observed *vs* predicted data, taking into account the entire data set, i.e. all $O_2$ concentrations, in triplicates.

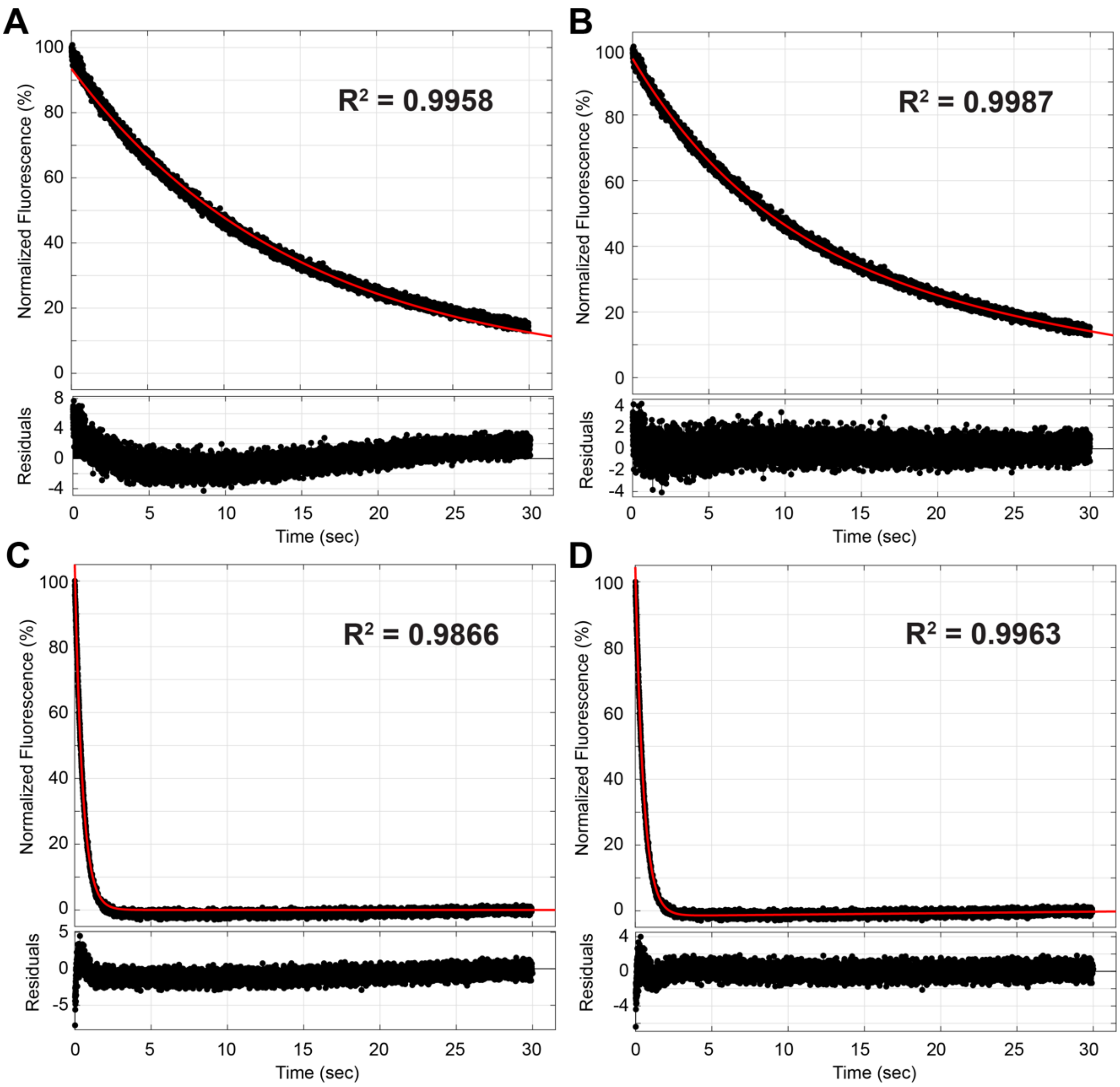


**Figure S16. Fitting data for $H_2O_2$-mediated re-oxidation of *Sm*AA10A-Cu(I) to a single exponential model.** The graphs show experimental data (black dots) and best fit (red line) for re-oxidation of *Sm*AA10A-Cu(I) (2 µM final) by $H_2O_2$ over time using (**A-B**) 5 µM (i.e. low) [$H_2O_2$] and (**C-D**) 400 µM (i.e. high) [$H_2O_2$]. In panels A and C show fitting to a single exponential function ($y = a.e^{-bt}$), whereas in panels B and D show fitting to a single exponential function with a baseline correction factor ($y = a.e^{-b.t} + c.t + d$). In agreement with usual practice related to stopped-flow data analysis, the latter procedure has been used for fitting all data presented in this study. For the sake of clarity, only one replicate is displayed. The residuals are plotted below each corresponding graph. Fitting was performed with the Matlab "Curve Fitting" application.

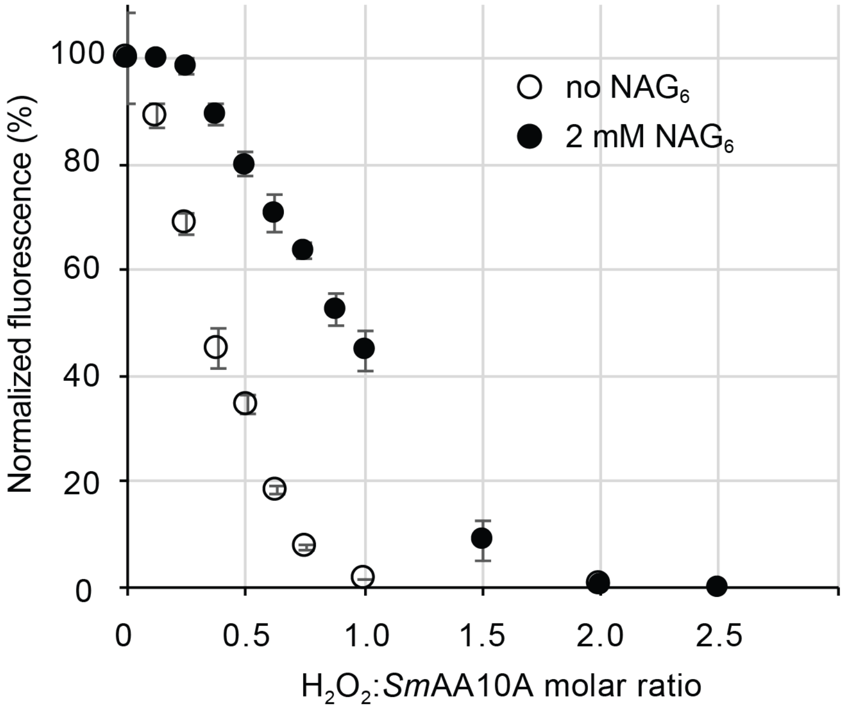


**Figure S17. Titration of *Sm*AA10A-Cu(I) with $H_2O_2$.** *Sm*AA10A-Cu(I) (2 µM final) was prepared anaerobically in a screw-cap quartz cuvette, with (black filled circles) or without (empty circles) $NAG_6$ (2 mM final) and mixed with $H_2O_2$ (0 to 5 µM final) before measuring the residual fluorescence (excitation 280 nm, emission 340 nm). The y-axis shows the relative variation in fluorescence, indicating the amount of *Sm*AA10A-Cu(I) (see material and methods). All reactions were performed at 25 °C in sodium phosphate buffer (50 mM, pH 7.0). Error bars show ± s.d. (n = 3, independent experiments).

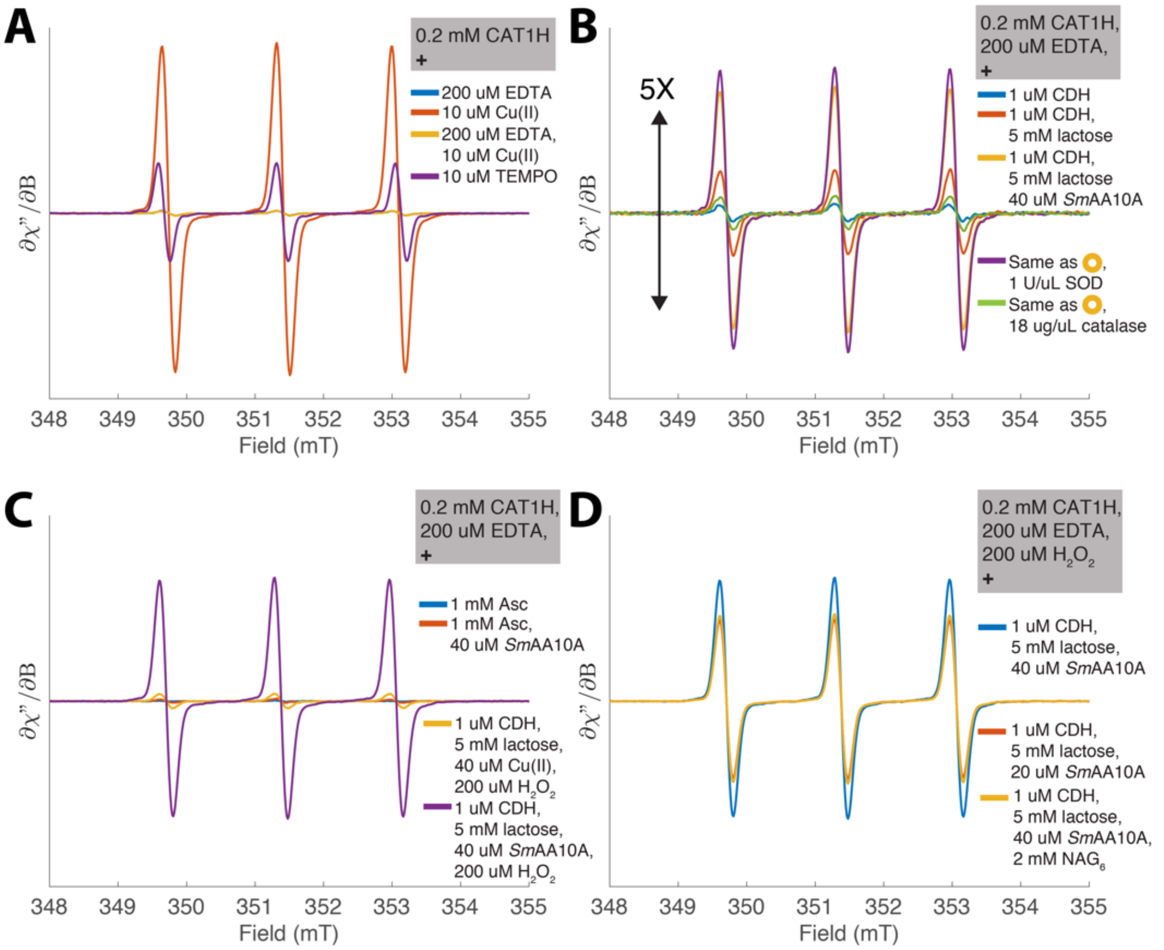


**Figure S18. Spin-trapping of reaction products generated when incubating *Sm*AA10A-Cu(I) with $H_2O_2$.** The spin probe cyclic hydroxylamine (CAT1H, 200 µM final) was used to monitor hydroxyl radical formation in different reactions (all performed in aerobic conditions), using either CDH/lactose or AscA (panel **C** only) as reductant (CDH, cellobiose dehydrogenase). Radical formation is reflected by the intensity of what we see in the graphs. Note that the ordinate axis of panel B is magnified 5x relative to the other panels. 10 µM TEMPO was used as the standard for all experiments ([A]purple) when double integrating the spectra to quantify radical formation. Panel **(A)** shows that the presence of EDTA (200 µM) is required to quench unspecific spin probe radical formation ([A]blue and [A]yellow, < 1 µM radical) arising from reaction of free Cu(II) with CAT1H ([A]red, 43 µM radical). Panel **(B)** shows that CDH alone contributes to minor amount of radical signal ([B]blue, < 1 µM radical) (similar to the background signal ([A]blue/[A]yellow) in panel **(A)**). After the CDH substrate (lactose, 5 mM final) was added, only a small amount of radical was formed ([B]red, 2.3 µM radical), perhaps from $H_2O_2$ generated by the CDH flavo-domain. In contrast, when *Sm*AA10A-Cu(II) was added to the reaction (40 µM final), a higher amount of radical was observed ([B]yellow, 6.6 µM radical), originating from reaction between $H_2O_2$ produced by CDH and *Sm*AA10A-Cu(I) (reduced by CDH).[61] Addition of SOD did not alter the amount of formed radical significantly ([B]purple, 7.4 µM), indicating that disproportioning superoxide in solution is not a source of $H_2O_2$. $H_2O_2$ is indeed thought to be directly produced by a two-electron reduction of $O_2$ at the CDH flavin domain.[62,63] Accordingly, addition of the $H_2O_2$-scavenging catalase led to almost complete suppression of radical formation ([B]green, 1.1 µM radical). Panel (**C**) shows that ascorbic acid in buffer ([C]blue) or ascorbic acid with *Sm*AA10A ([C]red) produced less than 0.5 µM radical. This control experiment is in agreement with the very low apparent production capacity of $H_2O_2$ by LPMO in presence of AscA [64] and indicates that, in contrast to the CDH/lactose system, the use of AscA is not appropriate to generate efficiently $H_2O_2$ *in situ* and/or that AscA scavenges the produced $H_2O_2$. Adding an excess of $H_2O_2$ (200 µM final) to a mixture of Cu(II) (40 µM final) mixed with CDH and lactose showed little formation of radical (2 µM, $^{C}$yellow). In contrast, replacing Cu(II) with the same amount of *Sm*AA10A-Cu(II) (i.e. 40 µM final) resulted in 32 µM radical ($^{C}$purple), indicating that *Sm*AA10A-Cu(II) reduced to *Sm*AA10A-Cu(I) by CDH reacts efficiently with $H_2O_2$. Panel **(D)** shows that in presence of excess $H_2O_2$ (200 µM final) about 35 % less radical was detected in the presence of 2 mM $NAG_6$ ($^{D}$yellow, 22 µM radical) than in absence of substrate ($^{D}$blue, 32 µM radical). Reducing the amount of *Sm*AA10A-Cu(II) by 50 % (i.e. 20 µM final) had a similar effect ($^{D}$red, 21 µM radical). All samples contained 0.2 mM CAT1H and were recorded exactly after 4 minutes incubation at room temperature. The CAT1H in buffer controls, prepared as first and last experiment were identical, verifying a stable stock solution of the spin probe. All reactions were performed in sodium phosphate buffer (50 mM pH 7.0).

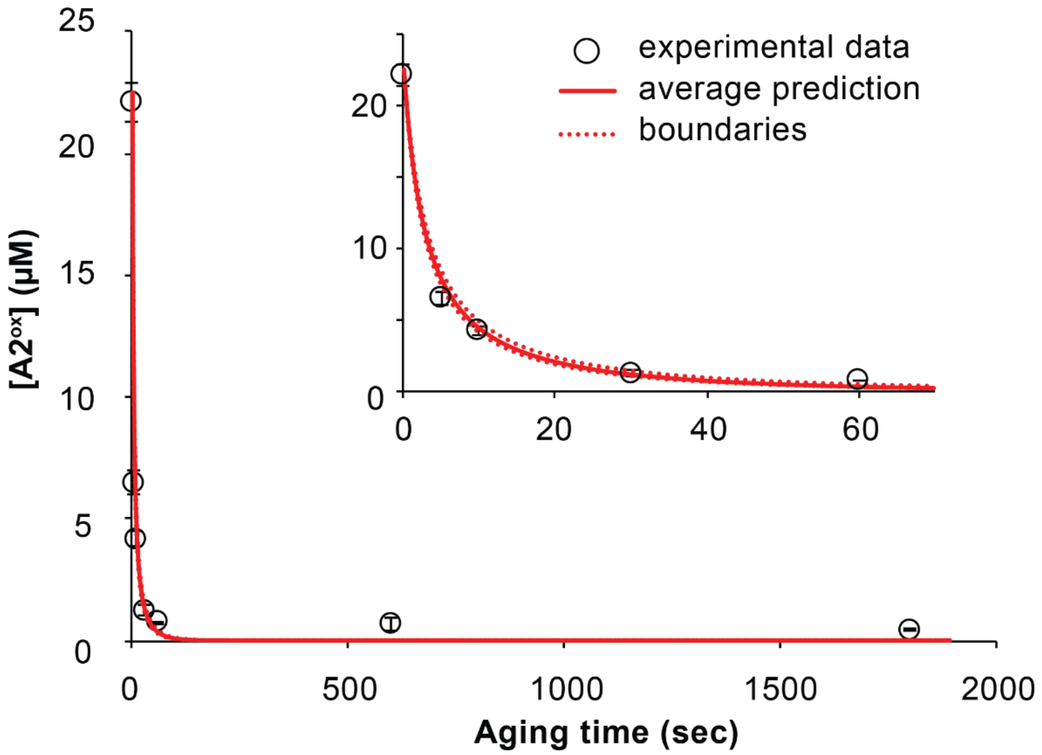


**Figure S19. Monitoring *Sm*AA10A-Cu(I) decay upon aging with $H_2O_2$ by measuring the residual oxidative activity.** *Sm*AA10A-Cu(I) (50 µM) was mixed with 0.9 equivalent of $H_2O_2$ (i.e. 45 µM) for varying amounts of time (5 s, 10 s, 30 s, 60 s, 10 min and 30 min) before addition of $NAG_6$ (2 mM final), under anaerobic conditions. After addition of the substrate, the reaction was incubated for 10 min, after which *Sm*GH20 was added to convert oxidized products to chitobionic acid ($A2^{ox}$). The graph shows the concentration of oxidized products detected in the final reaction mixture (expressed as $A2^{ox}$, black circles). The inset shows a zoom-in view of the first 60 seconds. The aging time "0 sec" was obtained by adding $H_2O_2$ after $NAG_6$. In negative control reactions, either $H_2O_2$ was replaced by anaerobic buffer or *Sm*AA10A-Cu(I) was replaced by stoichiometrically reduced $CuSO_4$. Both negative controls showed < 1 µM $A2^{ox}$ (not shown).

Using the second order rate constant of 6851 ± 597 $M^{-1}$ $s^{-1}$ for the reaction between *Sm*AA10A-Cu(I) and $H_2O_2$ as determined by stopped-flow kinetics (see **Table S3**), we can predict the amounts of residual reduced LPMO, i.e. *Sm*AA10A-Cu(I), and $H_2O_2$ available after a given aging time and, thus, the maximum amount of oxidized $NAG_6$ that can be expected (assuming that once $NAG_6$ is added, *Sm*AA10A-Cu(I) and $H_2O_2$ engage into $NAG_6$ oxidation). We show here that the predicted (red solid line) and measured (black circles) residual LPMO activity are in good agreement.

All reactions were performed at 25 °C, anaerobically, in sodium phosphate buffer (50 mM, pH 7.0). Error bars show ± s.d. (n = 3, independent experiments).

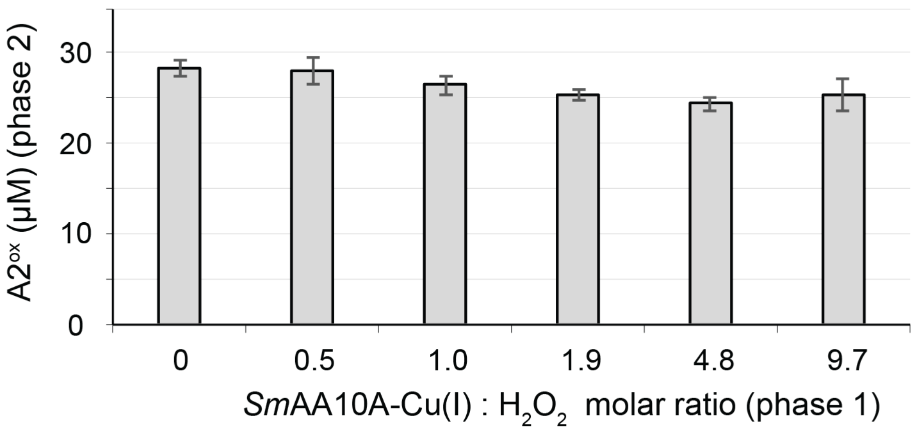


**Figure S20. Residual activity of *Sm*AA10A after treatment of *Sm*AA10A-Cu(I) wild-type with different $H_2O_2$ concentrations.** This experiment was composed of two phases: during phase 1, *Sm*AA10A-Cu(I) was isolated and mixed (10.3 µM final) with varying concentrations of $H_2O_2$ (0 to 100 µM final) during 20 min. Then, in phase 2, $(NAG)_6$ was added to the mixture (1.33 mM final), the $H_2O_2$ concentration was adjusted in to 200 µM final concentration, AscA was added as reductant (50 µM final) and the reaction was further incubated for 30 min. Both the pre-treatment reaction (phase 1) and activity test on $(NAG)_6$ (phase 2) were carried out in anaerobic conditions, in sodium phosphate buffer (50 mM, pH 7.0), at 25 °C. The reaction mixture was then hydrolyzed with the chitobiase *Sm*GH20 to yield chitobionic acid (A2$^{ox}$) as single final product, the quantity of which is shown in the graph for each pre-treatment condition. Error bars show ± s.d. (n = 3, independent experiments).

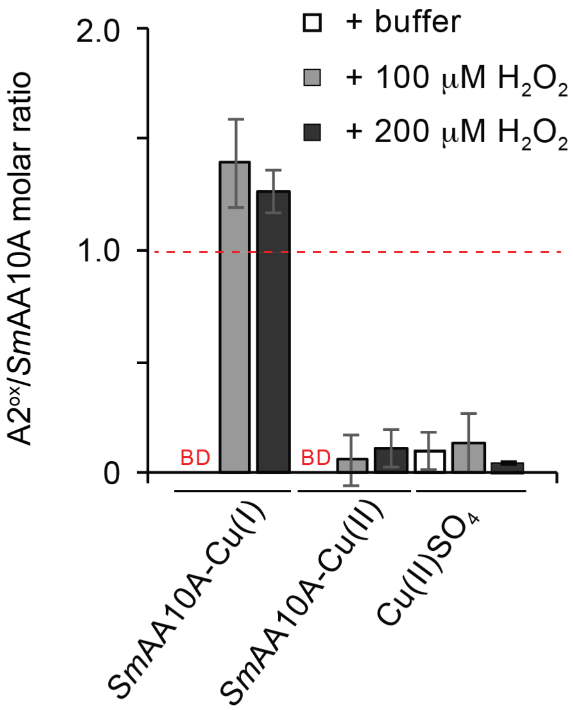


**Figure S21. $NAG_6$ is a suitable substrate for single turnover conditions.** *Sm*AA10A-Cu(II) (10.6 µM final) or *Sm*AA10A-Cu(I) (9.7 µM final) or $CuSO_4$ (10 µM final) was mixed in sodium phosphate buffer (50 mM, pH 7.0) with $NAG_6$ (2 mM final). After 15 min pre-incubation, $H_2O_2$ was added (0, 100 or 200 µM final) and the reaction was further incubated for 20 min before addition of the chitobiase *Sm*GH20 to yield chitobionic acid ($A2^{ox}$) as single final product. The graph shows the molar ratio of $A2^{ox}$/*Sm*AA10A (or $CuSO_4$) measured in each condition. Error bars show ± s.d. (n = 3, independent experiments). Abbreviations: BD, below detection limit.

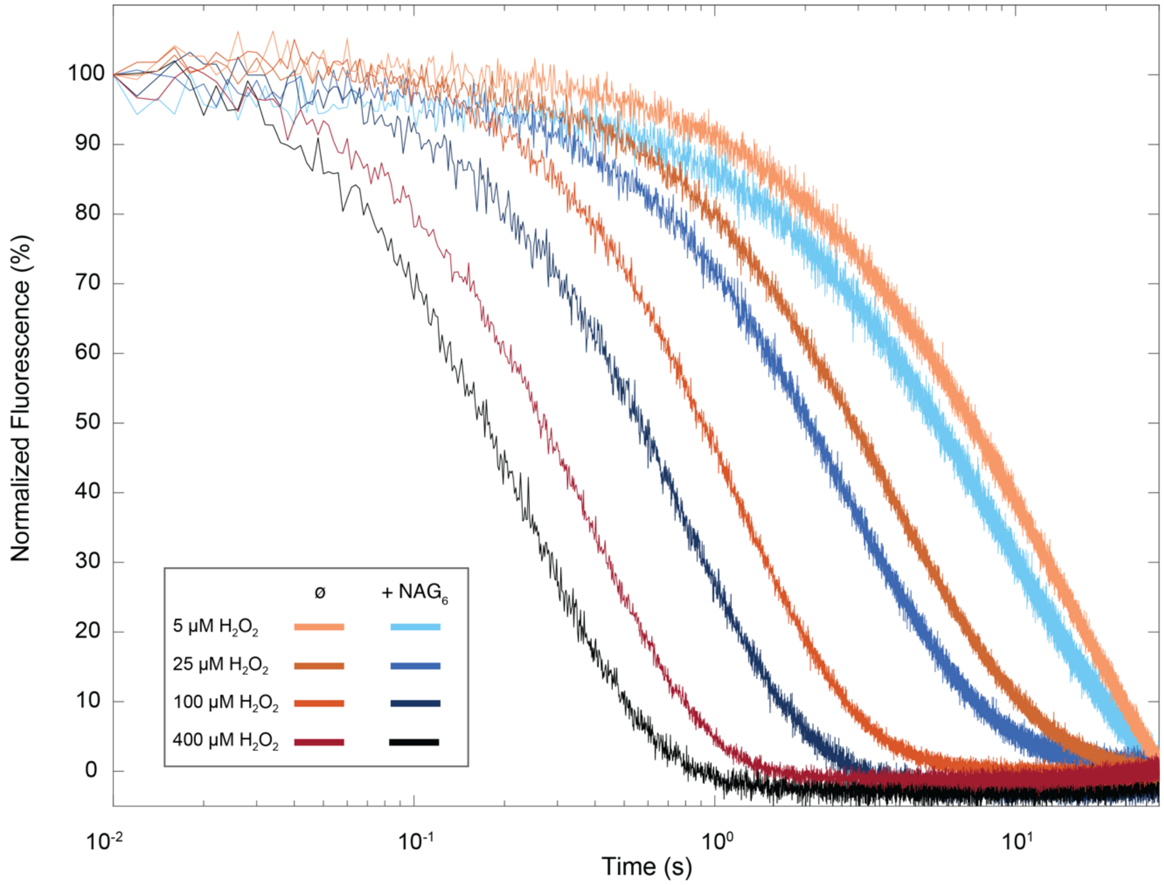


**Figure S22. Re-oxidation of *Sm*AA10A-Cu(I) by $H_2O_2$ in presence or absence of 0.5 mM $NAG_6$.** *Sm*AA10A-Cu(I) (final concentration 2 µM) was anaerobically mixed with buffer (lines with shades of orange) or with buffer containing $NAG_6$ (lines with shades of blue; final $NAG_6$ concentration 0.5 mM) and rapidly mixed in the stopped-flow spectrophotometer with various amounts of $H_2O_2$. The final $H_2O_2$ concentrations and corresponding color codes are indicated in the figure. Changes in fluorescence associated with re-oxidation of the Cu(I) were monitored as a function of time. All reactions were carried out in sodium phosphate buffer (50 mM, pH 7.0), at 25 °C. Data were fit with single exponential functions to give observed rate constants ($k_{obs}$) at each $H_2O_2$ concentration and plotted versus the latter in **Figure 6B** of the main manuscript. For the sake of clarity, the trace of only one replicate for each condition is shown in the figure. All experiments were performed at least in triplicate.

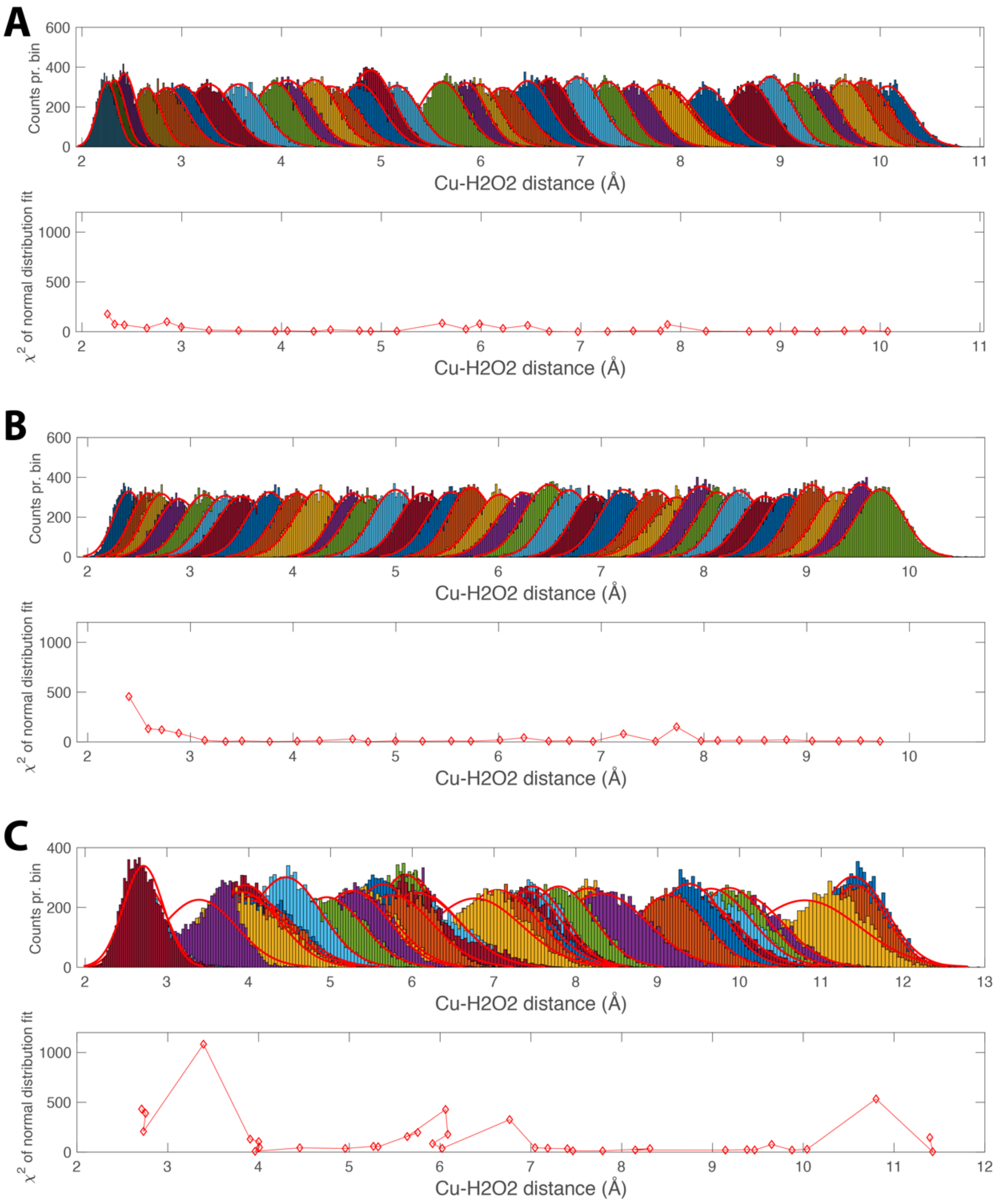


**Figure S23. Quality evaluation of umbrella-sampling data.** To provide properly sampled data for free energy integrators such as pyMBAR and WHAM, umbrella sampling should be carried out using an appropriate biasing potential (here 5.0 kcal/mol Å$^2$). Panels **A** and **B** show the histograms of the measured distances for the "pulling in" and "pushing out" experiments, respectively. All the histograms were fitted by the normal distribution function, and the corresponding $\chi^2$ values, indicating the quality of the fit, are shown directly below the histograms. In panel **A**, histograms in the range 1.75-2.5 Å were resampled using a higher biasing potential (10.0 kcal/mol Å$^2$), resulting in lower $\chi^2$ values at that interval. An example of using an inappropriate biasing potential (1.0 kcal/mol Å$^2$) is shown in panel **C**, where high $\chi^2$ values can be observed.

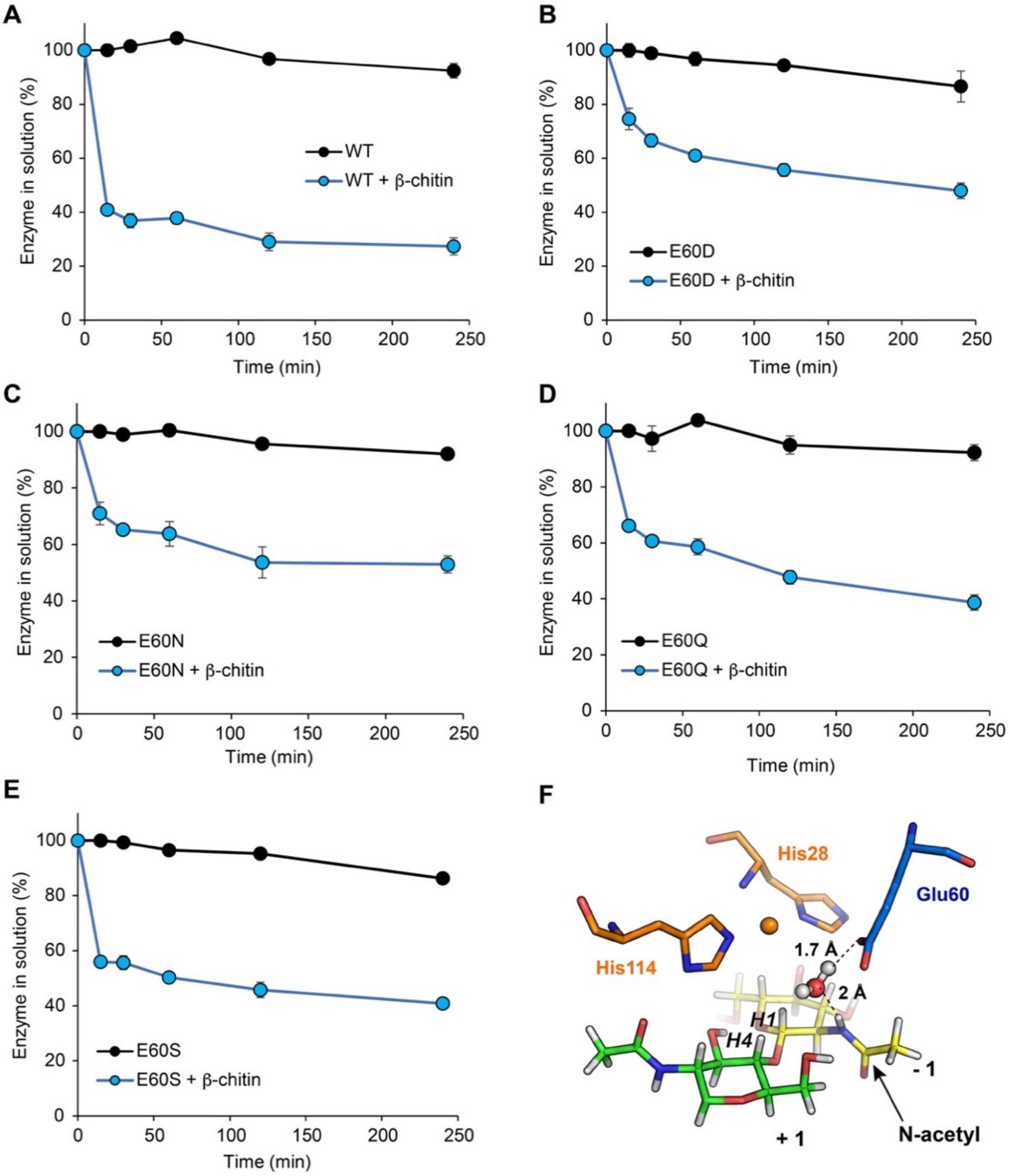


**Figure S24. Time-courses for binding of *Sm*AA10A wild-type and mutants to β-chitin.** For each experiment, a suspension of β-chitin in sodium phosphate buffer (50 mM, pH 7.0) was incubated at 40 °C in a thermomixer (1000 rpm). After 20 min of pre-incubation, **(A)** *Sm*AA10A-WT, **(B)** E60D, **(C)** E60N, **(D)** E60Q or **(E)** E60S (1 μM) were added to the substrate suspension as starting point of the binding time-courses (blue circles). The protein content in solution was determined after 15, 30, 60, 120 and 240 min of incubation. A control without β-chitin was carried out for each enzyme (black circles). Error bars show ± s.d. (n = 3, independent experiments). **(F)** View of *Sm*AA10A active site model showing a "water bridge" connecting the <correct name of the nitrogen atom> in the *N*-acetyl moiety of the -1 NAG unit and the Glu60 side chain.[13]

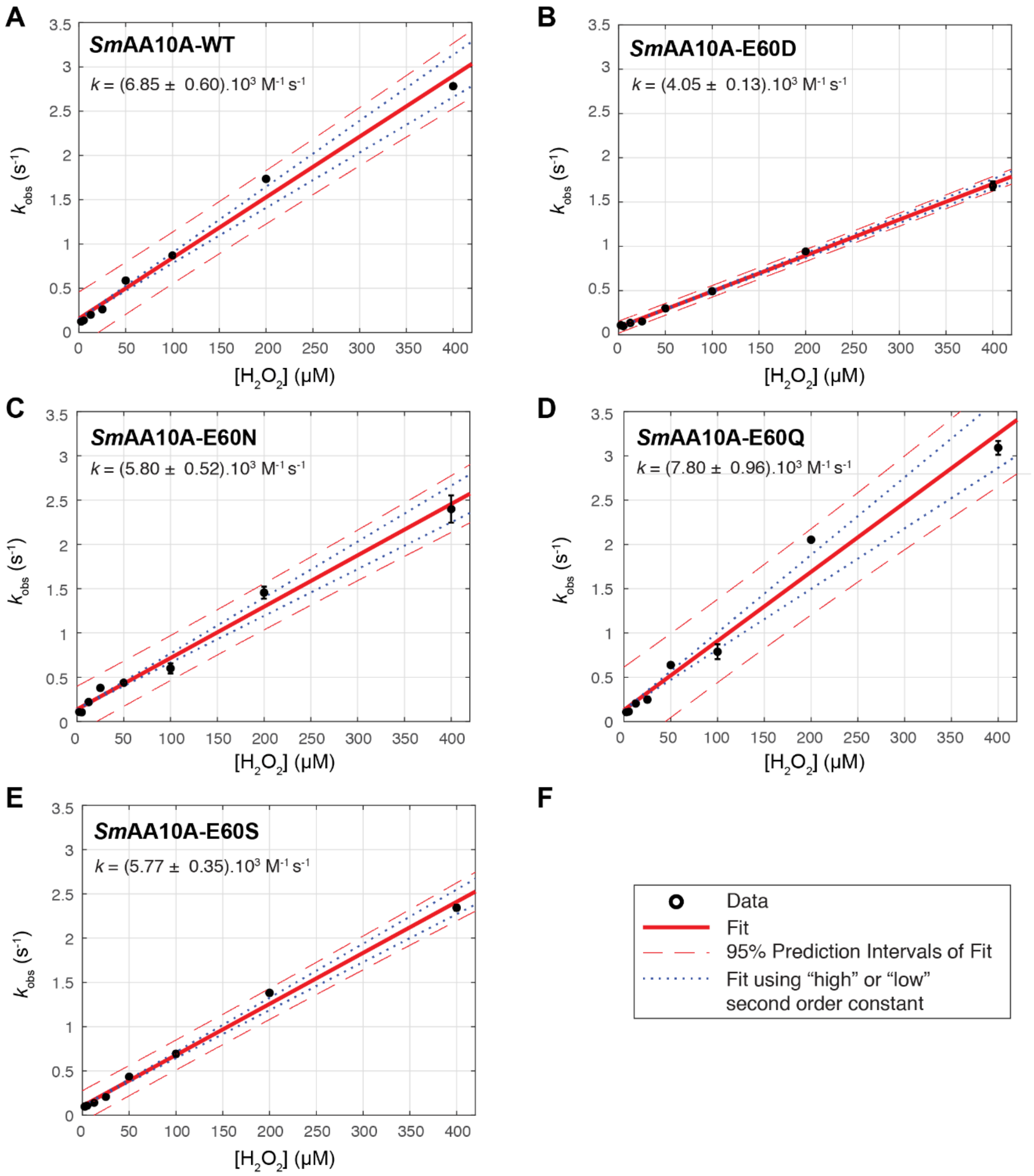


**Figure S25. Reactivity between $H_2O_2$ and *Sm*AA10A-Cu(I) wild-type and mutants.** Under anaerobic conditions, *Sm*AA10A-Cu(II) was reduced by addition of 20 eq. of AscA and desalted on a PD-10 column to yield *Sm*AA10A-Cu(I). *Sm*AA10A-Cu(I) was then rapidly mixed (2 µM final) with anaerobic buffer containing varying concentrations of $H_2O_2$ (2.5 to 400 µM final) and the subsequent change in protein fluorescence associated with Cu(I) re-oxidation was monitored as a function of time (See **Figure 5** in main article for examples of traces). All reactions were carried out in sodium phosphate buffer (50 mM, pH 7.0), at 25 °C. Time-course data were fit with single exponential functions with baseline correction factor (see **Figure S16**), yielding values for observed pseudo first-order rate constants ($k_{obs}$, $s^{-1}$). **Panels A to E** show the plot of $k_{obs}$ values as a function of the $H_2O_2$ concentration, for *Sm*AA10A-WT, E60D, E60N, E60Q and E60S, respectively. The plots were fit to a linear equation, yielding the second order rate constant *k* ($M^{-1}$ $s^{-1}$) (see **Table S3**). **Panel F** provides the legend code of panels A to E: average $k_{obs}$ values are shown as black dots, the best fit of the plot is shown as a red solid line and its associated 95% confidence interval as red dashed lines. The fits calculated using a slope of (*k*+error) (“high”) or (*k*-error) (“low”) are also shown as blue dotted lines. Error bars show ± s.d. (n ≥ 3, independent experiments).

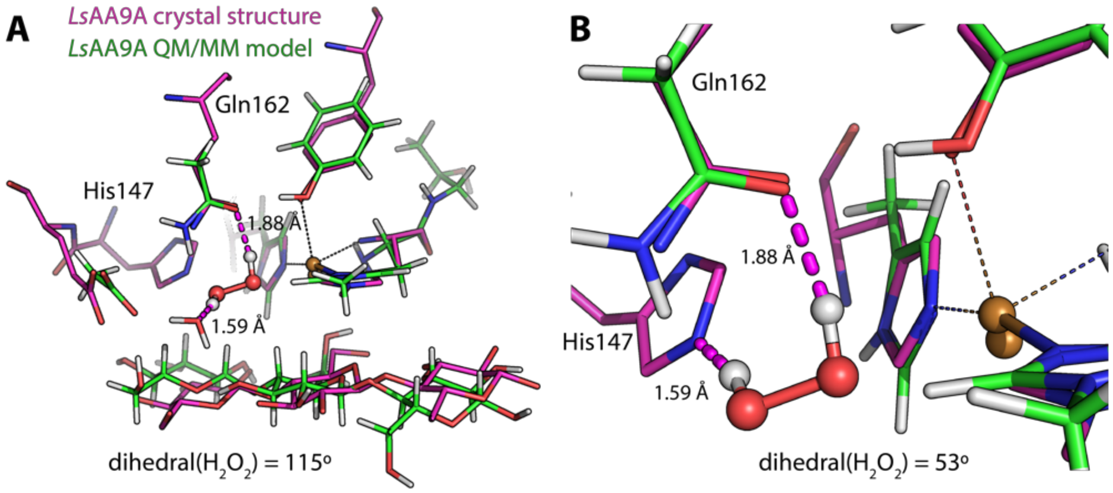


**Figure S26. Hydrogen bonding patterns involving $H_2O_2$ in the active site cavity of *Ls*AA9A. (A)** Superposition of the crystal structure of the fungal *L*sAA9A (magenta carbons), PDB ID 5ACF,[45] with a cellotriose molecule bound close to the active site and the QM/MM model, named $^1$RC1, generated by Wang et al. (green carbons),[41] where $H_2O_2$ is included in the active site. The $H_2O_2$ molecule assumes a relaxed conformation in the $^1$RC1 model, displaying a dihedral angle of 115°. In panel **B,** we have adjusted the dihedral angle of the $H_2O_2$ molecule in the $^1$RC1 model to 53°, which is the value that we observed for the analogous state **1** presented in this study. After altering the dihedral angle, the $H_2O_2$ is hydrogen bonded to His147 (HID state), a highly conserved residue not included in the QM/MM model. We suggest that His147 is involved in activating $H_2O_2$ for reaction with Cu(I), and that the Gln/His pair in AA9s provides the functionality provided by Glu60 in *Sm*AA10A. This would require His147 to be in the HID protonation state. We speculate that $H_2O_2$ reactivity with AA9s is controlled by the His147 protonation state, i.e. being predominantly protonated at the ε-position (HIE) while in solution and at the δ-position (HID) when interacting with substrate (see discussion in main article).

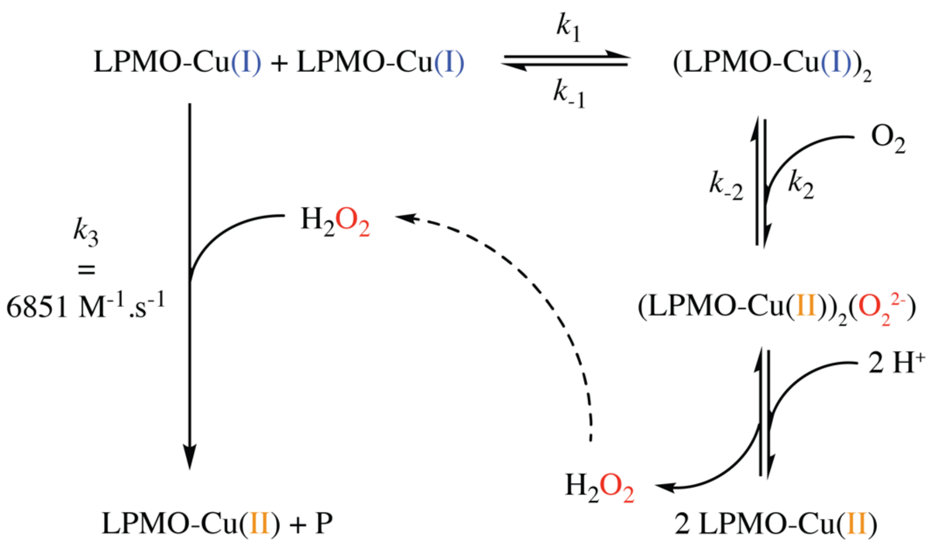


**Scheme S1. Proposed reaction scheme of *Sm*AA10A-Cu(I) with $O_2$ involving LPMO dimerization.** In the proposed reaction, $O_2$ reacts with the dimerized form of LPMO-Cu(I), allowing a two-electron reduction leading to a peroxo intermediate, which is rapidly protonated at pH 7.0 yielding $H_2O_2$. Such an $O_2$ reduction mechanism is analogous to what has been described for some organic dimers of mono-copper complexes,[65] and, potentially, to what is observed in binuclear coupled copper-enzymes such as tyrosinases.[66] The $H_2O_2$ produced *in situ* may react in a 1:1 stoichiometry with LPMO-Cu(I) (**Figure S16**), with a second order rate constant of 6851 $M^{-1}$ $s^{-1}$ (**Table S3**), yielding LPMO-Cu(II) and oxidative species denoted as P (e.g. $HO^{\bullet}$ + $HO^{-}$)(see **Figure S18**). **Figure S14** shows predictions of the time-course evolution of LPMO-Cu(I) using this proposed reaction scheme (modelled with the application SimBiology from Matlab). Nb. In the presence of substrate, the $H_2O_2$ produced *in situ* is preferentially used for hydroxylation of the substrate (Kuusk et al., 2018).

**Table S1. AMBER force field parameters for $H_2O_2$**

**Bond stretching**[a]

| | | $r_0$ (Å) | $k_r$ (kcal mol$^{-1}$ Å$^{-2}$) |
|---|---|---|---|
| O-O | | 1.448 | 322.0 |
| O-H | | 0.969 | 530.0 |

**Bond angle bending**[a]

| | | $\theta_0$ (°) | $k_\theta$ (kcal mol$^{-1}$ rad$^{-2}$) |
|---|---|---|---|
| H-O-O | | 60.55 | 101.5 |

**Dihedral angle parameters (H-O-O-H)**[a]

| | $k_\theta$ (kcal mol$^{-1}$) | Dihedral periodicity | Dihedral phase |
|---|---|---|---|
| 1 | 1.577 | 101.7 | -1 |
| 2 | -2.164 | 171.1 | -2 |
| 3 | -0.406 | 67.1 | 3 |

**Charges**[b]

| | |
|---|---|
| O | -0.36 |
| H | 0.36 |

[a] Calculated using Paramfit (AMBER) and Gaussian 09 (B3LYP/Def2-TZVP)

[b] Taken from Akiya et al.[67]

**Table S2. Population and spin parameters along the calculated reaction pathway**

| | Reaction states[b] | | | | | | | | |
|---|---|---|---|---|---|---|---|---|---|
| | **1** | **2** | **3** | **4** | **5** | **6** | **7** | **8** | **9** |
| *Atom*[a] | *Löwdin spin populations* | | | | | | | | |
| Cu | 0.00 | 0.30 | 0.60 | 0.64 | 0.61 | 0.50 | 0.60 | 0.39 | 0.00 |
| $O_{per\text{-}ox}$ | 0.00 | 0.00 | 0.02 | 0.64 | 1.11 | -0.32 | 0.13 | 0.14 | 0.00 |
| $O_{per\text{-}wat}$ | 0.00 | -0.35 | -0.78 | 0.43 | 0.01 | 0.00 | 0.00 | 0.00 | 0.00 |
| C1 | 0.00 | 0.00 | 0.00 | 0.00 | 0.00 | -0.14 | -0.61 | -0.43 | 0.00 |
| | *Unrestricted corresponding orbital (UCO) overlaps* | | | | | | | | |
| | 1 | 0.82 | 0.33 | 0 | 0 | 0.55 | 0.22 | 0.64 | 1 |
| | *Broken symmetry coupling constants (cm$^{-1}$)* | | | | | | | | |
| | 0 | 7940 | 1253 | 0 | 0 | 1990 | 420 | 3546 | 0 |

[a] Cu: copper; $O_{per\text{-}ox}$ : $H_2O_2$ derived O-atom that becomes the oxyl intermediate and that is incorporated in the product ; $O_{per\text{-}wat}$ : $H_2O_2$ derived O-atom that ends up as water ; C1 : carbon atom from which the H-atom is abstracted.

[b] Numbers correspond to the reaction states indicated in **Figure 3** of the main manuscript.

**Table S3. Second order rate constants of the re-oxidation by $H_2O_2$ of *Sm*AA10A-Cu(I) wild-type and mutants thereof [a,b]**

| *Sm*AA10A variant | $k_{H2O2}$[c] | | |
|---|---|---|---|
| | average ($M^{-1}$ $s^{-1}$) | error ($M^{-1}$ $s^{-1}$) | rel. $k_{H2O2}$ (%)[d] |
| WT | 6851 | 597 | 100 |
| E60D | 4047 | 133 | 59 |
| E60N | 5799 | 518 | 85 |
| E60Q | 7800 | 962 | 114 |
| E60S | 5768 | 347 | 84 |

[a] All reactions were carried out in sodium phosphate buffer (50 mM, pH 7.0), in anaerobic conditions, with 2 µM (final concentration) of the *Sm*AA10A-Cu(I) variant.
[b] Each experiment has been carried out at least in triplicate (n ≥ 3, independent experiments).
[c] The second order rate constant ($M^{-1}$ $s^{-1}$) corresponds to the slope of the plot of pseudo-first order rate constants ($s^{-1}$) as a function of $H_2O_2$ concentration (see **Figure S24**).
[d] Value of the second order rate constant relative to that of *Sm*AA10A-WT.

**Table S4. PCR primers used for site-directed mutagenesis of *Sm*AA10A**

| Introduced mutation | Primer sequence 5' to 3' * |
|---|---|
| E60D-forward | accgcagagcgtcga**t**ggcctgaaagg |
| E60D-reverse | cctttcaggcc**a**tcgacgctctgcggt |
| E60N-forward | gaaccgcagagcgtc**a**a**t**ggcctgaaaggcttc |
| E60N-reverse | gaagcctttcaggcc**a**t**t**gacgctctgcggttc |
| E60Q-forward | ccgcagagcgtc**c**agggcctgaaag |
| E60Q-reverse | ctttcaggccct**g**gacgctctgcgg |
| E60S-forward | gaaccgcagagcgtc**tc**gggcctgaaaggctt |
| E60S-reverse | aagcctttcaggccc**ga**gacgctctgcggttc |

* the target codon ("gag" in Sm*aa10a-wt*) is underlined and the introduced mutation(s) are shown as bold letters.

**Movie S1. QM/MM simulation of the proposed chitinolytic peroxygenase mechanism.**
The movie shows the reaction of $H_2O_2$ with *Sm*AA10A-Cu(I) in complex with β-chitin, as simulated by QM/MM. For the sake of clarity, the movie only includes one chitobiose unit from the chitin surface . The copper-coordinating histidines (His28 and His 114) and Glu60 (on the upper right hand corner) are shown are grey sticks. See **Figure S4-S5** for a complete description of the QM and MM regions, and **Figures 3, S8** and **Table S1** and **S2** for fine details of the simulation results.

In short, $H_2O_2$ reacts with Cu(I) inducing homolytic cleavage of the O-O bond and formation of Cu(II)-OH and $HO^•$. The resulting "precision-guided" $HO^•$ abstracts a hydrogen atom from Cu(II)-OH leading to release of a water molecule and formation of a $[CuO]^+$ core, which then catalyzes hydrogen atom abstraction from chitin. This yields a Cu(II)-OH species and a substrate radical (on the C1 carbon), which will merge via a re-bound mechanism,[36] yielding a C1-hydroxylated product. This orthoester is unstable and induces glycosidic bond cleavage (not shown here).[41,68]
To help the viewer, the movie has been paused (indicated by a "pause" sign in the upper left hand corner) at key steps of the mechanism.
The movie has been prepared with Pymol and Adobe After Effects, and represents 0.5 ps of simulation.

**Movie S2. Diffusion of $H_2O_2$ through the tunnel that leads from bulk solvent to the copper active site of *Sm*AA10A in complex with β-chitin.**
The movie shows the diffusion of $H_2O_2$ in the active site cavity in the*Sm*AA10A-Cu(I)-β-chitin complex, as simulated by Amber 16.
The enzyme active site is shown as beige sticks with surface, and sits on top of the upper layer of the chitin surface (only 3 out of 5 chitin chains are shown as blue sticks with surface). A side view (top panel) and a top view (lower panel) of the system are provided. The copper atom is shown as an orange sphere, and the gating residues, Glu60 and Asn185, are shown as blue and green sticks respectively. $H_2O_2$ is shown as red and white balls. See **Figures 8** and **S22** for more details.
The movie has been prepared with Pymol and Adobe After Effects, and represents 24 ns of simulation.

### 6. List of abbreviations

AscA: Ascorbic acid
AR : AmplexRed®
$A2^{ox}$ : Chitobionic acid or GlcNAcGlcNAc1A
GH: Glycoside hydrolases
HAA: Hydrogen atom abstraction
HRP: Horseradish peroxidase
LPMO: Lytic polysaccharide monooxygenases
*Ls*AA9A: LPMO9 from *Lentinus similis*
MD: Molecular dynamics
MM: Molecular mechanics
$NAG_6$ : Hexa-*N*-acetyl-chitohexaose
NEB : Nudged elastic band
QM : Quantum mechanics
*Sm*AA10A (or CBP21): LPMO10 from *Serratia marcescens*
*Sm*GH20: Chitobiase from *Serratia marcescens*
EPR: Electron paramagnetic resonance
SOMO: single occupied molecular orbital
HOMO: highest occupied molecular orbital
LUMO: Lowest unoccupied molecular orbital

### 8. QM/MM optimized xyz coordinates of states 1-9

Coordinates for the QM-region of QM/MM optimized states **1** to **9**. These structures were optimized using B3LYP, Def2-SVP for non-Cu atoms and Def2-TZPP for Cu.

**state1**
N 66.455002 49.006001 85.014999
C 66.655998 49.043999 86.461998
C 67.652000 47.966999 86.899002
O 67.500999 46.787998 86.594002
C 65.333000 48.861000 87.238998
C 64.487000 50.089001 87.212997
C 64.070999 50.848999 88.280998
N 64.099998 50.730000 86.045998
C 63.492001 51.859001 86.401001
N 63.455002 51.962002 87.745003
H 67.055000 50.034000 86.720001
H 64.803001 47.987000 86.842003
H 65.557999 48.631001 88.287003
H 64.202003 50.701000 89.347000
H 63.118000 52.616001 85.714996
H 63.000000 52.709000 88.264000
H 66.400002 48.042000 84.662003
H 67.227997 49.466999 84.532997
N 68.707001 48.453999 87.607002
H 68.703003 49.424999 87.928001
C 66.919998 55.962002 85.206001
C 66.663002 55.995998 83.696999
C 65.237999 55.566002 83.323997
O 64.323997 55.736000 84.195000
O 65.031998 55.087002 82.180000
H 66.818001 54.929001 85.564003
H 66.160004 56.566002 85.724998
H 66.797997 57.009998 83.277000
H 67.355003 55.335999 83.151001
C 63.498001 46.766998 85.001999
H 62.872002 46.758999 85.903000
H 64.155998 47.639999 85.011002
H 62.862000 46.841000 84.109001
C 65.938004 48.125000 79.294998
C 65.156998 48.433998 80.528000
C 65.535004 49.078999 81.684998
N 63.805000 48.205002 80.660004
C 63.401001 48.702000 81.847000
N 64.429001 49.243000 82.498001
H 66.969002 48.471001 79.455002
H 65.535004 48.693001 78.439003
H 63.221001 47.752998 79.933998
H 66.519997 49.443001 81.966003
H 62.362000 48.666000 82.163002
C 71.223000 52.724998 84.686996
C 69.713997 52.585999 84.681000
C 68.991997 52.438999 83.483002
C 69.001999 52.604000 85.891998
C 67.596001 52.362000 83.491997
C 67.607002 52.507000 85.903999
C 66.897003 52.405998 84.703003
H 71.691002 52.033001 83.973000
H 71.608002 52.493999 85.694000
H 69.529999 52.384998 82.531998
H 69.545998 52.709000 86.834999
H 67.042000 52.278000 82.553001
H 67.068001 52.530998 86.855003
H 65.807999 52.374001 84.696999
C 64.468002 50.020000 84.275002
C 60.987999 45.410999 77.806999
C 60.187000 50.123001 77.781998
C 60.695000 49.083000 78.792999
C 61.421001 46.797001 78.231003
N 60.464001 47.729000 78.322998
O 62.620998 47.051998 78.444000
C 60.028999 49.291000 80.158997
O 60.501999 48.320999 81.069000
C 60.292999 50.721001 80.635002
O 59.549999 51.014999 81.820000
C 59.870998 51.745998 79.556000
O 60.441002 51.439999 78.283997

C 60.265999 53.168999 79.928001
O 61.643002 53.221001 80.248001
H 59.098000 49.970001 77.639999
H 61.780998 49.200001 78.914001
H 58.937000 49.174000 80.035004
H 61.369999 50.848000 80.833000
H 58.764000 51.720001 79.484001
H 60.888000 45.363998 76.710999
H 60.019001 45.131001 78.245003
H 59.486000 47.460999 78.193001
H 61.764999 44.700001 78.112999
H 59.763000 48.091999 81.668999
H 60.015999 53.820999 79.070000
H 59.638000 53.473000 80.779999
H 61.925999 54.140999 80.432999
C 59.681000 55.507000 82.815002
C 60.151001 50.841999 83.057999
C 59.650002 51.914001 84.030998
C 59.147999 54.243999 83.450996
N 60.021999 53.221001 83.545998
O 57.974998 54.154999 83.819000
C 60.188999 51.680000 85.466003
O 59.595001 52.571999 86.377998
C 59.999001 50.233002 85.931999
O 60.890999 49.890999 87.019997
C 60.412998 49.251999 84.815002
O 59.792999 49.571999 83.583000
C 60.083000 47.801998 85.122002
O 58.686001 47.598000 85.123001
H 61.256001 50.898998 82.962997
H 58.551998 51.873001 84.055000
H 61.277000 51.841999 85.426003
H 58.943001 50.051998 86.197998
H 61.516998 49.334000 84.721001
H 60.764999 55.473000 82.643997
H 59.421001 56.363998 83.452003
H 61.009998 53.375999 83.329002
H 59.166000 55.646000 81.852997
H 60.132000 53.405998 86.376999
H 60.574001 47.182999 84.348999
H 60.541000 47.554001 86.095001
H 58.528000 46.681000 85.449997
O 63.924000 52.750999 83.224998
O 62.841999 53.555000 83.726997
H 63.292000 54.417999 83.967003
H 64.306000 53.341999 82.538002

**state2**

N 66.397003 49.070000 85.008003
C 66.651001 49.103001 86.455002
C 67.646004 48.007999 86.867996
O 67.486000 46.835999 86.542000
C 65.348000 48.939999 87.264000
C 64.500000 50.168999 87.227997
C 64.003998 50.897999 88.282997
N 64.163002 50.826000 86.056999
C 63.500999 51.930000 86.392998
N 63.389000 52.002998 87.732002
H 67.073997 50.087002 86.695999
H 64.806000 48.057999 86.899002
H 65.595001 48.730000 88.310997
H 64.077003 50.730999 89.351997
H 63.138000 52.681999 85.693001
H 62.888000 52.730000 88.239998
H 66.309998 48.103001 84.663002
H 67.181999 49.491001 84.507004
N 68.699997 48.473999 87.594002
H 68.696999 49.441002 87.930000
C 66.915001 55.983002 85.209000
C 66.679001 56.002998 83.698997
C 65.258003 55.569000 83.285004
O 64.336998 55.646000 84.163002
O 65.084000 55.191002 82.108002
H 66.792999 54.957001 85.581001
H 66.155998 56.598999 85.716003
H 66.822998 57.013000 83.272003
H 67.384003 55.344002 83.167999
C 63.495998 46.824001 85.028999
H 62.873001 46.811001 85.931000
H 64.164001 47.688000 85.054001
H 62.856998 46.917000 84.139999
C 65.934998 48.127998 79.260002
C 65.207001 48.466999 80.514999
C 65.641998 49.138000 81.638000
N 63.858002 48.265999 80.704002
C 63.502998 48.805000 81.886002
N 64.569000 49.344002 82.484001
H 66.968002 48.487999 79.365997
H 65.489998 48.672001 78.408997
H 63.242001 47.818001 80.000000
H 66.641998 49.499001 81.866997
H 62.476002 48.800999 82.240997
C 71.200996 52.723999 84.679001
C 69.693001 52.587002 84.609001
C 69.017998 52.451000 83.382004
C 68.929001 52.610001 85.789001
C 67.621002 52.410000 83.329002
C 67.531998 52.541000 85.740997
C 66.873001 52.470001 84.510002
H 71.698997 52.028999 83.988998
H 71.544998 52.502998 85.702003
H 69.594002 52.389999 82.454002
H 69.432999 52.706001 86.754997
H 67.103996 52.362000 82.366997
H 66.955002 52.577999 86.667999
H 65.783997 52.494999 84.459000
C 64.615997 50.241001 84.223999
C 61.103001 45.441002 77.820999
C 60.187000 50.127998 77.786003
C 60.706001 49.099998 78.805000
C 61.479000 46.835999 78.261002
N 60.495998 47.738998 78.348000
O 62.671001 47.123001 78.488998
C 60.049000 49.305000 80.174004
O 60.560001 48.353001 81.087997
C 60.291000 50.738998 80.643997
O 59.537998 51.027000 81.822998
C 59.875000 51.762001 79.559998
O 60.438000 51.446999 78.285004
C 60.297001 53.179001 79.927002

O 61.683998 53.209000 80.200996
H 59.099998 49.966999 77.646004
H 61.791000 49.233002 78.920998
H 58.959000 49.160999 80.066002
H 61.366001 50.872002 80.848000
H 58.768002 51.750000 79.494003
H 61.342999 45.320999 76.751999
H 60.040001 45.212002 77.973999
H 59.523998 47.451000 78.209999
H 61.734001 44.736000 78.376999
H 59.839001 48.131001 81.709999
H 60.028999 53.841000 79.082001
H 59.702000 53.485001 80.801003
H 61.977001 54.109001 80.456001
C 59.679001 55.512001 82.841003
C 60.132000 50.854000 83.065002
C 59.626999 51.918999 84.042000
C 59.132000 54.250000 83.467003
N 59.999001 53.223999 83.558998
O 57.956001 54.165001 83.824997
C 60.181999 51.681000 85.471001
O 59.603001 52.573002 86.390999
C 59.990002 50.233002 85.933998
O 60.882999 49.883999 87.021004
C 60.400002 49.254002 84.815002
O 59.779999 49.576000 83.584000
C 60.073002 47.803001 85.120003
O 58.678001 47.597000 85.127998
H 61.236000 50.919998 82.980003
H 58.528999 51.875000 84.073997
H 61.270000 51.840000 85.416000
H 58.935001 50.053001 86.203003
H 61.502998 49.335999 84.721001
H 60.768002 55.476002 82.703003
H 59.402000 56.372002 83.466003
H 61.000999 53.383999 83.388000
H 59.191002 55.646999 81.863998
H 60.143002 53.405998 86.384003
H 60.561001 47.185001 84.343002
H 60.536999 47.553001 86.091003
H 58.521000 46.679001 85.454002
O 63.618000 51.916000 83.484001
O 62.743000 53.442001 83.543999
H 63.449001 54.119999 83.713997
H 63.713001 51.937000 82.515999

**state3**
N 66.316002 49.077999 85.015999
C 66.635002 49.105999 86.459000
C 67.651001 48.013000 86.835999
O 67.491997 46.847000 86.492996
C 65.371002 48.924000 87.322998
C 64.521004 50.148998 87.311996
C 63.998001 50.870998 88.356003
N 64.213997 50.810001 86.139999
C 63.536999 51.910000 86.458000
N 63.393002 51.978001 87.792999
H 67.058998 50.094002 86.681000
H 64.820000 48.040001 86.975998
H 65.665001 48.715000 88.357002
H 64.035004 50.698002 89.426003
H 63.188999 52.647999 85.741997
H 62.866001 52.695000 88.290001
H 66.228996 48.110001 84.670998
H 67.089996 49.497002 84.492996
N 68.696999 48.476002 87.574997
H 68.689003 49.441002 87.917999
C 66.911003 55.986000 85.218002
C 66.676003 56.009998 83.709000
C 65.248001 55.609001 83.286003
O 64.317001 55.710999 84.154999
O 65.084000 55.245998 82.105003
H 66.786003 54.960999 85.589996
H 66.154999 56.605999 85.723999
H 66.841003 57.015999 83.281998
H 67.369003 55.338001 83.179001
C 63.481998 46.799000 85.060997
H 62.844002 46.764999 85.953003
H 64.147003 47.662998 85.117996
H 62.858002 46.909000 84.164001
C 65.921997 48.103001 79.272003
C 65.221001 48.425999 80.546997
C 65.689003 49.047001 81.684998
N 63.868999 48.261002 80.745003
C 63.542000 48.773998 81.945999
N 64.632004 49.255001 82.549004
H 66.952003 48.473999 79.357002
H 65.450996 48.647999 78.436996
H 63.236000 47.837002 80.042000
H 66.700996 49.380001 81.906998
H 62.516998 48.801998 82.307999
C 71.196999 52.723000 84.681000
C 69.688004 52.580002 84.612000
C 69.013000 52.429001 83.387001
C 68.921997 52.612000 85.791000
C 67.615997 52.382000 83.333000
C 67.526001 52.540001 85.744003
C 66.865997 52.455002 84.513000
H 71.695000 52.028000 83.991997
H 71.541000 52.502998 85.704002
H 69.589996 52.362999 82.459999
H 69.427002 52.722000 86.755997
H 67.099998 52.325001 82.371002
H 66.948997 52.591999 86.670998
H 65.776001 52.488998 84.459999
C 64.632004 50.223999 84.290001
C 61.106998 45.445000 77.825996
C 60.182999 50.122002 77.785004
C 60.712002 49.104000 78.808998
C 61.486000 46.837002 78.276001
N 60.502998 47.740002 78.358002
O 62.676998 47.123001 78.511002
C 60.064999 49.313999 80.181000
O 60.591999 48.369999 81.096001
C 60.299000 50.751999 80.642998
O 59.548000 51.039001 81.824997
C 59.868000 51.764000 79.555000
O 60.419998 51.444000 78.277000
C 60.305000 53.181000 79.900002
O 61.705002 53.206001 80.067001

H 59.097000 49.946999 77.643997
H 61.798000 49.241001 78.916000
H 58.974998 49.160000 80.083000
H 61.374001 50.896999 80.841003
H 58.759998 51.750999 79.501999
H 61.272999 45.355000 76.738998
H 60.056999 45.199001 78.040001
H 59.530998 47.455002 78.217003
H 61.779999 44.735001 78.322998
H 59.882000 48.160000 81.733002
H 59.973999 53.848999 79.081001
H 59.772999 53.479000 80.816002
H 62.007000 54.060001 80.440002
C 59.668999 55.514999 82.851997
C 60.145000 50.862999 83.066002
C 59.644001 51.924999 84.050003
C 59.137001 54.249001 83.485001
N 60.016998 53.235001 83.583000
O 57.960999 54.154999 83.844002
C 60.198002 51.674999 85.475998
O 59.624001 52.566002 86.403000
C 59.997002 50.229000 85.934998
O 60.888000 49.873001 87.025002
C 60.402000 49.252998 84.811996
O 59.779999 49.581001 83.584999
C 60.070999 47.801998 85.115997
O 58.675999 47.598999 85.127998
H 61.249001 50.921001 82.987000
H 58.546001 51.883999 84.084000
H 61.287998 51.823002 85.418999
H 58.941002 50.054001 86.203003
H 61.505001 49.334000 84.716003
H 60.757999 55.490002 82.714996
H 59.380001 56.375000 83.472000
H 61.015999 53.407001 83.372002
H 59.181000 55.634998 81.873001
H 60.159000 53.402000 86.385002
H 60.556000 47.183998 84.335999
H 60.536999 47.549999 86.085999
H 58.519001 46.680000 85.453003
O 63.478001 51.602001 83.677002
O 62.703999 53.673000 83.009003
H 63.286999 54.292000 83.528000
H 63.578999 51.780998 82.730003

**state4**
N 66.327003 49.099998 85.024002
C 66.646004 49.125999 86.468002
C 67.653000 48.023998 86.841003
O 67.486000 46.860001 86.497002
C 65.383003 48.956001 87.332001
C 64.535004 50.181999 87.311996
C 63.977001 50.886002 88.349998
N 64.237000 50.844002 86.136002
C 63.521999 51.922001 86.447998
N 63.356998 51.980999 87.780998
H 67.080002 50.110001 86.690002
H 64.828003 48.070999 86.991997
H 65.677002 48.750999 88.365997
H 63.997002 50.706001 89.418999
H 63.147999 52.652000 85.733002
H 62.801998 52.681000 88.273003
H 66.236000 48.131001 84.680000
H 67.101997 49.515999 84.500000
N 68.704002 48.480000 87.575996
H 68.702003 49.444000 87.921997
C 66.921997 55.984001 85.208000
C 66.686996 56.007999 83.698997
C 65.260002 55.591999 83.289001
O 64.347000 55.695999 84.177002
O 65.079002 55.213001 82.116997
H 66.797997 54.958000 85.579002
H 66.163002 56.599998 85.712997
H 66.841003 57.016998 83.273003
H 67.384003 55.341999 83.166000
C 63.478001 46.805000 85.084000
H 62.838001 46.768002 85.974998
H 64.147003 47.665001 85.151001
H 62.855000 46.925999 84.188004
C 65.947998 48.106998 79.269997
C 65.254997 48.425999 80.549004
C 65.737000 49.005001 81.702003
N 63.898998 48.285000 80.742996
C 63.584000 48.771999 81.956001
N 64.684998 49.208000 82.571999
H 66.982002 48.471001 79.350998
H 65.476997 48.658001 78.439003
H 63.258999 47.886002 80.029999
H 66.757004 49.305000 81.933998
H 62.561001 48.825001 82.318001
C 71.160004 52.734001 84.682999
C 69.651001 52.597000 84.574997
C 69.004997 52.451000 83.332001
C 68.856003 52.618000 85.735001
C 67.609001 52.397999 83.245003
C 67.460999 52.539001 85.653999
C 66.832001 52.457001 84.406998
H 71.671997 52.028000 84.014000
H 71.474998 52.521000 85.717003
H 69.600998 52.391998 82.417999
H 69.335999 52.723999 86.712997
H 67.113998 52.344002 82.272003
H 66.861000 52.580002 86.566002
H 65.742996 52.491001 84.330002
C 64.651001 50.207001 84.297997
C 61.136002 45.457001 77.885002
C 60.186001 50.134998 77.787003
C 60.716999 49.125000 78.817001
C 61.502998 46.862000 78.299004
N 60.514000 47.757000 78.377998
O 62.694000 47.160000 78.515999
C 60.068001 49.338001 80.186996
O 60.605999 48.405998 81.107002
C 60.283001 50.778999 80.643997
O 59.512001 51.056999 81.818001
C 59.859001 51.785000 79.547997
O 60.422001 51.458000 78.274002
C 60.284000 53.206001 79.889000
O 61.681000 53.244999 80.079002
H 59.099998 49.958000 77.649002

H 61.801998 49.268002 78.925003
H 58.980000 49.169998 80.086998
H 61.353001 50.933998 80.860001
H 58.750999 51.768002 79.485001
H 61.410000 45.306999 76.827003
H 60.068001 45.233002 78.012001
H 59.542999 47.462002 78.246002
H 61.749001 44.766998 78.478996
H 59.903999 48.205002 81.754997
H 59.959999 53.866001 79.060997
H 59.735001 53.506001 80.795998
H 61.974998 54.124001 80.398003
C 59.698002 55.527000 82.843002
C 60.105000 50.873001 83.059998
C 59.622002 51.939999 84.046997
C 59.146000 54.268002 83.469002
N 60.015999 53.245998 83.582001
O 57.962002 54.181000 83.805000
C 60.181999 51.682999 85.471001
O 59.622002 52.578999 86.403000
C 59.976002 50.237999 85.929001
O 60.870998 49.875999 87.015999
C 60.368999 49.264000 84.799004
O 59.723000 49.596001 83.584999
C 60.051998 47.810001 85.106003
O 58.658001 47.598000 85.133003
H 61.207001 50.910000 82.980003
H 58.523998 51.917999 84.085999
H 61.271999 51.824001 85.403999
H 58.921001 50.066002 86.203003
H 61.467999 49.352001 84.681999
H 60.787998 55.490002 82.716003
H 59.410999 56.391998 83.459000
H 61.028000 53.419998 83.460999
H 59.219002 55.653000 81.860001
H 60.168999 53.405998 86.389999
H 60.532001 47.195000 84.320999
H 60.529999 47.561001 86.070000
H 58.509998 46.676998 85.456001
O 63.296001 51.284000 83.542000
O 62.787998 53.506001 83.496002
H 63.403000 54.209999 83.831001
H 63.296001 52.308998 83.223999

**state5**
N 66.355003 49.034000 85.044998
C 66.634003 49.112000 86.495003
C 67.627998 48.012001 86.900002
O 67.457001 46.844002 86.571999
C 65.347000 48.995998 87.335999
C 64.516998 50.236000 87.240997
C 63.945000 51.002998 88.225998
N 64.271004 50.852001 86.028000
C 63.582001 51.967999 86.264000
N 63.373001 52.084000 87.587997
H 67.075996 50.097000 86.692001
H 64.785004 48.105000 87.024002
H 65.615997 48.833000 88.385002
H 63.930000 50.879002 89.303001
H 63.270000 52.695000 85.509003
H 62.859001 52.844002 88.032997
H 66.314003 48.051998 84.727997
H 67.123001 49.473000 84.528999
N 68.689003 48.474998 87.614998
H 68.692001 49.443001 87.950996
C 66.928001 55.952000 85.224998
C 66.638000 55.936001 83.723999
C 65.157997 55.609001 83.445999
O 64.317001 55.993999 84.296997
O 64.871002 55.000999 82.374001
H 66.818001 54.932999 85.619003
H 66.168999 56.571999 85.725998
H 66.822998 56.924999 83.263000
H 67.265999 55.209999 83.185997
C 63.477001 46.763000 85.100998
H 62.824001 46.721001 85.982002
H 64.151001 47.618000 85.188004
H 62.869999 46.893002 84.196999
C 65.931000 48.043999 79.223000
C 65.241997 48.334000 80.503998
C 65.732002 48.852001 81.681000
N 63.889999 48.172001 80.695999
C 63.580002 48.587002 81.933998
N 64.684998 48.998001 82.564003
H 66.959000 48.424999 79.295998
H 65.443001 48.584000 78.393997
H 63.250999 47.798000 79.967003
H 66.751999 49.140999 81.925003
H 62.560001 48.618999 82.307999
C 71.167999 52.734001 84.691002
C 69.665001 52.570999 84.546997
C 69.056999 52.384998 83.291000
C 68.835999 52.599998 85.682999
C 67.667000 52.300999 83.166000
C 67.445000 52.488998 85.565002
C 66.853996 52.368000 84.303001
H 71.706001 52.027000 84.044998
H 71.461998 52.541000 85.735001
H 69.681999 52.317001 82.396004
H 69.285004 52.737000 86.671997
H 67.199997 52.212002 82.181999
H 66.819000 52.535000 86.459000
H 65.767998 52.372002 84.190002
C 64.630997 49.988998 84.288002
C 61.098000 45.452999 77.842003
C 60.181999 50.129002 77.785004
C 60.705002 49.116001 78.817001
C 61.477001 46.851002 78.265999
N 60.495998 47.750999 78.370003
O 62.674000 47.138000 78.473999
C 60.042000 49.327000 80.181000
O 60.570000 48.396999 81.109001
C 60.243999 50.770000 80.641998
O 59.455002 51.048000 81.803001
C 59.837002 51.777000 79.540001
O 60.417000 51.452000 78.275002
C 60.256001 53.199001 79.887001
O 61.653999 53.243999 80.087997
H 59.097000 49.955002 77.641998
H 61.789001 49.254002 78.933998

H 58.955002 49.155998 80.070000
H 61.311001 50.924999 80.874001
H 58.730000 51.762001 79.461998
H 61.393002 45.301998 76.791000
H 60.026001 45.242001 77.948997
H 59.522999 47.462002 78.239998
H 61.691002 44.754002 78.446999
H 59.854000 48.180000 81.737999
H 59.938999 53.858002 79.056000
H 59.701000 53.497002 80.789001
H 61.937000 54.132000 80.389000
C 59.699001 55.537998 82.827003
C 60.043999 50.868000 83.050003
C 59.588001 51.952000 84.031998
C 59.137001 54.283001 83.449997
N 59.994999 53.250000 83.552002
O 57.955002 54.208000 83.797997
C 60.153000 51.695999 85.456001
O 59.595001 52.591000 86.389000
C 59.951000 50.250999 85.922997
O 60.862000 49.897999 87.000999
C 60.327000 49.273998 84.791000
O 59.650002 49.604000 83.589996
C 60.023998 47.820000 85.107002
O 58.632999 47.594002 85.139000
H 61.141998 50.882000 82.960999
H 58.491001 51.948002 84.083000
H 61.243999 51.840000 85.394997
H 58.900002 50.084999 86.212997
H 61.418999 49.368999 84.642998
H 60.792000 55.502998 82.728996
H 59.396000 56.404999 83.430000
H 61.001999 53.410000 83.378998
H 59.243999 55.657001 81.831001
H 60.132999 53.424999 86.365997
H 60.507999 47.205002 84.322998
H 60.507999 47.581001 86.070000
H 58.493999 46.672001 85.460999
O 63.146000 50.812000 83.515999
O 62.776001 53.450001 83.400002
H 63.436001 54.068001 83.011002
H 63.057999 52.544998 83.147003

**state6**
N 66.269997 49.036999 84.980003
C 66.607002 49.105999 86.417000
C 67.623001 48.015999 86.824997
O 67.470001 46.846001 86.494003
C 65.332001 48.980000 87.276001
C 64.499001 50.216999 87.172997
C 64.031998 51.028999 88.177002
N 64.183998 50.814999 85.962997
C 63.570999 51.970001 86.223000
N 63.465000 52.122002 87.555000
H 67.043999 50.095001 86.607002
H 64.779999 48.078999 86.974998
H 65.608002 48.834000 88.327003
H 64.092003 50.923000 89.253998
H 63.263000 52.702999 85.470001
H 63.036999 52.925999 88.009003
H 66.251999 48.061001 84.643997
H 66.989998 49.521000 84.436996
N 68.671997 48.481998 87.566002
H 68.662003 49.448002 87.905998
C 66.921997 55.958000 85.224998
C 66.638000 55.935001 83.722000
C 65.157997 55.609001 83.439003
O 64.315002 55.994999 84.288002
O 64.875999 55.000999 82.364998
H 66.805000 54.941002 85.624001
H 66.165001 56.584999 85.719002
H 66.828003 56.921001 83.257004
H 67.267998 55.205002 83.191002
C 63.473999 46.756001 85.075996
H 62.820999 46.712002 85.956001
H 64.147003 47.611000 85.164001
H 62.869999 46.886002 84.168999
C 65.989998 48.054001 79.250000
C 65.257004 48.341000 80.507004
C 65.681999 48.921001 81.678001
N 63.907001 48.113998 80.664001
C 63.535999 48.557999 81.873001
N 64.595001 49.048000 82.517998
H 67.026001 48.398998 79.372002
H 65.553001 48.623001 78.412003
H 63.297001 47.703999 79.931000
H 66.677002 49.271000 81.942001
H 62.506001 48.564999 82.210999
C 71.175003 52.731998 84.689003
C 69.668999 52.576000 84.573997
C 69.036003 52.397999 83.330002
C 68.864998 52.605000 85.726997
C 67.642998 52.319000 83.233002
C 67.472000 52.500000 85.637001
C 66.853996 52.383999 84.387001
H 71.697998 52.028999 84.027000
H 71.488998 52.528999 85.724998
H 69.640999 52.333000 82.421997
H 69.334999 52.735001 86.707001
H 67.156998 52.237000 82.258003
H 66.865997 52.547001 86.544998
H 65.765999 52.387001 84.301003
C 64.400002 49.976002 84.213997
C 61.047001 45.428001 77.800003
C 60.209999 50.126999 77.796997
C 60.737999 49.088001 78.804001
C 61.455002 46.813000 78.239998
N 60.492001 47.734001 78.348999
O 62.655998 47.076000 78.448997
C 60.127998 49.306999 80.196999
O 60.591999 48.306999 81.081001
C 60.466999 50.723999 80.665001
O 59.772999 51.050999 81.877998
C 59.988998 51.747002 79.598999
O 60.485001 51.438999 78.294998
C 60.391998 53.174999 79.931999
O 61.792999 53.261002 80.059998
H 59.118999 49.976002 77.675003
H 61.828999 49.198002 78.889999
H 59.028000 49.237000 80.111000

H 61.551998 50.827999 80.824997
H 58.880001 51.709999 79.596001
H 61.273998 45.307999 76.727997
H 59.980999 45.220001 77.962997
H 59.513000 47.465000 78.221001
H 61.669998 44.709000 78.346001
H 59.860001 48.090000 81.689003
H 60.012001 53.827999 79.123001
H 59.876999 53.456001 80.863998
H 62.062000 54.159000 80.351997
C 59.683998 55.488998 82.886002
C 60.283001 50.835999 83.135002
C 59.759998 51.908001 84.098999
C 59.186001 54.207001 83.512001
N 60.105000 53.226002 83.635002
O 58.004002 54.070999 83.841003
C 60.268002 51.659000 85.537003
O 59.658001 52.544998 86.443001
C 60.049999 50.209999 85.980003
O 60.935001 49.839001 87.067001
C 60.460999 49.220001 84.871002
O 59.912998 49.567001 83.607002
C 60.076000 47.780998 85.151001
O 58.675999 47.608002 85.120003
H 61.506001 50.851002 83.162003
H 58.660000 51.845001 84.101997
H 61.356998 51.806999 85.517998
H 58.991001 50.049000 86.247002
H 61.563000 49.278000 84.804001
H 60.772999 55.493999 82.749001
H 59.377998 56.340000 83.510002
H 61.110001 53.435001 83.492996
H 59.188999 55.604000 81.910004
H 60.173000 53.394001 86.421997
H 60.567001 47.154999 84.383003
H 60.505001 47.509998 86.133003
H 58.497002 46.693001 85.444000
O 62.844002 50.745998 83.404999
O 62.900002 53.395000 83.549004
H 63.521999 53.959000 83.039001
H 63.021999 52.450001 83.275002

**state7**

N 66.311996 49.035999 85.012001
C 66.629997 49.105999 86.455002
C 67.636002 48.011002 86.852997
O 67.475998 46.841999 86.521004
C 65.361000 48.979000 87.320999
C 64.528000 50.216999 87.246002
C 63.983002 50.979000 88.250000
N 64.253998 50.837002 86.042000
C 63.574001 51.952000 86.301003
N 63.398998 52.066002 87.629997
H 67.070000 50.092999 86.647003
H 64.799004 48.085999 87.014000
H 65.648003 48.811001 88.364998
H 63.993000 50.848000 89.325996
H 63.244999 52.683998 85.558998
H 62.896000 52.824001 88.086998
H 66.279999 48.057999 84.684998
H 67.056000 49.494999 84.481003
N 68.688004 48.476002 87.584000
H 68.681999 49.443001 87.924004
C 66.931000 55.952000 85.219002
C 66.647003 55.931999 83.716003
C 65.166000 55.601002 83.436996
O 64.324997 56.000000 84.279999
O 64.883003 54.978001 82.372002
H 66.817001 54.933998 85.615997
H 66.169998 56.574001 85.712997
H 66.831001 56.919998 83.254997
H 67.278999 55.206001 83.181999
C 63.480000 46.756001 85.103996
H 62.834999 46.706001 85.989998
H 64.152000 47.612000 85.190002
H 62.865002 46.887001 84.206001
C 65.926003 48.042000 79.237000
C 65.218002 48.311001 80.514000
C 65.667999 48.883999 81.681000
N 63.879002 48.063999 80.707001
C 63.541000 48.479000 81.939003
N 64.611000 48.983002 82.558998
H 66.950996 48.426998 79.328003
H 65.445000 48.588001 78.407997
H 63.259998 47.662998 79.975998
H 66.666000 49.244999 81.919998
H 62.519001 48.446999 82.305000
C 71.177002 52.730999 84.686996
C 69.670998 52.573002 84.571999
C 69.037003 52.403999 83.327003
C 68.867996 52.596001 85.725998
C 67.643997 52.325001 83.231003
C 67.474998 52.492001 85.636002
C 66.857002 52.382000 84.386002
H 71.700996 52.028000 84.024002
H 71.490997 52.526001 85.723000
H 69.641998 52.344002 82.417999
H 69.338997 52.722000 86.705002
H 67.156998 52.250999 82.255997
H 66.867996 52.533001 86.543999
H 65.767998 52.384998 84.301003
C 64.521004 49.992001 84.272003
C 61.069000 45.431999 77.794998
C 60.188000 50.118999 77.785004
C 60.712002 49.090000 78.801003
C 61.462002 46.820000 78.238998
N 60.490002 47.730999 78.339996
O 62.659000 47.094002 78.459000
C 60.057999 49.291000 80.174004
O 60.564999 48.327999 81.077003
C 60.289001 50.726002 80.648003
O 59.490002 51.014999 81.809998
C 59.873001 51.747002 79.558998
O 60.435001 51.435001 78.285004
C 60.285000 53.167999 79.916000
O 61.681999 53.222000 80.115997
H 59.101002 49.952999 77.646004
H 61.798000 49.216000 78.913002
H 58.966999 49.148998 80.061996
H 61.353001 50.875999 80.889999

H 58.764999 51.724998 79.497002
H 61.304001 45.313000 76.724998
H 60.004002 45.214001 77.952003
H 59.514999 47.452999 78.203003
H 61.695000 44.717999 78.346001
H 59.842999 48.104000 81.695999
H 59.960999 53.830002 79.088997
H 59.729000 53.458000 80.820000
H 61.959999 54.112999 80.411003
C 59.707001 55.526001 82.842003
C 59.976002 50.853001 83.068001
C 59.608002 51.942001 84.043999
C 59.145000 54.264999 83.454002
N 60.006001 53.231998 83.539001
O 57.963001 54.187000 83.804001
C 60.194000 51.695000 85.459999
O 59.625999 52.583000 86.391998
C 59.991001 50.250000 85.920998
O 60.883999 49.890999 87.009003
C 60.405998 49.266998 84.807999
O 59.797001 49.577999 83.553001
C 60.070999 47.819000 85.110001
O 58.674999 47.609001 85.111000
H 62.779999 50.667999 82.716003
H 58.506001 51.977001 84.150002
H 61.279999 51.841000 85.380997
H 58.935001 50.080002 86.195999
H 61.498001 49.373001 84.699997
H 60.799999 55.491001 82.749001
H 59.402000 56.386002 83.454002
H 61.015999 53.401001 83.382004
H 59.251999 55.653999 81.847000
H 60.165001 53.417999 86.380997
H 60.563000 47.196999 84.336998
H 60.529999 47.570999 86.083000
H 58.519001 46.692001 85.441002
O 62.945999 50.869999 83.646004
O 62.751999 53.513000 83.512001
H 63.446999 54.032001 83.052002
H 62.970001 52.553001 83.433998

**state8**

N 66.333000 49.028999 85.005997
C 66.623001 49.092999 86.445000
C 67.625999 48.004002 86.866997
O 67.470001 46.830002 86.550003
C 65.330002 48.963001 87.276001
C 64.502998 50.206001 87.199997
C 64.028999 50.986000 88.227997
N 64.190002 50.835999 86.005997
C 63.570999 51.977001 86.302002
N 63.458000 52.095001 87.639000
H 67.055000 50.081001 86.655998
H 64.776001 48.076000 86.941002
H 65.584000 48.783001 88.328003
H 64.089996 50.849998 89.302002
H 63.255001 52.730000 85.574997
H 63.012001 52.875000 88.116997
H 66.302002 48.056999 84.667000
H 67.067001 49.508999 84.480003
N 68.682999 48.476002 87.589996
H 68.678001 49.444000 87.922997
C 66.928001 55.951000 85.220001
C 66.643997 55.925999 83.717003
C 65.162003 55.599998 83.438004
O 64.320999 56.000999 84.280998
O 64.876999 54.978001 82.372002
H 66.815002 54.933998 85.619003
H 66.167999 56.575001 85.712997
H 66.833000 56.910999 83.250999
H 67.272003 55.194000 83.188004
C 63.491001 46.770000 85.068001
H 62.865002 46.743000 85.969002
H 64.163002 47.629002 85.110001
H 62.858002 46.875000 84.178001
C 65.945000 48.047001 79.246002
C 65.202003 48.320000 80.500999
C 65.607002 48.937000 81.663002
N 63.867001 48.035999 80.672997
C 63.491001 48.473999 81.887001
N 64.528999 49.029999 82.514000
H 66.973999 48.412998 79.371002
H 65.500000 48.605999 78.405998
H 63.269001 47.606998 79.943001
H 66.587997 49.334000 81.916000
H 62.465000 48.416000 82.238998
C 71.179001 52.731998 84.688004
C 69.672997 52.575001 84.574997
C 69.038002 52.404999 83.331001
C 68.872002 52.596001 85.730003
C 67.646004 52.321999 83.237999
C 67.478996 52.487000 85.642998
C 66.860001 52.374001 84.393997
H 71.702003 52.028999 84.026001
H 71.494003 52.528999 85.724998
H 69.641998 52.345001 82.421997
H 69.343002 52.721001 86.709000
H 67.157997 52.242001 82.263000
H 66.874001 52.520000 86.551003
H 65.771004 52.360001 84.313004
C 64.384003 50.028000 84.217003
C 61.023998 45.417999 77.769997
C 60.195999 50.118000 77.788002
C 60.714001 49.075001 78.792999
C 61.435001 46.800999 78.213997
N 60.473000 47.723999 78.324997
O 62.634998 47.062000 78.427002
C 60.078999 49.279999 80.178001
O 60.547001 48.283001 81.061996
C 60.389999 50.702000 80.649002
O 59.672001 51.039001 81.852997
C 59.936001 51.733002 79.583000
O 60.463001 51.430000 78.293999
C 60.339001 53.158001 79.939003
O 61.733002 53.221001 80.153000
H 59.106998 49.966999 77.654999
H 61.804001 49.182999 78.897003
H 58.981998 49.194000 80.074997
H 61.469002 50.820000 80.830002

H 58.827000 51.702000 79.554001
H 61.250000 45.299000 76.697998
H 59.958000 45.210999 77.933998
H 59.494999 47.456001 78.191002
H 61.646000 44.696999 78.314003
H 59.806000 48.040001 81.652000
H 60.018002 53.811001 79.103996
H 59.773998 53.451000 80.835999
H 62.007999 54.125000 80.417000
C 59.699001 55.507999 82.870003
C 60.102001 50.842999 83.111000
C 59.708000 51.928001 84.079002
C 59.174000 54.230000 83.481003
N 60.073002 53.229000 83.582001
O 57.991001 54.111000 83.813004
C 60.254002 51.683998 85.505997
O 59.645000 52.566002 86.417000
C 60.044998 50.235001 85.950996
O 60.919998 49.868000 87.043999
C 60.486000 49.248001 84.851997
O 59.967999 49.575001 83.554001
C 60.104000 47.806000 85.122002
O 58.702999 47.625999 85.087997
H 62.686001 50.667000 82.555000
H 58.603001 51.929001 84.152000
H 61.339001 51.842999 85.464996
H 58.984001 50.064999 86.208000
H 61.582001 49.328999 84.792999
H 60.791000 55.500000 82.763000
H 59.384998 56.359001 83.489998
H 61.084000 53.430000 83.459000
H 59.230000 55.632000 81.882004
H 60.169998 53.410000 86.413002
H 60.601002 47.185001 84.353996
H 60.533001 47.535999 86.101997
H 58.522999 46.716000 85.421997
O 62.650002 50.831001 83.508003
O 62.830002 53.479000 83.609001
H 63.497002 53.978001 83.087997
H 62.910000 52.512001 83.416000

**state9**

N 66.445000 48.963001 85.028000
C 66.651001 49.042999 86.474998
C 67.639000 47.963001 86.924004
O 67.486000 46.784000 86.619003
C 65.328003 48.907001 87.262001
C 64.492996 50.143002 87.178001
C 64.056999 50.952000 88.200996
N 64.129997 50.734001 85.977997
C 63.521000 51.882000 86.264999
N 63.459000 52.042000 87.601997
H 67.068001 50.033001 86.700996
H 64.788002 48.020000 86.907997
H 65.556000 48.722000 88.319000
H 64.162003 50.851002 89.275002
H 63.175999 52.606998 85.525002
H 63.021999 52.830002 88.073997
H 66.401001 47.988998 84.703003
H 67.216003 49.414001 84.532997
N 68.695000 48.450001 87.629997
H 68.691002 49.422001 87.949997
C 66.921997 55.970001 85.207001
C 66.669998 55.994999 83.697998
C 65.226997 55.615002 83.324997
O 64.321999 55.862000 84.192001
O 65.007004 55.119999 82.197998
H 66.809998 54.941002 85.572998
H 66.163002 56.583000 85.717003
H 66.834999 57.001999 83.269997
H 67.344002 55.312000 83.160004
C 63.495998 46.743999 85.056999
H 62.879002 46.724998 85.963997
H 64.160004 47.612999 85.074997
H 62.852001 46.832001 84.171997
C 65.888000 48.091999 79.260002
C 65.122002 48.375999 80.505997
C 65.496002 49.049000 81.650002
N 63.798000 48.051998 80.682999
C 63.403999 48.508999 81.888000
N 64.412003 49.129002 82.502998
H 66.907997 48.476002 79.394997
H 65.445999 48.637001 78.407997
H 63.215000 47.597000 79.956001
H 66.461998 49.483002 81.897003
H 62.389000 48.372002 82.250000
C 71.190002 52.731998 84.689003
C 69.680000 52.592999 84.632004
C 68.992996 52.456001 83.411003
C 68.927002 52.598000 85.819000
C 67.597000 52.382000 83.374001
C 67.530998 52.499001 85.787003
C 66.861000 52.411999 84.561996
H 71.680000 52.030998 84.000000
H 71.540001 52.512001 85.709999
H 69.558998 52.411999 82.476997
H 69.439003 52.695999 86.780998
H 67.071999 52.320999 82.417000
H 66.963997 52.514000 86.720001
H 65.769997 52.382999 84.531998
C 64.498001 49.914001 84.268997
C 61.013000 45.428001 77.744003
C 60.179001 50.122002 77.779999
C 60.688999 49.081001 78.790001
C 61.423000 46.806000 78.205002
N 60.459999 47.727001 78.318001
O 62.623001 47.066002 78.425003
C 60.029999 49.290001 80.157997
O 60.514999 48.324001 81.070999
C 60.307999 50.719002 80.625999
O 59.616001 51.005001 81.843002
C 59.865002 51.748001 79.560997
O 60.424000 51.439999 78.283997
C 60.273998 53.167999 79.931000
O 61.667999 53.227001 80.168999
H 59.091999 49.960999 77.634003
H 61.776001 49.196999 78.911003
H 58.938000 49.169998 80.049004
H 61.391998 50.853001 80.755997

H 58.757000 51.728001 79.500999
H 61.181000 45.344002 76.657997
H 59.958000 45.202999 77.954002
H 59.484001 47.456001 78.181000
H 61.669998 44.699001 78.236000
H 59.776001 48.085999 81.667000
H 59.972000 53.832001 79.098999
H 59.696999 53.456001 80.822998
H 61.948002 54.141998 80.382004
C 59.675999 55.509998 82.874001
C 60.238998 50.853001 83.087997
C 59.689999 51.916000 84.055000
C 59.153000 54.231998 83.487999
N 60.048000 53.230000 83.593002
O 57.971001 54.116001 83.820000
C 60.202000 51.676998 85.498001
O 59.588001 52.567001 86.399002
C 60.006001 50.229000 85.958000
O 60.897999 49.879002 87.043999
C 60.431999 49.240002 84.855003
O 59.879002 49.573002 83.585999
C 60.063999 47.797001 85.142998
O 58.667000 47.605000 85.112999
H 62.016998 50.292000 82.522003
H 58.595001 51.848000 84.044998
H 61.286999 51.841000 85.478996
H 58.949001 50.054001 86.225998
H 61.535000 49.293999 84.810997
H 60.762001 55.486000 82.718002
H 59.401001 56.360001 83.514000
H 61.055000 53.431000 83.487000
H 59.167999 55.653999 81.907997
H 60.126999 53.400002 86.404999
H 60.564999 47.172001 84.379997
H 60.499001 47.537998 86.124001
H 58.500000 46.689999 85.443001
O 61.617001 51.000000 83.045998
O 62.831001 53.644001 83.704002
H 63.264000 54.520000 83.872002
H 63.240002 53.339001 82.880997
